# Autophagy receptor NDP52 alters DNA conformation to modulate RNA Polymerase II transcription

**DOI:** 10.1101/2022.02.01.478690

**Authors:** Ália dos Santos, Daniel E. Rollins, Yukti Hari-Gupta, Hannah C. W. Reed, Mingxue Du, Sabrina Yong Zi Ru, Kseniia Pidlisna, Ane Stranger, Faeeza Lorgat, Ian Brown, Kevin Howland, Jesse Aaron, Lin Wang, Peter J. I. Ellis, Teng-Leong Chew, Marisa Martin-Fernandez, Alice L. B. Pyne, Christopher P. Toseland

## Abstract

NDP52 is an autophagy receptor involved in the recognition and degradation of invading pathogens and damaged organelles. Although NDP52 was first identified in the nucleus and is expressed throughout the cell, to date, there is no clear nuclear function for NDP52. Here, we use a multidisciplinary approach to characterise the biochemical properties and nuclear roles of NDP52. We found that NDP52 clusters with RNA Polymerase II (RNAPII) at transcription initiation sites and that its overexpression promotes the formation of additional transcriptional clusters. We also show that depletion of NDP52 impacts overall gene-expression levels in two model mammalian cells, and that transcription inhibition affects the spatial organisation and molecular dynamics of NDP52 in the nucleus. This directly links NDP52 to a role in RNAPII-dependent transcription. Furthermore, we also show that NDP52 binds specifically and with high affinity to double-stranded DNA (dsDNA) and that this interaction leads to changes in DNA structure *in vitro*. This, together with our proteomics data indicating enrichment for interactions with nucleosome remodelling proteins and DNA structure regulators, suggests a possible function for NDP52 in chromatin regulation. Overall, here we uncover novel nuclear roles for NDP52 in gene expression and DNA structure regulation.

## INTRODUCTION

NDP52/CALCOCO2, a 446 amino-acid autophagy receptor, was first identified in the nucleus, as a component of nuclear dots – multiprotein sub-compartments that respond to environmental stresses, such as viral infections^1^. However, later reports showed that the protein is distributed throughout the cell, with higher levels in the cytoplasm^2^. NDP52 has since been linked to cytoplasmic roles in autophagy and cell adhesion, where it is known to be required for pathogen-containing autophagosome maturation and membrane ruffle formation^3–5^; however, no nuclear function has been attributed to this protein.

NDP52 comprises a skeletal muscle and kidney enriched inositol phosphatase (SKIP) carboxyl homology (SKICH) domain, which facilitates membrane localisation^3^; a long coiled-coil (CC) region that includes a predicted leucine-zipper (LZ) domain, and two zinc finger domains at the C-terminal - ZF1 and ZF2 (Fig.1A)^1^. The CC region of NDP52 has been identified as a potential homodimerisation domain for the protein^6^. At the C-terminal, ZF1 has been characterised as an unconventional dynamic zinc finger, whilst ZF2 is a canonical C_2_H_2_-type zinc finger^7^. The C-terminal domains of NDP52 are responsible for interactions with ubiquitin, which allows binding to ubiquitylated pathogens, as well as interactions with actin-based motor Myosin VI (MVI)^3,7–9^. In the cytoplasm, interactions between NDP52 and MVI allow autophagosome maturation ^5^. However, there is little information available regarding the biochemical and structural properties of the full-length protein, which limits our understanding of its functions.

**Figure 1:**
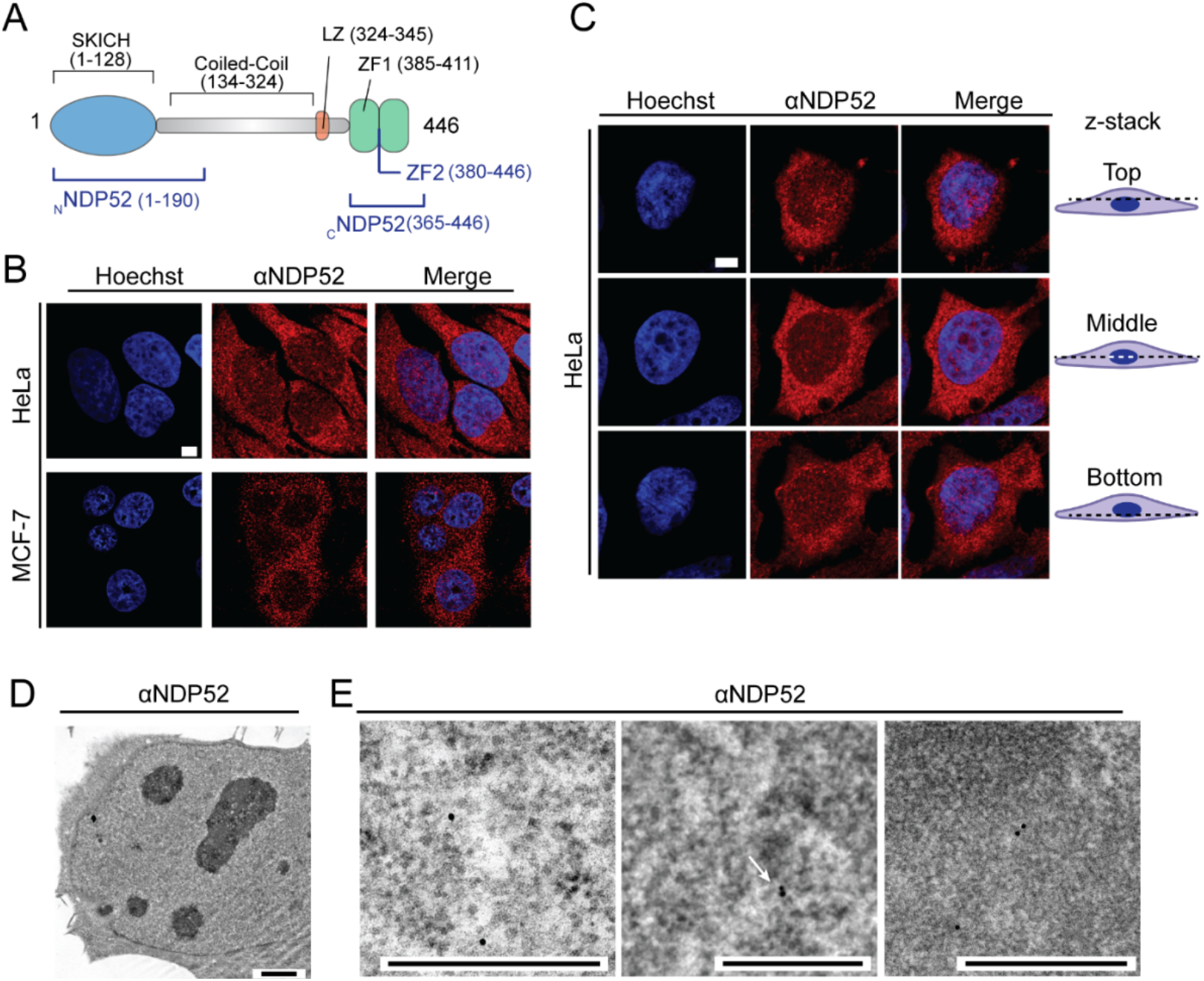
NDP52 is distributed throughout the nucleus. **(A)** Diagram of NDP52 displaying protein domains and key features, as well as recombinant constructs used in this study (in blue). **(B)** Confocal imaging of HeLa and MCF-7 cells labelled by immunofluorescence against NDP52 (red), with DNA staining shown in blue. Scale bar = 5μm. **(C)** Confocal imaging of the intracellular distribution of NDP52 at different z points in HeLa. Scale bar = 5μm. **(D)** Electron microscopy of HeLa cells following immune-gold labelling of NDP52. Scale bar = 2μm. Approximate thickness = 80 nm **(E)** Electron microscopy detail images of immune-gold labelled NDP52 (black dots) in the nucleus. Scale bar = 500 nm.

NDP52 is a member of the Calcium-binding and coiled-coil domain containing (CALCOCO) family. Other members are TAX1BP1 and CoCoA. NDP52 shares high sequence homology with both TAX1BP1 and CoCoA, and all three proteins have similar domain structure. Interestingly, whilst TAX1BP1 is also a known autophagy receptor^3,10^, CoCoA is a well-characterised transcription coactivator^11^. Recently, CoCoA has also been linked to roles in autophagy, further highlighting potential functional similarities between these proteins^12,13^. Furthermore, a recent study by Fili *et al*. has revealed that the interaction between NDP52 and MVI enhances RNA Polymerase II (RNAPII) transcriptional activity *in vitro*^14–16^.

Here, we explore the spatial organisation of NDP52 in the nucleus, as well as its dynamic behaviour; and assess how perturbation of this protein affects gene expression in cells. We found that NDP52 forms clusters in the nucleus at RNAPII transcription initiation sites and that knockdown of NDP52 significantly affects gene expression in both HeLa and MCF-7 cells. Furthermore, our biochemical analysis shows that NDP52 binds to double-stranded DNA with high affinity and Atomic Force Microscopy (AFM) suggests this results in changes to DNA shape and structure *in vitro*. We have also explored the nuclear interactome of NDP52, which shows enrichment for proteins involved in DNA structure and nucleosome regulation. We suggest that NDP52 has a regulatory role in RNAPII-dependent transcription, and that this arises both from direct interactions with chromatin as well as from protein-protein interactions with chromatin regulators and transcription factors. Overall, this highlights a wider role of NDP52 across the cell and it remains to be determined if there are links between its cytoplasmic and nuclear functions.

## RESULTS

### Nuclear organisation and dynamics of NDP52

To attribute a nuclear function to NDP52, we first assessed its nuclear localisation in two example mammalian cancer cell lines. Immunofluorescence staining of NDP52 in both HeLa and MCF-7 cells shows that NDP52 is distributed throughout the cytoplasm and nucleus (Fig.1B). Confocal imaging of different focal planes also shows distribution of the protein throughout the organelle (Fig.1C). To further confirm this, we also used electron microscopy with gold-immunolabelling of endogenous NDP52. Imaging of negative stained HeLa sections (c.a. 70 nm thickness) (Fig.1D) clearly shows NDP52 particles in nuclear regions, which can be observed in zoomed-in sections in Fig.1E. The presence of NDP52 in the nucleus is consistent with previous reports ^1,2,15^.

Within the nuclear region, NDP52 appears to cluster into small punctate regions of high fluorescence intensity (Fig.2A). To explore the nuclear organisation of NDP52, we used Stochastic Optical Reconstruction Microscopy (STORM) in both HeLa and MCF-7 cells (Fig.2B).

**Figure 2:**
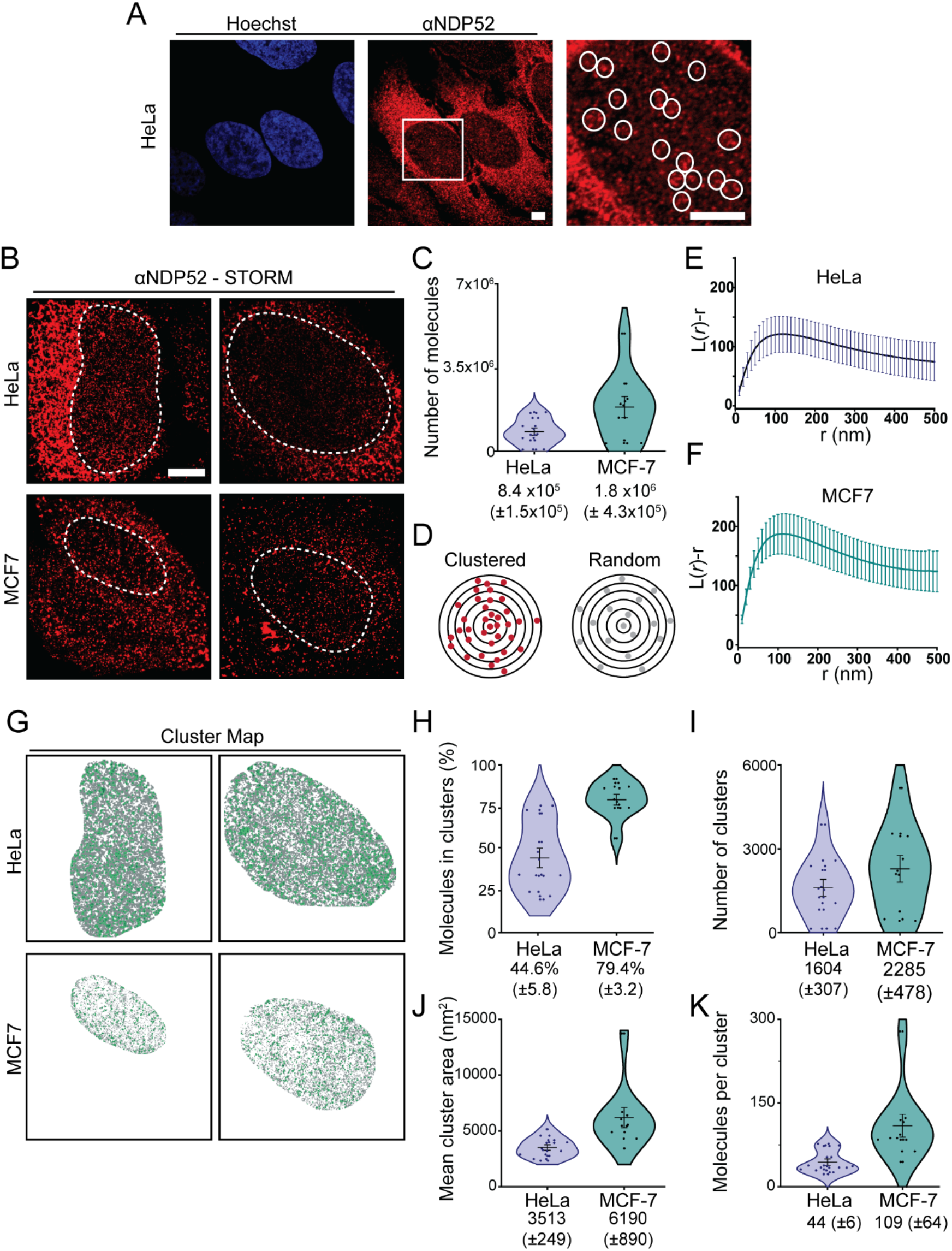
Spatial organisation of NDP52 in the nucleus. **(A)** Confocal image of NDP52 in HeLa cells showing detail of dense nuclear staining in white circles (zoomed-in right panel). Scale bar = 5μm **(B)** Example STORM images of NDP52 in HeLa and MCF-7 cells. Dotted lines represent selected regions of interest (ROIs) for the nucleus. These regions were used for cluster analysis. Scale bar = 5μm **(C)** Linearized Ripley’s K Function, L(r)-r (where r is the radius), calculated for selected ROIs from STORM images in HeLa and MCF-7. A value of zero in this plot signifies molecules are randomly distributed, whilst positive values indicate molecular clustering. Mean values are plotted ± SEM. n= 12 (HeLa), n= 10 (MCF-7) **(D)** Diagram depicting molecular clustering and random distribution. **(E)** Cluster maps generated for ROIs displayed in (C), using parameters specified in Methods. Clustered molecules are shown in green. **(F-J)** Cluster analysis of NDP52 in the nucleus of HeLa and MCF-7 showing: **(F)** total number of molecules; **(G)** percentage of molecules in clusters; **(H)** number of clusters in ROIs; **(I)** mean cluster area in nm^2^ and **(J)** number of molecules per clusters. Mean ± SEM values are shown. n= 12 (HeLa), n= 10 (MCF-7).

STORM allows us to visualise with high spatial precision and quantify individual molecules of NDP52 within a specified region of interest (ROI) (Fig. 2C), in this case the nuclear region. Furthermore, in-depth analysis of STORM data can also provide information regarding the clustering behaviour of the protein (Fig.2D). Protein clustering is often related to the molecular function of a protein and is particularly important in the enhancement of enzymatic processes such as transcription, DNA repair and DNA replication ^17–21^. Hence, as we investigate a nuclear function for NDP52, it is important to study its spatial organisation and how this might be linked to its nuclear role. To determine if NDP52 is randomly distributed or forms clusters (Fig.2D), we used a linearised Ripley’s K function ^22^. In both cell lines we observe a high probability for nuclear clustering of NDP52, as the Ripley’s K function deviates from zero towards positive values (Fig.2E and F). To further understand the organisation of NDP52 clusters in the nucleus, we used ClusDoC software ^22^. We defined NDP52 clusters, as regions where a minimum of 5 neighbouring molecules are spaced at a distance smaller than the mean value of localisation precision from STORM acquisition (described Methods). This allowed us to generate cluster maps for selected nuclear regions (Fig.2G) and determine that approximately 45% (±6) and 79% (±3) of NDP52 molecules are clustered in HeLa and MCF-7 cells, respectively.

This corresponds to an average of 1604 (±307) and 2285 (±478) clusters per cell in HeLa and MCF-7 cells, respectively, with an average size of 3513 nm^2^ (±249) and 6190 nm^2^ (±890), and 44 (±6) and 109 (±64) molecules of NDP52 per cluster (Fig.2H-K). In both HeLa and MCF-7 cells, we observe large cell-to-cell variation for clustering data. Although STORM provides detailed information regarding the spatial organisation of molecules, it is also a low-throughput technique. Cell variability could be a result of cells not being synchronised; however, due to this limited throughput, it is also not possible to identify multiple subpopulations within the data.

To assess how the spatial distribution of NDP52 relates to its molecular dynamics in the nucleus, we transiently expressed Halo-NDP52 in HeLa cells. This allowed us to use Fluorescence Recovery After Photobleaching (FRAP) to assess how dynamic NDP52 molecules are in the nucleus of live-cells (Fig.3A). Our data show that NDP52 has a recovery half-time of 7.5 s (±0.8) (Fig.3B and Supplementary Fig.1A) and a mobile fraction of 0.65 (±0.02) (Fig.3C). This is in agreement with molecular clustering data showing that approximately 45% of NDP52 molecules are clustered, and would therefore be expected to be less dynamic. To obtain more detailed information on the dynamic behaviour of nuclear NDP52, we used aberration-corrected Multi-Focal Microscopy (acMFM). This technique allows us to simultaneously track single-molecules across nine focal planes in live-cells, covering 4μm in the *z* axis and 20 x 20 μm in *xy* (Fig.3D). We obtained 3D trajectories of Halo-NDP52 molecules in the nucleus of HeLa cells (Fig.3E). Analysis of the different trajectories can then provide information on how confined or diffuse molecules are (Fig. 3F). By measuring the Mean Squared Displacement (MSD) of each molecule (Supplementary Fig.1B), we were also able to calculate diffusion coefficients (Fig.3G) and anomalous diffusion constants (α) (Supplementary Fig.1C) for each track. These were then plotted as the average diffusion coefficient or average α per cell (Fig.3H and I). Under normal conditions, NDP52 nuclear diffusion is relatively slow (*D* =0.24 (±0.008) μm^2^/s) and molecules are mostly confined, displaying an α value lower than 1 (α = 0.7 (±0.006)) (Fig.3H and I). Furthermore, acMFM data show that approximately 55% (±0.84) of nuclear NDP52 molecules are static (*D* < 0.1μm^2^/s), which closely relates to the estimated percentage of clustered molecules calculated from STORM data.

**Figure 3:**
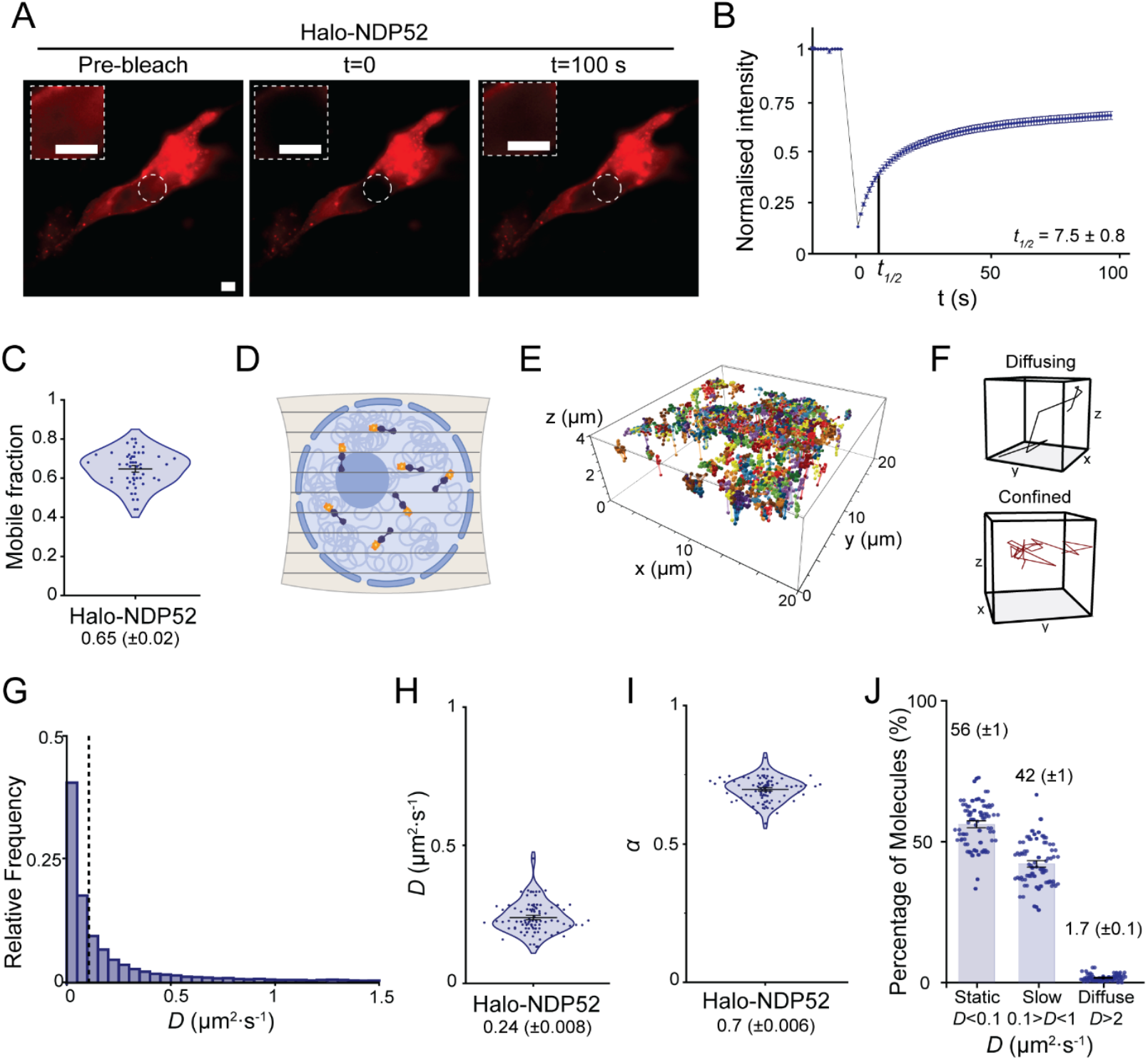
Molecular dynamics of NDP52 in the nucleus. **(A)** Example of Fluorescence Recovery After Photobleaching (FRAP) image acquired in HeLa cells transiently expressing Halo-NDP52. Insets display zoomed-in detail of nuclear area selected for photobleaching. Scale bar= 5μm. **(B)** Normalised fluorescence intensity profile, in function of time for FRAP experiments. Estimated value of fluorescence recovery *t_1/2_* is shown on the graph. Mean values ± SEM are shown. n= 27 cells. **(C)** Calculated mobile fraction from FRAP data. Mean values ± SEM are shown. n= 27 cells. **(D)** Diagram depicting simultaneous acquisition of nine focal planes (covering 4μm in z and 20μm x20μm in xy) for 3D single-molecule tracking of Halo-NDP52 in the nucleus. **(E)** Example of 3D reconstructed trajectories for a single nucleus over time. **(F)** Example of diffusive and confined trajectories over time. **(G)** Histogram of diffusion constants from the nucleus of Hela cells transiently expressing Halo-NDP52. Dotted lines represent the applied threshold to differentiate between static and dynamic molecules (14 322 molecules from 51 cells). **(H)** Diffusion coefficient values for Halo-NDP52. Each data point represents the mean diffusion coefficient for a cell. **(I)** Anomalous diffusion constant, α, values. Each data point represents the mean α value per cell. **(J)** Percentage of molecules considered static (*D* < 0.1μm^2^/*s*), slow moving (0.1 < *D* < 1 μm^2^/s) or diffuse (*D* > 1μm^2^/s) per cell. n = *5*1 cells.

Overall, the clustering behaviour and confined dynamics of NDP52 molecules in the nucleus support our hypothesis of a nuclear function for this protein.

### NDP52 oligomerisation and structure

Having investigated the spatial organisation and molecular dynamics of nuclear NDP52, which suggests a nuclear function, we wanted to investigate the biochemical properties of the protein to understand its potential roles. For this, we used different recombinant NDP52 constructs, including the full-length protein (NDP52-FL), an N-terminal truncated region (_N_NDP52), which includes the SKICH domain and part of the coiled-coil region (amino acid residues 1-190), a C-terminal region (_C_NDP52), which includes both zinc finger domains (amino acid residues 365-446) and the last zinc finger domain (ZF2 – amino acid residues 380-446) (Fig.4A). All the recombinant proteins presented stable secondary structure, as shown by circular dichroism and/or nano-differential scanning fluorimetry (nano-DSF) (Supplementary Fig.2A-K). The first zinc finger domain of NDP52 (ZF1) was not selected for biochemical studies, as it lacked a stable secondary structure. This is in agreement with previous structural reports for this domain ^7^.

**Figure 4:**
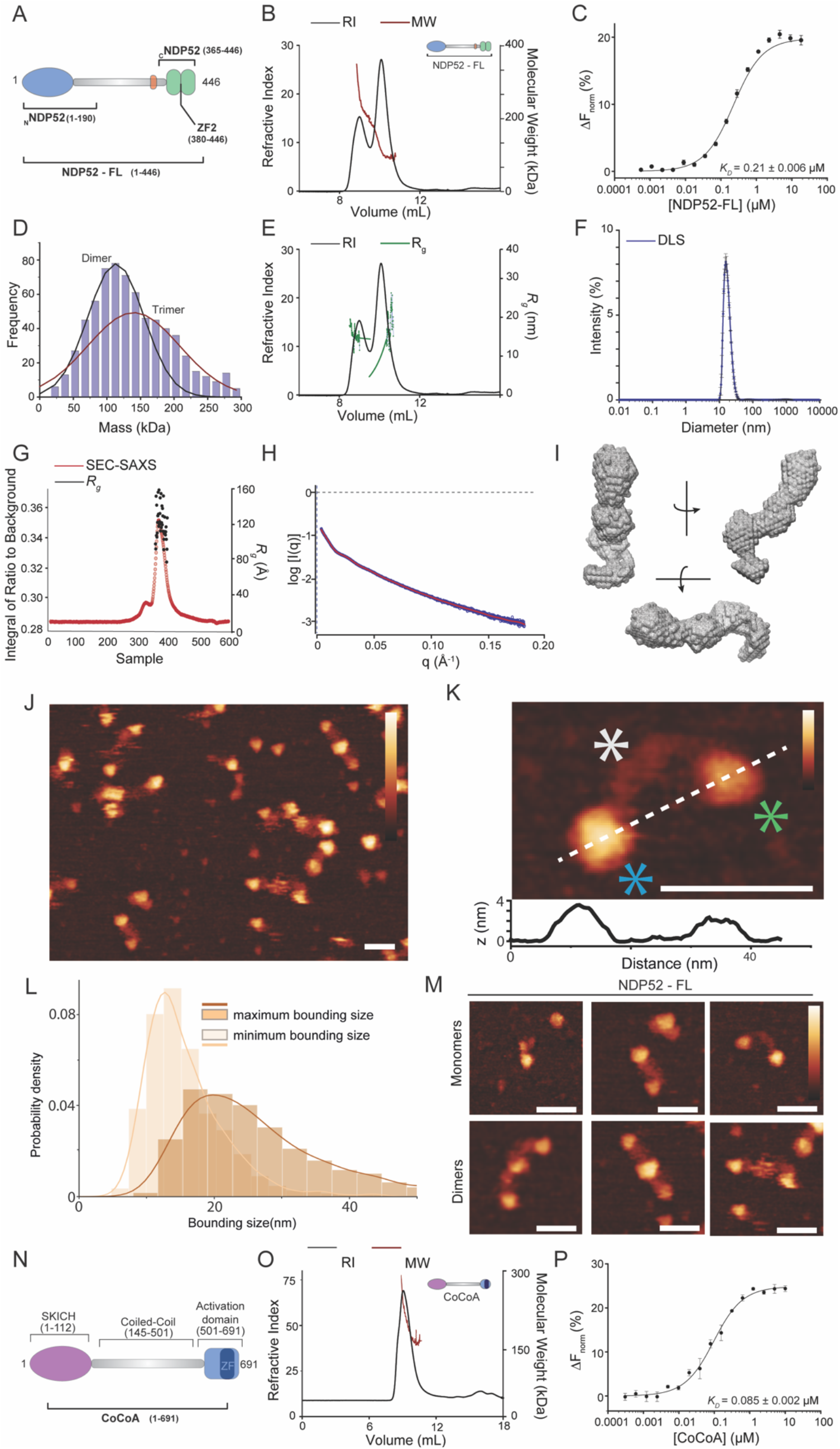
Oligomerisation and structure of NDP52. **(A)** Diagram of NDP52 showing recombinant constructs used for biochemical assays with NDP52. **(B)** Size-exclusion chromatography with multi-angle light scattering (SEC-MALS) profile for recombinant full-length NDP52 (NDP52-FL). Refractive index (RI) trace is shown in black, as well as the calculated molecular weight values, across the peaks (in red). **(C)** Microscale thermophoresis, showing oligomerisation of NDP52. Calculated *K_D_* (as specified in Methods) is displayed in the graph. Values plotted represent average ± SEM of three individual experiments. **(D)** Histogram showing calculated mass of NDP52-FL from mass photometry assays. Two Gaussian curves could be fitted to the data set. Mean values of Gaussian curves closely correspond to dimeric and trimeric molecular weights of NDP52 (Gaussian max values = 105kDa and 140kDa, respectively). **(E)** Radius of gyration (*R_g_*) calculated from SEC-MALS data shown in (B). Estimated *R_g_* for peak one was 14.84nm and 11.7nm for the second peak. RI trace shown again for NDP52-FL in black, and *R_g_* across peaks shown in green. **(F)** Dynamic light scattering trace for NDP52-FL showing calculated diameter for the protein, with values between 10 and 43 nm and maximum at 15.5 nm. **(G)** SEC-Small-angle X-ray light scattering (SEC-SAXS) for NDP52-FL showing *R_g_* values across peaks. Radii values between 9-15nm. **(H)** Experimental SAXS curve for NDP52-FL. **(I)** Beads model of NDP52-FL obtained from SEC-SAXS data. **(J)** AFM image of NDP52-FL showing multiple molecules. Scale bar = 25 nm. Height scale = 4.5 nm. **(K)** High-resolution AFM image of an individual NDP52-FL molecule. Protein domains are indicated by asterisks, with SKICH in blue, coiled coil in grey and c-terminal domain in green. Below, a line profile taken from the AFM image of the NDP52-FL protein along the white dotted line from left to right is shown. Scale bar = 25 nm. Height scale = 4.5 nm. **(L)** Histogram and kernel density estimate (KDE) plots for maximum and minimum bounding size of NDP52-FL molecules, measured from masks generated by Topostats (Supplementary Fig.3K). Peaks in KDE plots were used to determine particle size (KDE max ± SD) minimum = 13 ± 6 nm, maximum = 20 ± 12. N = 1365 particles. **(M)** AFM images of NDP52-FL showing the protein in monomeric and dimeric forms. Scale bar = 25 nm. Height scale = 4.5 nm. **(N)** Diagram depicting CALCOCO2/CoCoA, which belongs to the same family as NDP52 and has high sequence and domain similarity. Domains and key features are specified. A recombinant full-length CoCoA construct was used in biochemical assays. **(O)** SEC-MALS trace for CoCoA. RI trace is shown in black and calculated molecular weight values are shown in red (MW values between 140 and 300kDa). **(P)** Microscale thermophoresis, showing oligomerisation of CoCoA, with the calculated *K_D_* displayed in the graph. Curve fitting was performed as described in Methods. Values represent average ± SEM of three individual experiments.

Previous work showed that full-length NDP52 is mainly a dimer in solution ^6,9^. To confirm this, we used Size-Exclusion Chromatography with Multi-Angle Light Scattering (SEC-MALS).

SEC-MALS allows us to obtain accurate molecular weight information from gel filtration elution profiles and to identify different oligomeric species in solution. Our SEC-MALS data show that the majority of NDP52 is present in the dimeric form (second peak average molecular weight = 117kDa), but it also shows the presence of higher oligomeric forms, such as trimers and tetramers (first elution peak with an average molecular weight of 333 kDa) (Fig.4B). Through titrations of RED-tris-NTA labelled NDP52-FL with unlabelled NDP52-FL, microscale thermophoresis (MST) confirms oligomerisation of the full-length protein, with an estimated *K_D_* value of 0.21μM (±0.006) (Fig.4C) which supports the presence of a minimal dimer complex at the concentrations used in the SEC-MALS experiments. Mass photometry data enables mass determination at the single molecule level. At 100 nM NDP52, we observed a mostly dimeric state, with a small population of trimers also present (Fig. 4D). This is consistent with the MST and SEC-MALS analysis and we conclude that NDP52 readily oligomerizes.

From the SEC profile, NDP52 appears to be an elongated protein, eluting at a much earlier elution volume than expected for a globular protein. The estimated radii of gyration from SEC-MALS data are 11.7nm, for the second peak (corresponding to NDP52 dimers) and 14.8nm for the higher-oligomeric forms (first elution peak) of NDP52 (Fig.4E). This translates into an approximate end-to-end measurement of 23 nm for dimeric NDP52. To directly measure NDP52 particle size, we used Dynamic Light-Scattering (DLS) which showed that NDP52-FL particles can be measured at a range of diameters between 10-43nm, with a maximum at 15.5 nm (Fig.4F). To obtain more information regarding the overall shape of NDP52-FL, we used Small-Angle X-Ray Scattering (SAXS). SEC-SAXS is a robust technique for the study of macromolecule conformation in solution. Our SAXS data estimate a radius of gyration for NDP52 between 9-15 nm (end-to-end value 18-30 nm) (Fig.4G), consistent with a predicted rod-shape structure for the protein. Variability in radii measurements for NDP52 could be a direct consequence of its elongated shape, as measurements for different profiles of the protein will be more varied than in a globular protein. Using SAXS we were also able to generate an envelope model for NDP52-FL, showing the predicted elongated shape (Fig. 4H and I).

To directly visualise and measure protein shape and size, we used Atomic Force Microscopy (AFM) imaging. In agreement with the biochemical data (Fig.4B-H), AFM imaging of NDP52-FL shows a distribution of proteins of elongated shape (Fig.4J). The resolution of AFM imaging resolves the different domains of the NDP52-FL protein, with the larger SKICH domain and smaller C-terminal region distinguishable by height and linked together by a thin, flexible linker (Fig.4K). The length of the protein (maximum bounding size) has a wide distribution with a clear peak at 20 ± 12 nm (Fig.4L), as expected from the SAXS data. This variability is driven by the thin, coiled coil, flexible linker which can adopt a variety of conformations, allowing the protein to bend, and leading to variability in the protein length.

In agreement with the observed elongated shape for the protein the width (minimum bounding size) of NDP52-FL was significantly less than the length with a peak at 13 ± 6 nm (Fig.4L). The widths of NDP52-FL also occupy a narrower distribution compared to the lengths since the coiled-coil only allows for flexibility along the length of the protein (Fig.4K). It is therefore likely that the width of NDP52-FL corresponds to the diameter of the globular domains at NDP52-FL ends (Fig.4K). To probe this hypothesis, we measured the dimensions of a truncated version of the protein, _C_NDP52 (Fig.4A). AFM imaging (Supplementary Fig.3C) showed that the minimum and maximum bounding sizes for _C_NDP52 largely overlap, with the peak in the probability distributions occurring at values of 13 ± 6 nm and 9 ± 3 nm respectively, indicating relatively globular conformations (Supplementary Fig.3D). These measurements closely match the minimum bounding size of NDP52-FL (Supplementary Fig.3E), showing that the width of NDP52-FL is determined by the size of its globular domains.

Although α-helical coiled-coil domains are often drivers of protein oligomerisation ^23,24^, AFM imaging also showed the protein’s terminal domains acting as the interface for dimerisation of the protein (Fig.4M). Dimers were observed as even longer elongated molecules, with two smaller globular domains linked by two thin linkers to one central globular domain, most likely formed of two terminal regions. To test which regions of NDP52 are capable of oligomerising, we used different truncated regions of the protein (Fig.4A). SEC-MALS of _C_NDP52 shows that this region is mostly present in dimeric and monomeric forms, although trimers could also be detected (Supplementary Fig.3A). MST data confirms oligomerisation of this domain, generating a *K_D_* of 0.05 μM (±0.006) (Supplementary Fig.3B). These observations were confirmed by AFM imaging where we could identify monomeric, dimeric and trimeric forms of _C_NDP52 (Supplementary Fig.3C). Oligomerisation of _C_NDP52, which lacks the presence of the coiled-coil region, suggests that these domains might also be important for interactions between monomers during dimerization of the full-length protein.

We also investigated the ability of the ZF2 domain to oligomerise, using SEC-MALS and MST. Our data show that this domain can also homo-oligomerise, presenting itself as a monomer, dimer and trimer in solution, with an oligomerisation *K_D_* of 0.18 μM (±0.017) (Supplementary Fig.3E and F). When testing oligomerisation of the N-terminal region of NDP52, _N_NDP52, containing the SKICH domain and part of the coiled-coil region, we observe a clearer preference for the dimeric form (Supplementary Fig. 3G and H). Interestingly, we could also observe an interaction between _c_NDP52 and _N_NDP52 (Supplementary Fig. 3I). It is possible that these two opposing regions interact in the full-length protein, due to the presence of a relatively flexible central coiled-coil region, or between homo-oligomers of NDP52.

As previously mentioned, NDP52 shares high sequence identity with its family member CoCoA - a protein with known nuclear functions in transcription co-activation ^6^. However, very little is known regarding the oligomeric states of CoCoA, or how this may align with NDP52. To test if recombinant CoCoA (Fig.4L) can also form dimers, we used SEC-MALS. Our data show that the main peak for CoCoA is a complex mixture of molecular weights, ranging from 148kDa (equivalent to the dimeric form of CoCoA) to 300kDa (Fig.4M). Using MST, we further confirmed the ability of CoCoA to oligomerise, with a calculated KD of 0.085 μM (±0.002) (Fig.4N). Essentially, CoCoA and NDP52 display similar biochemical properties.

### NDP52 binds and oligomerises with doublestranded DNA

Having determined the oligomeric state of NDP52-FL and clarified its nuclear localisation, we decided to test NDP52 binding to DNA. Previously, Fili *et al*. showed that NDP52 can bind to double-stranded DNA (dsDNA) with high-affinity ^15^. As we have established that NDP52 is present in confined clusters within the nucleus, DNA binding could be an essential part of its nuclear role. Hence, we used an electrophoretic mobility shift assay (EMSA) to investigate the formation of NDP52-dsDNA complexes. Using 250nM of dsDNA 40bp long (ds40) and concentrations of NDP52-FL ranging from 50 nM to 3μM, we show that NDP52 can form complexes with dsDNA, *in vitro*, evident by the formation of a higher band in the EMSA (Fig.5A). To explore this interaction in a quantitative manner, we used fluorescence spectroscopy. For this, NDP52-FL was titrated against two different lengths of FITC labelled DNA - 40 and 15bp (ds40 and ds15, respectively). Our data confirmed a high-affinity interaction between NDP52 and DNA, with *K_D_* values < 100nM for both DNA lengths (Fig.5B). To directly visualise this interaction, once again we employed AFM imaging. We used linearised dsDNA 339 bp long (ds339) - approximately 115 nm long to observe direct interactions between NDP52-FL and DNA (Fig.5C). We can observe direct interactions between NDP52-FL and ds339. Furthermore, we can also observe that more than one molecule of NDP52-FL can interact with DNA (Fig.5C and Fig.6A-B). This agrees with mass photometry measurements that show that when incubated with ds40, the measured mass for NDP52-FL increases from its dimer/trimer values (112 and 157 kDa, calculated for NDP52-FL alone, to 1334 kDa) (Fig.5D). Similar to NDP52, when testing CoCoA for dsDNA binding, we also observe high-affinity interactions in fluorescence spectroscopy assays (Fig.5E).

**Figure 5:**
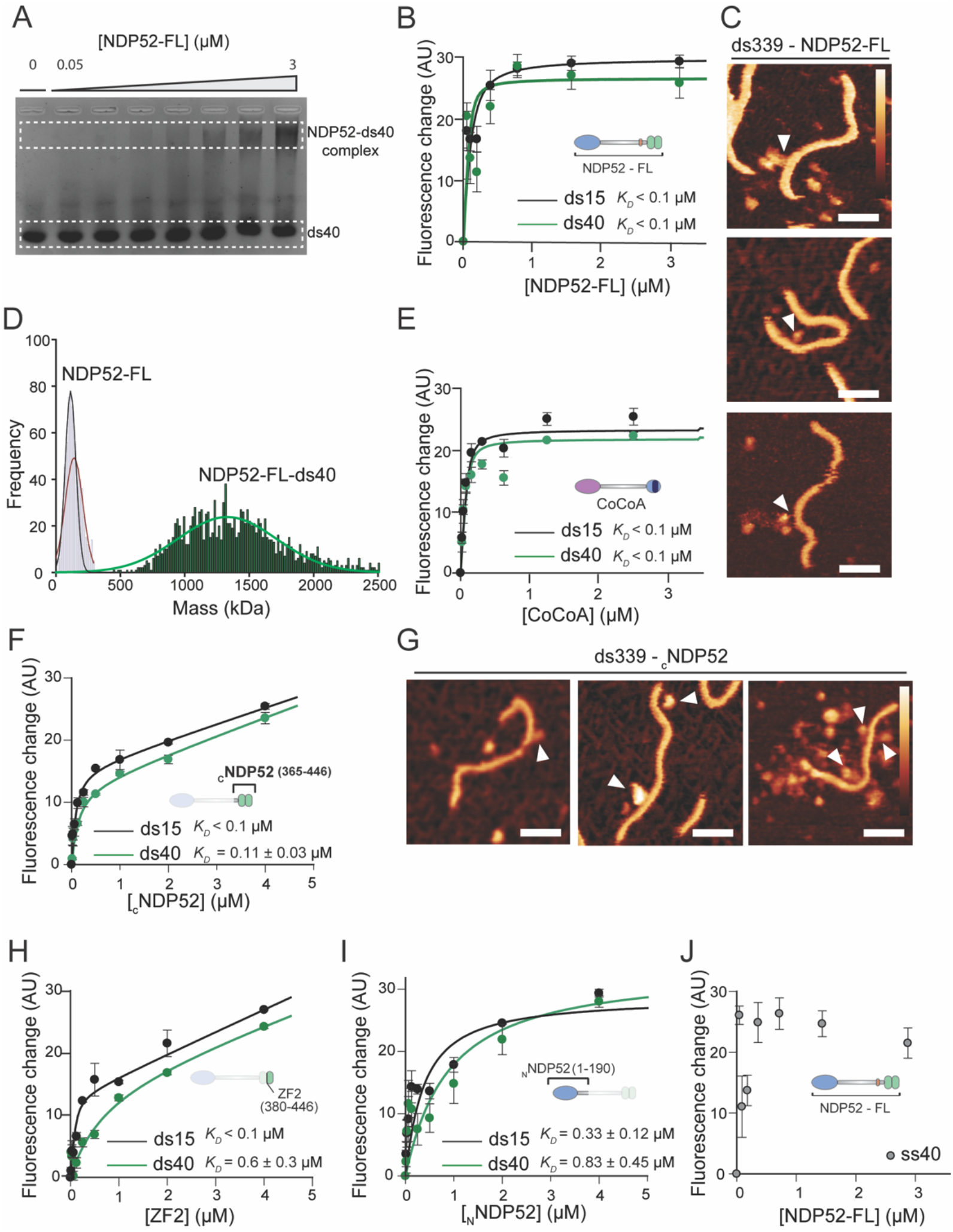
NDP52 binds to and oligomerises with double-stranded DNA through its C-terminal domain. **(A)** Electrophoretic-mobility shift assay (EMSA) for NDP52-FL with ds40. dsDNA was used at 250 nM, with increasing concentrations of NDP52-FL run in each well (0.05; 0.1; 0.2; 0.5; 1; 2 and 3 μM). Lower band represents free ds40 and top band represents DNA in complex with NDP52-FL. **(B)** Fluorescence spectroscopy titrations of NDP52-FL against 40bp or 15bp fluorescein amidite double-stranded DNA (ds40 and ds15, respectively). **(C)** AFM images showing direct visualisation of NDP52-FL binding to linear 339bp DNA (ds339). Binding events are marked by white arrowheads. Scale bar = 25 nm. Height scale = 4.5 nm (scale bar inset in C and G). **(D)** Mass photometry histogram for NDP52 showing a large shift in detected mass when NDP52-FL is incubated with ds40. Histograms and Gaussian fittings for NDP52 alone are the same as the ones shown in Figure 4D (in black and red). Histogram for NDP52-FL-ds40 (green) was also fitted to a Gaussian function. Mean value calculated as 1 334kDa). **I** Fluorescence spectroscopy titrations of CoCoA against ds40 and ds15DNA. **(F)** Fluorescence spectroscopy titrations of _C_NDP52 against ds40 and ds15. Calculated *K_D_* values are shown. **(G)** AFM images showing _C_NDP52 binding and clustering around linear ds339. Scale bar = 25 nm. Height scale = 4.5 nm (scale bar inset in C). **(H)** Fluorescence spectroscopy titrations of ZF2 against ds40 and ds16 DNA. **(I)** Fluorescence spectroscopy titrations of _N_NDP52 against ds40 and ds15 DNA. **(J)** Fluorescence spectroscopy titrations of NDP52-FL with single-stranded 40bases DNA (ss40). For all protein-DNA fluorescent assays, DNA concentration was kept at 100 nM and *K_D_* values represent mean ± SEM of n=3 independent experiments. Data fitting was performed as described in Methods.

**Figure 6:**
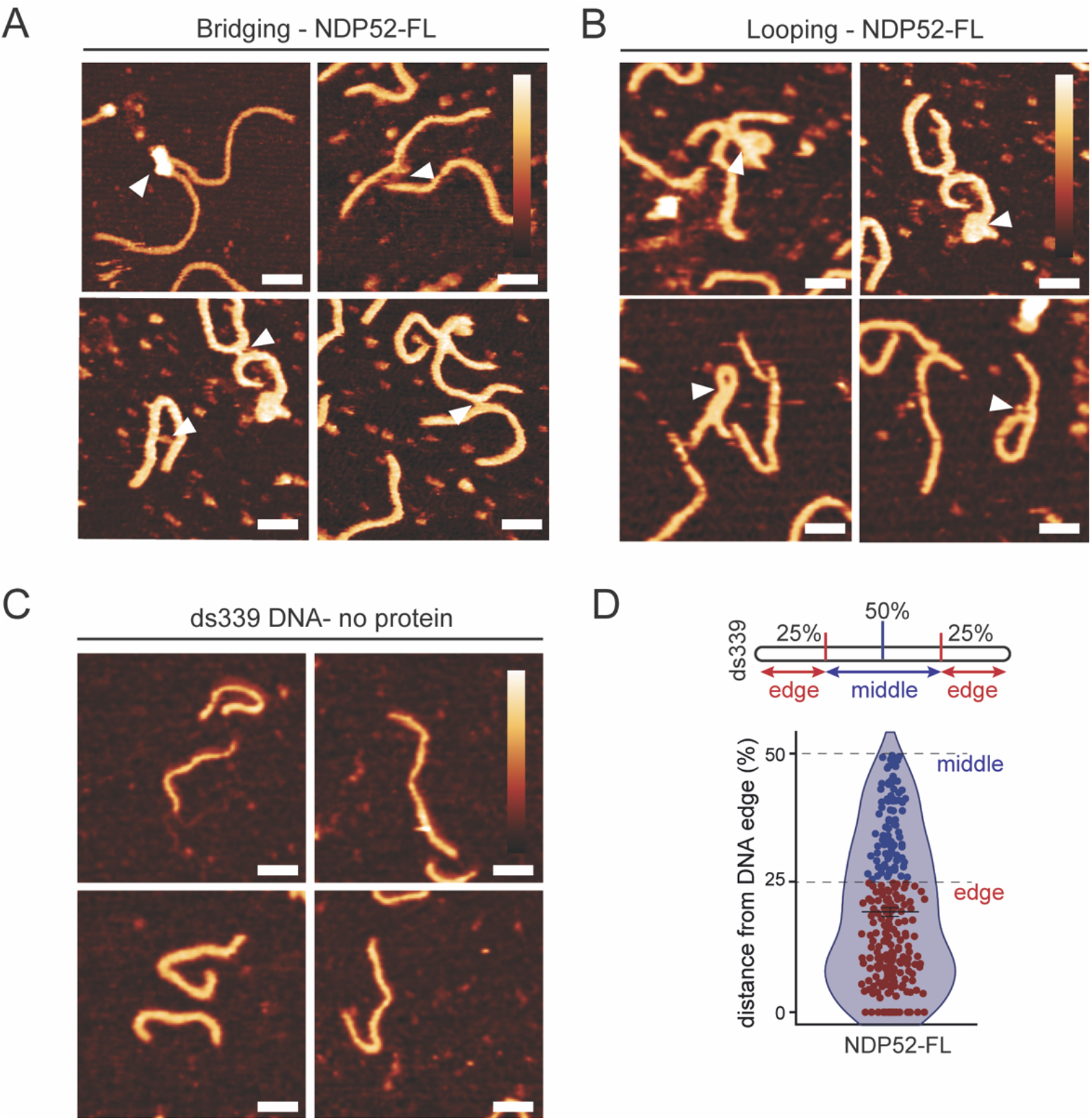
NDP52 changes DNA conformation *in vitro*. **(A)** AFM images of NDP52-FL bridging strands of linear (339 bp) dsDNA. Scale bar = 25 nm. Height scale = 4.5 nm **(B)** AFM images showing looping of linear ds339 DNA following incubation with NDP52-FL. Scale bar = 25 nm. Height scale = 4.5 nm **(C)** AFM images of linear ds339 DNA without incubation with protein. Scale bar = 25 nm. Height scale = 4.5 nm. **(D)** Preference for NDP52-FL binding on ds339 molecule. Diagram shows ds339 edge and middle references on linear DNA. Violin plot shows %distance from DNA edge values for NDP52-FL binding. Mean ± SEM is shown for n=270 binding events.

Zinc finger domains are well-known for their ability to bind DNA ^25^. To test if these domains are responsible for DNA binding abilities of full-length NDP52, we used the _C_NDP52 and ZF2 recombinant constructs in fluorescence spectroscopy assays. As expected, both constructs interact tightly with dsDNA, with *K_D_* values in the low nM range (Fig.5F and H). However, DNA binding curves for these constructs do not reach saturation at higher concentrations of protein, as they do for NDP52-FL. This could be explained by the clustering behaviour of _C_NDP52 around DNA, that we observe by AFM imaging of this domain complexed with ds339 (Fig.5G). _C_NDP52 oligomerises with, and around dsDNA, which can create very large DNA-protein complexes (Fig.5G - right).

Although some degree of interaction could be detected for _N_NDP52-dsDNA, this presents much lower affinity than NDP52-FL or C-terminal domains. This is represented by lower *K_D_* values estimated from fluorescence spectroscopy assays, and relatively poor fitting of these curves (Fig.5I).

Interestingly, we also observe specificity of NDP52-FL towards dsDNA. Although fluorescence spectroscopy assays using FITC labelled single-stranded DNA 40 bases (ss40) show changes in fluorescence, the data are highly variable and could not be fitted to a binding equation (Fig. 5J). This suggests that although some interaction may occur, NDP52-FL preferentially binds dsDNA.

Since we have established that NDP52 can bind to DNA *in vitro*, we hypothesised that this could also occur in cells, whereby NDP52 could directly interact with genomic regions. To test this, we performed chromatin immunoprecipitation (ChIP) with NDP52. Using this approach, we could detect the presence of NDP52 bound to chromatin-enriched cellular fractions (Supplementary Fig. 4A). We also tested different genomic *loci* for the presence of NDP52, through ChIP-qPCR, including genes regulated by nuclear receptors (previously linked to coactivator functions of CoCoA) ^11^, and inflammation-related targets, where NDP52 has been shown to have a role ^26,27^. Our ChIP-qPCR data suggest that NDP52 is present throughout the gene body of different genes (Supplementary Fig. 4B). This supports our hypothesis that NDP52 can bind DNA *in vitro* and *in cellulo* and this could be a mechanism through which the protein could impact gene expression.

### NDP52 can alter DNA shape

Having established that NDP52-FL can interact with DNA through its zinc finger domains, we then investigated if these interactions could cause local changes to DNA shape or structure. Using AFM imaging, we observed several instances where NDP52 appears to be able to bridge individual linear strands of dsDNA (Fig.6A). We also observed looping of DNA (Fig.6B) following incubation with NDP52-FL which was not observed in DNA samples incubated without NDP52-FL (Fig.6C and Supplementary Fig.4C). To test if NDP52-FL has a preferential binding location on linear dsDNA, i.e. at the flexible DNA ends vs the constrained dsDNA in the middle, we divided ds339 molecules into two regions. These two regions were the edge – accounting for 50% of the DNA molecule (25% at each end) and middle – accounting for the central 50% of ds339 (Fig.6D diagram). We observed that NDP52-FL preferentially binds at the ends of linear dsDNA, approximately 20-25 nm into the ds339 molecule (corresponding to 19.39±0.86% of ds339 length) (Fig.6D). This bias may be due to the extra conformational flexibility around the ends of the DNA strands and the confinement of the centre of the molecule.

Interestingly, we could also observe changes in DNA structure when incubating ds339 with _C_NDP52, with instances of DNA bridging and looping also observed (Supplementary Fig.4D and E). This, together with the fact that the N-terminal of NDP52 only displays low affinity for dsDNA in biochemical studies, strongly suggests that changes to DNA shape caused by the fulllength protein are most likely induced by the zinc finger domains at the C-terminal of NDP52.

### NDP52 in involved in RNAPII-dependent transcription

As previously mentioned, one of the most well-known binding partners of NDP52 is MVI. Previous work has linked the interaction between NDP52 and MVI to the enhancement of RNA Polymerase II (RNAPII) activity ^15^. Furthermore, colocalising foci of NDP52 and RNAPII have been previously observed in the nucleus ^15^, also shown in Fig.7A.

**Figure 7:**
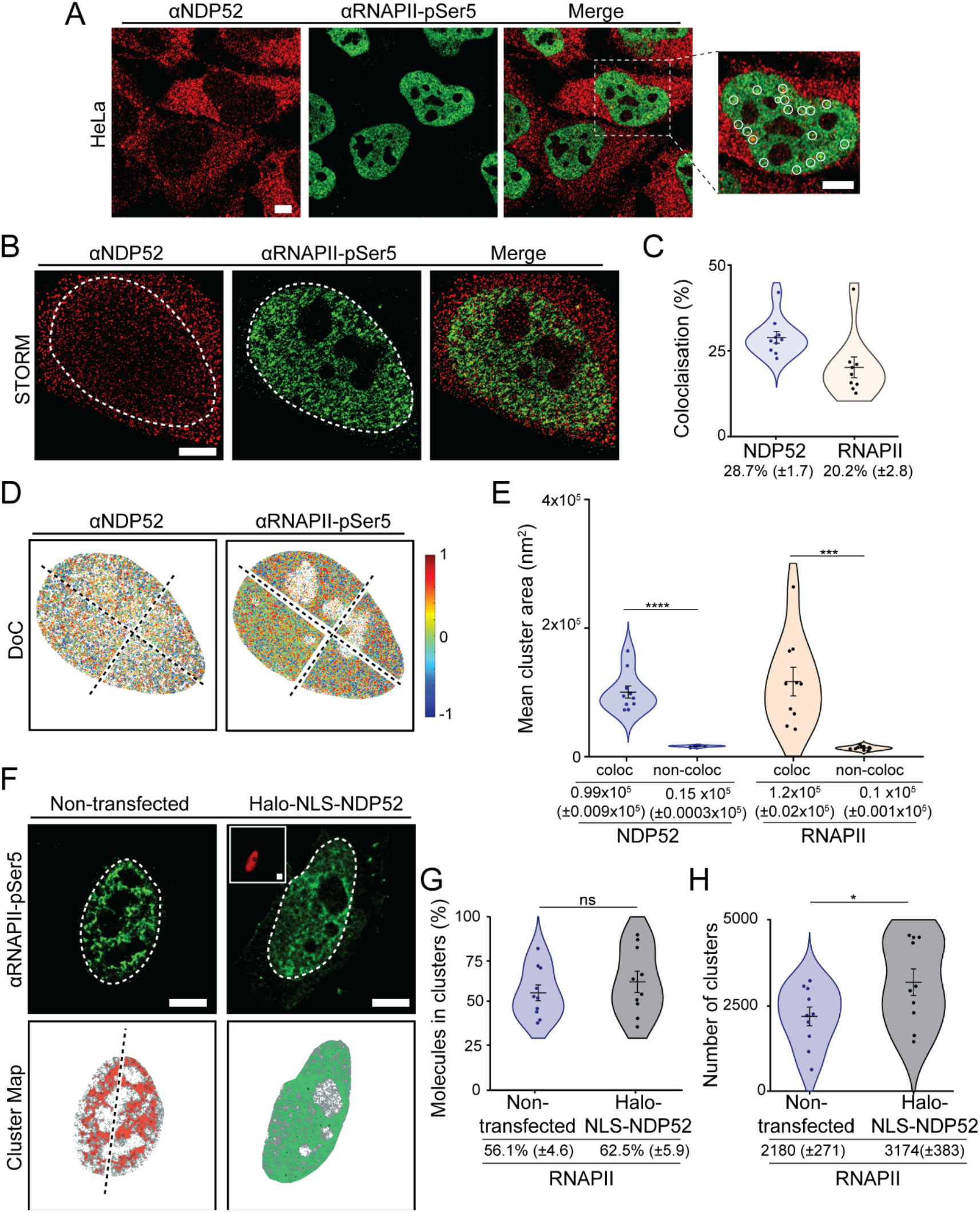
Colocalisation of NDP52 with RNAPII-pSer5. **(A)** Immunofluorescence confocal image of NDP52 (red) and RNAPII-pSer5 (green) in HeLa cells, showing detail of colocalising foci in white circles (zoomed-in right panel). Scale bar = 5μm **(B)** Example STORM images of NDP52 and RNAPII-pSer5 in HeLa. The nuclear region (determined by RNAPII-pSer5 fluorescence) was used for ClusDoC analysis (shown as dotted white line). Scale bar = 5μm **(C)** Colocalisation analysis of NDP52 and RNAPII-pSer5 clusters. **(D)** Cluster colocalisation heat maps for NDP52 and RNAPII-pSer5 generated from the STORM data shown in (B). DoC score of 1 represents perfect colocalisation between molecules, and DoC score −1 represents segregation. A DoC score of 0.4 was used as threshold for colocalisation. Due to high molecular density the nucleus was split into four ROIs for ClusDoC analysis. Axis of separation for the images are shown as dotted lines. **(E)** Mean cluster area is shown for colocalised and non-colocalised clusters of NDP52 and RNAPII-pSer5. n = 10 cells. **(F)** STORM rendering for RNAPII-pSer5 and generated cluster maps for HeLa cells non-transfected or transiently expressing a Halo-NLS-NDP52 construct. Inset in Halo-NLS-NDP52 panel shows wide-field channel Halo ligand-JF549 labelled cells. Scale bar = 5μm. **(G)** Percentage of RNAPII-pSer5 molecules in clusters in non-transfected or transiently expressing Halo-NLS-NDP52 HeLa cells. **(H)** Number of RNAPII-pSer5 clusters in non-transfected or transiently expressing Halo-NLS-NDP52 HeLa cells. Mean ± SEM values are shown. Each point represents the average value per cell. n = 10 cells (Nontransfected) n = 10 cells (Halo-NLS-NDP52). *p<0.05; ***p<0.001; ****p<0.0001 by two-tailed t-test.

To further explore the role of NDP52 in RNAPII-related transcription, we used STORM. STORM not only allows us to improve colocalisation estimates between NDP52 and RNAPII molecules, relative to conventional optical microscopy, but also allows us to measure colocalisation of clusters for both proteins. Here, we used phospho-Ser5-RNAPII immunofluorescence staining, which selects for the pool of RNAPII molecules involved in transcription initiation. STORM data show that, under normal conditions, approximately 28.7% (±1.7) of NDP52 is colocalised with RNAPII, and 20.2% (±2.8) of RNAPII is found colocalising with NDP52 (Fig. 7B and C). Colocalisation of clusters between NDP52 and RNAPII can also be observed in ClusDoC-generated heat maps and histograms (Fig. 7D and Supplementary Fig.5A), with nuclear regions of high colocalisation density for each channel represented in red. Interestingly, our data show that NDP52 clusters that colocalise with RNAPII clusters are approximately 6.5 times larger than non-colocalised clusters, and RNAPII clusters colocalised with NDP52 approximately 12-fold larger (Fig. 7E). Although there are more non-colocalised clusters than colocalised between NDP52 and RNAPII, colocalised clusters also have higher density of molecules (2.5 times higher density for NDP52 and 2 times higher for RNAPII) (Supplementary Fig. 5B-E) This further suggests a relationship between the nuclear organisation of NDP52 and transcription.

To test if NDP52 can affect the spatial organisation of RNAPII, we overexpressed the nuclear pool of NDP52. For this, we used a Halo-NLS-NDP52 construct (Supplementary Fig. 5F). We then used STORM and cluster analysis to quantify changes in the distribution of RNAPII in the nucleus (Fig.7F). Overexpression of nuclear NDP52 did not have an effect on the number of molecules of RNAPII, or the propensity for RNAPII to form clusters (Fig.7G, Supplementary Fig.5H-J). However, we did observe a significant increase in the number of RNAPII clusters in cells transiently expressing Halo-NLS-NDP52 (Fig.7H). This suggests that overexpression of NDP52 might allow the formation of new transcription hubs in the cell, but the overall size of each cluster is not dependent on NDP52.

Having determined that NDP52 can be found clustering at transcription initiation sites, we set out to explore how depletion of NDP52 would affect global gene expression. For this, we performed RNA-Seq in both HeLa and MCF-7 cells, following siRNA knockdown of NDP52 (Supplementary Fig.6 and Supplementary Fig.7). Overall, we observed significant changes in gene expression levels for both cell lines, with 1420 genes and 1140 genes differentially expressed in HeLa and MCF-7, respectively (−0.5>log_2_FC<0.5, and p_adj_<0.05) (Supplementary Fig.6A, Supplementary Fig.7A). In both HeLa and MCF-7 datasets, more genes are downregulated than upregulated, also showing an overall negative impact on transcription caused by depletion of NDP52. Gene Ontology analysis for both up and downregulated genes was then performed for both cell lines (Supplementary Fig.6B, Supplementary Fig.7B). For HeLa, genes involved in the ‘regulation of transcription’, as well as ‘cell migration’ and ‘tissue development’ were significantly affected (Supplementary Fig.6B). In MCF-7, NDP52 knockdown was shown to also affect the expression of genes involved in ‘cell migration’, ‘tissue development’, as in HeLa, but also ‘cell cycle’, ‘DNA replication’ and ‘chromosome segregation’ (Supplementary Fig.7B).

This suggests that whilst some genes and processes are equally affected by NDP52 knockdown in both cell lines, others might be more susceptible depending on the unique characteristics of the cell line.

### NDP52 nuclear interactome

Following our observation that NDP52 colocalises and clusters at RNAPII transcription initiation sites, and can drive the formation of additional RNAPII clusters when overexpressed, we decided to explore the nuclear interactome of NDP52. To identify partners of NDP52, we used label-free quantitative LC-MS/MS to analyse pulldowns of recombinant NDP52-FL, _c_NDP52, CoCoA and ZF2 from HeLa nuclear extracts (Supplementary Fig.8A, Fig. 8A-D, Supplementary Fig.8B and C). Proteins were identified using Progenesis software (Waters), as described in Methods. Following protein identification, we selected proteins enriched in NDP52-FL pull-downs, compared to control beads without recombinant protein. We used log_2_FC>1 and p_adj_<0.05 as a threshold to investigate the interactome of NDP52. Following this, we performed GO analysis to determine novel biological processes that could shed light on a new nuclear role for NDP52. Interestingly, the top enriched biological processes for NDP52 interactions were ‘DNA geometric change’ and ‘DNA duplex unwinding’, with ‘chromosome organisation’ also scoring high (Supplementary Fig.9A). Top enriched molecular functions also relate to ‘nucleosome-dependent ATPase activity’, ‘DNA helicase activity’ or DNA binding (Supplementary Fig.9B). These data reinforce the concept that NDP52 has significant roles in DNA binding and structure, as indicated in our biochemical studies and AFM imaging, which could, in turn, impact gene expression. Although several known transcription factors were identified in proteomics (Fig.8B), gene expression-related GO functions were not particularly enriched. This suggests that although NDP52 might directly interact with transcription-related proteins and also affect gene expression through these interactions, its largest contribution might, instead, arise through regulation of DNA structure.

**Figure 8:**
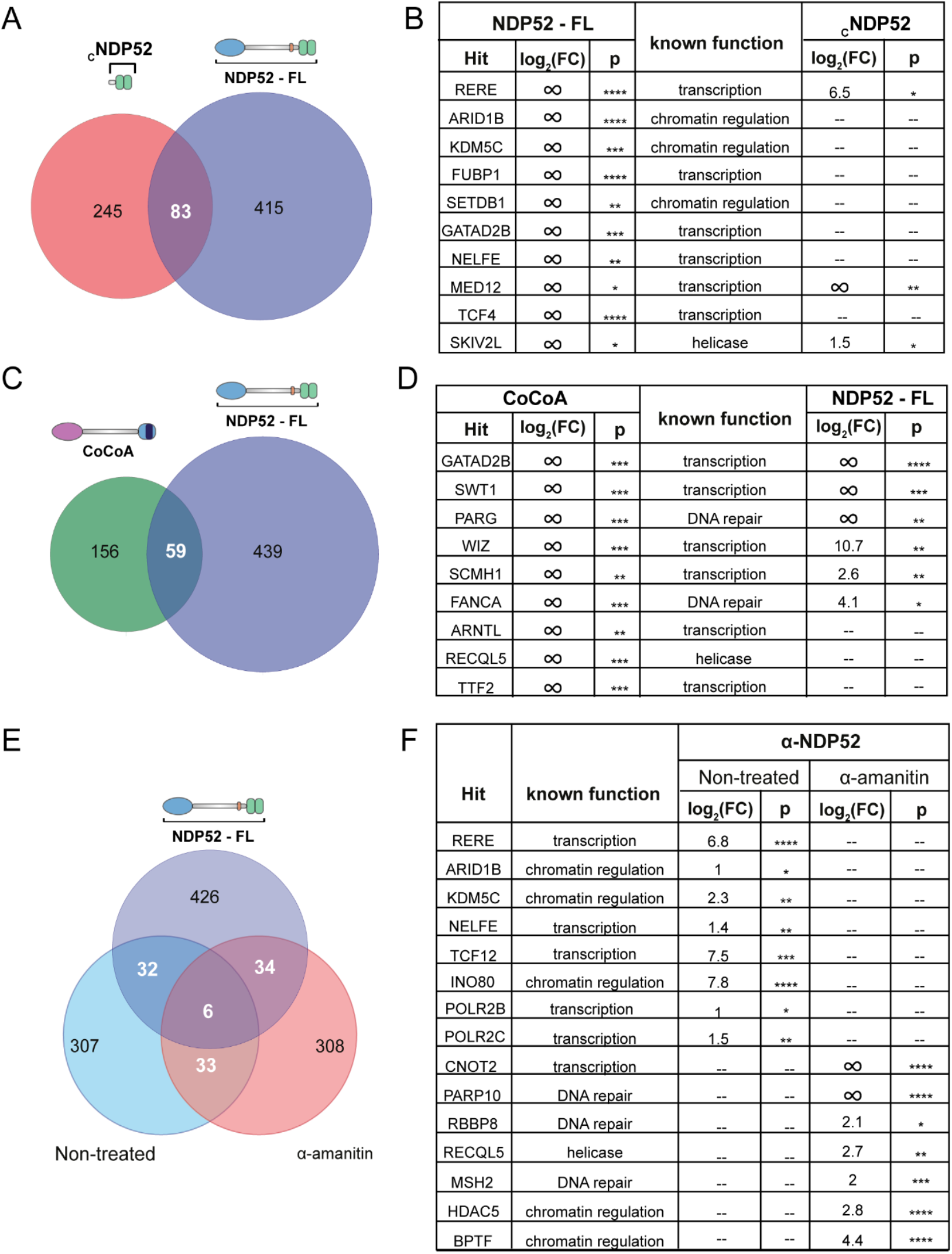
NDP52-FL, CoCoA and _C_NDP52 interactomes from HeLa nuclear extract. **(A)** Venn diagram of hits found in NDP52-FL and _C_NDP52 **(B)** Examples of top hits for NDP52, and their identification in _C_NDP52 proteomics data. log_2_FC is relative to beads control. **(C)** Venn diagram of hits found in NDP52-FL and CoCoA **(D)** Examples of top hits for CoCoA, and their identification in NDP52-FL proteomics data. log_2_FC is relative to beads control. **(E)** Venn diagram showing overlap of identified hits between recombinant NDP52-FL proteomics and coimmunoprecipitation of endogenous NDP52 for non-treated and α-amanitin treated cells. **(F)** Examples of hits identified in non-treated and α-amanitin treated cells. log_2_FC is relative to beads control. *FDR<0.05, **FDR<0.01, ***FDR<0.001, ****FDR<0.0001.

We were able to map 16% of NDP52-FL interactions to the C-terminal of NDP52 (_C_NDP52), also through LC-MS/MS with recombinant _C_NDP52 (Fig.8A). Figure 8B shows examples of top hits for NDP52-FL and the identification of some of these hits in _C_NDP52 proteomics. Fold change values, are relative to control beads for pull-downs, with infinity foldchange (∞) indicating hits only present in pulldowns and not in control samples.

We also compared how the interactome of NDP52 relates to its close family member CoCoA (Fig.8C). Similar to NDP52, the nuclear interactome of CoCoA also showed enrichment for ‘DNA duplex unwinding’ and ‘DNA geometric change’. However, ‘gene expression’ was clearly enriched for CoCoA (Supplementary Fig.10A and B). Furthermore, when comparing CoCoA and NDP52 interactions, a quarter of CoCoA hits were common to NDP52 (Fig.8C), showing a degree of overlap between both interactomes, as expected for proteins with high homology. Importantly, our data show that whilst both proteins could have similar functions and overlapping interactomes, they do not appear to be redundant. Figure 8D shows some of the top hits for CoCoA and identification in NDP52-FL pull-downs.

As ZF2 is highly conserved in both NDP52 and CoCoA, we also tested if some of the interactions in common between CoCoA and NDP52 could be mapped to this domain (Supplementary Fig.8B). Interestingly, only 5% of common interactions between NDP52 and CoCoA occur independently of ZF2-binding, suggesting that this domain could account for similarities in the interactomes between both proteins. Furthermore, 86% of proteomics common hits between NDP52-FL and _C_NDP52, could be mapped to ZF2. This, together with the fact that ZF1 is largely unstructured, could indicate that protein-protein interactions at the C-terminal of NDP52 are mostly sustained by ZF2. Examples of top hits for CoCoA and NDP52 are shown for ZF2 proteomics in Supplementary Fig.8C.

### Changes to the nuclear organisation and dynamics of NDP52 following transcription inhibition

As we have established that NDP52 has a role in transcription, and can impact the organisation of RNAPII, we tested whether transcription inhibition would affect its nuclear organisation and dynamics. To address this, we used α-amanitin, an irreversible RNAPII inhibitor that promotes degradation of RNAPII ^28^. As expected, α-amanitin treatment leads to depletion not only of RNAPII molecules, but also of its clusters (Fig. 9A-D and Supplementary Fig.11A-C).

**Figure 9:**
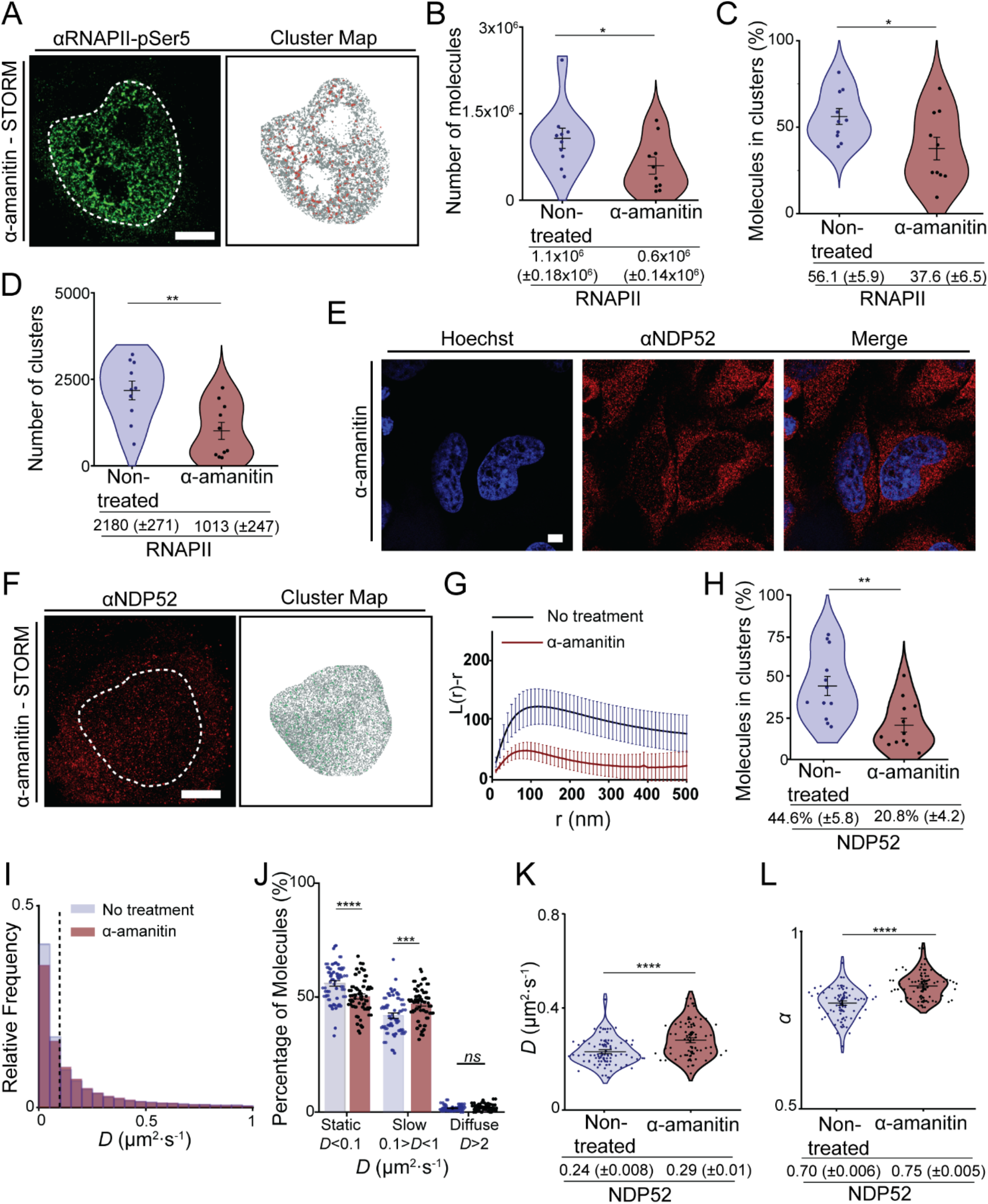
Organisation and dynamics of nuclear NDP52 following transcription inhibition. **(A)** Example of STORM rendering and cluster map, generated following DBSCAN analysis, of RNAPII-pSer5 following α-amanitin treatment. Scale bar = 5μm. **(B)** Calculated number of RNAPII-pSer5 molecules in the nucleus of non-treated vs α-amanitin treated cells. **(C)** Percentage of RNAPII-pSer5 molecules in clusters for non-treated vs α-amanitin treated cells. **(D)** Number of clusters in selected ROIs for RNAPII-pSer5 in non-treated vs α-amanitin treated cells. n = 10 cells (non-treated) n = 10 cells (α-amanitin). **(E)** Confocal image of NDP52 in HeLa cells following treatment with transcription inhibitor α-amanitin. Scale bar = 5μm. Hoechst DNA stain is shown in blue. **(F)** Example STORM image of NDP52 in HeLa cells treated with α-amanitin and corresponding cluster map. Scale bar = 5μm. Clustered molecules are shown in green. **(G)** Linearized Ripley’s K Function, *L*(*r*)-*r*, calculated for selected ROIs from STORM images. Ripley’s K values are shown in red for α-amanitin treated cells and in blue for non-treated cells. For nontreated cells, values are the same as shown in Figure 2C. Mean values are plotted ± SEM. n= 11 (α-amanitin) n=12 (non-treated). **(H)** Percentage of molecules in clusters in non-treated HeLa cells compared to α-amanitin treatment. Values for non-treated cells are the same as shown in Figure 2G. Mean ± SEM values are shown. n= 11 (α-amanitin) n=12 (non-treated) ** p<0.01 by a two-tailed t-test. **(I)** Histogram of diffusion constants from the nucleus of non-treated and α-amanitin-treated Hela cells transiently expressing Halo-NDP52 in blue and red, respectively. Dotted lines represent the applied threshold to differentiate between static and dynamic molecules (14 322 molecules from 51 cells for non-treated condition and 14 492 molecules from 50 cells). Non-treated cell values are the same as shown in Figure 3G **(J)** Percentage of molecules considered static (*D* < 0.1μm^2^/s), slow moving (0.1 < *D* < 1 μm^2^/s) or diffuse (*D* > 1 μm^2^/s) per cell. n = 51 cells. (non-treated – same as Figure 3J) and n = 50 (α-amanitin). **(K)** Diffusion coefficient values for Halo-NDP52 under normal conditions and after α-amanitin treatment. Each data point represents the mean diffusion coefficient for a cell. n = 51 (non-treated – same as Figure 3H) and n = 50 (α-amanitin). **(L)** Anomalous diffusion constant, α, values under normal conditions and after α-amanitin treatment.

We then used STORM to compare the spatial organisation of NDP52 in the nucleus of nontreated HeLa cells versus cells treated with α-amanitin (Fig. 9E-F). The linearised Ripley’s K function clearly shows that, compared to nontreated cells, there is a reduced probability for NDP52 clustering in the nucleus, following α-amanitin treatment (Fig. 9G). This is further confirmed through cluster analysis, which shows a reduction from 44.6% (±5.8) to 20.8% (±4.2) in the percentage of NDP52 molecules forming clusters following α-amanitin treatment (Fig. 9H).

We also explored how reduced clustering of NDP52 in the nucleus, following transcription inhibition, would affect its molecular dynamics. As the number of NDP52 in clusters is markedly reduced, we expected an increase in the dynamic behaviour of the protein. To test this hypothesis, we used acMFM to determine diffusion coefficient and anomalous diffusion changes in cells transiently expressing Halo-NDP52, in nontreated and α-amanitin-treated cells. As expected, loss of NDP52 clusters, observed in STORM data, correlated with a significant increase in the diffusion coefficient and anomalous diffusion constant, α, for Halo-NDP52 in the nucleus of α-amanitin-treated cells (Fig. 9I-L, Supplementary Fig.11D). A proportion of static NDP52 molecules was lost following α-amanitin treatment (reduction from 56.3% (±1.2) in nontreated to 50.5% (±1.1) in α-amanitin-treated) (Fig.9J). This was accompanied by a significant increase in molecules in slow diffusion (increase from 42.1% (±1.2) in non-treated to 47.4 (±1.0) in α-amanitin-treated) and a small, non-significant, increase in diffuse molecules (from 1.7% (±0.1) in non-treated to 2.1 (±0.2) in α-amanitin-treated). Overall, our data show that transcription inhibition by α-amanitin disrupts global nuclear NDP52 clustering, which correlates with higher molecular diffusion of the protein.

Having shown that the spatial organisation of nuclear NDP52 is altered following transcription inhibition, we also tested if this would also cause changes to the interactome of the protein. For this we used label-free quantitative LC-MS/MS of coimmunoprecipitation assays, for endogenous NDP52 from whole-cell HeLa extracts, with and without α-amanitin treatment (Fig.9E). Co-immunoprecipitation assays were performed as six replicates and compared to protein A controls. The same log_2_FC>1 and p_adj_<0.05 threshold was used to identify enriched GO processes. Interestingly, whilst in non-treated cells we can observe ‘gene expression’ as an enriched process, α-amanitin treatment disrupts this and appears to change NDP52 interactome to ‘regulation of DNA replication’, ‘signal transduction in response to DNA damage’, and ‘chromosome organisation’ (Supplementary Fig.11E and F). Figure 9F shows examples of top hits for endogenous NDP52 pull-downs in nontreated versus α-amanitin treated cells. The shift observed in interacting partners suggests that NDP52 preferentially interacts with different proteins, depending on cell state, and could change its interactome in response to environmental stresses. Interestingly, in nontreated cells we observe different GO enrichments to those observed in recombinant protein pull-down assays (Supplementary Fig.9A and Supplementary Fig.11E). The high concentration of recombinant NDP52-FL used could have allowed the identification of different interactions, that under normal conditions and cellular levels of NDP52 are less enriched.

Overall, our data show NDP52 as a novel transcription regulator, with functions in DNA structure.

## DISCUSSION

In this study, we used a multidisciplinary approach to shed light on the nuclear role of autophagy receptor NDP52. Although NDP52 was first observed in the nucleus ^1^, until now, no clear nuclear function had been attributed to this protein. By investigating its nuclear organisation, dynamics, interactome and biochemical characteristics, we have been able to link its function to transcription and DNA regulation.

To enhance their activity and functional efficiency, many nuclear proteins involved in transcription and other nuclear processes form molecular clusters ^17,18,29–31^. Here, we have determined that NDP52 clusters at regions of transcription initiation with RNAPII, and that its overexpression can increase the number of transcriptional clusters available in the nucleus. RNAPII clustering is directly related to transcription activity ^32–34^ and changes to its spatial organisation and clustering behaviour impact whole gene-expression levels ^35^. This provides a direct association between NDP52 and transcriptional regulation. Furthermore, we have also shown that knockdown of NDP52 impacts gene expression in both HeLa and MCF-7 cells. Altogether, these data support a role for NDP52 in RNAPII-related transcription.

Although future studies will be necessary to determine a mechanism for the regulatory role of NDP52 in transcription, here we propose two different strategies: i) through interactions with transcriptional machinery and regulatory factors at transcriptional sites and/or ii) through direct DNA structure regulation and interaction with chromatin remodellers (Fig. 10). In support of our first hypothesis, we show that NDP52 colocalises with RNAPII at transcription initiation sites and that clustering of NDP52 is abrogated following transcription inhibition. Although we observed more non-colocalised clusters of NDP52-Ser5 RNAPII than colocalised, it is important to note that not all Ser5-RNAPII represents actively transcribing complexes, and that part of this population is stalled/paused. Hence, it is possible that NDP52 preferentially localises with a subset of RNAPII, for example, with molecules that are going through initiation. Following transcription inhibition with α-amanitin, the observed increase in NDP52 molecular dynamics suggests a reduction in the available binding sites. These data indicate that loss of RNAPII molecules, due to its degradation, directly affects the nuclear organisation of NDP52. Furthermore, our proteomics data, both with recombinant and endogenous NDP52, also shows interactions with different transcriptional regulators. These interactions are markedly reduced in endogenous NDP52 pull-downs when cells are pre-treated with α-amanitin, which further supports the hypothesis that interactions between NDP52 and transcriptional regulators are an important part of the regulatory function of this protein.

**Figure 10:**
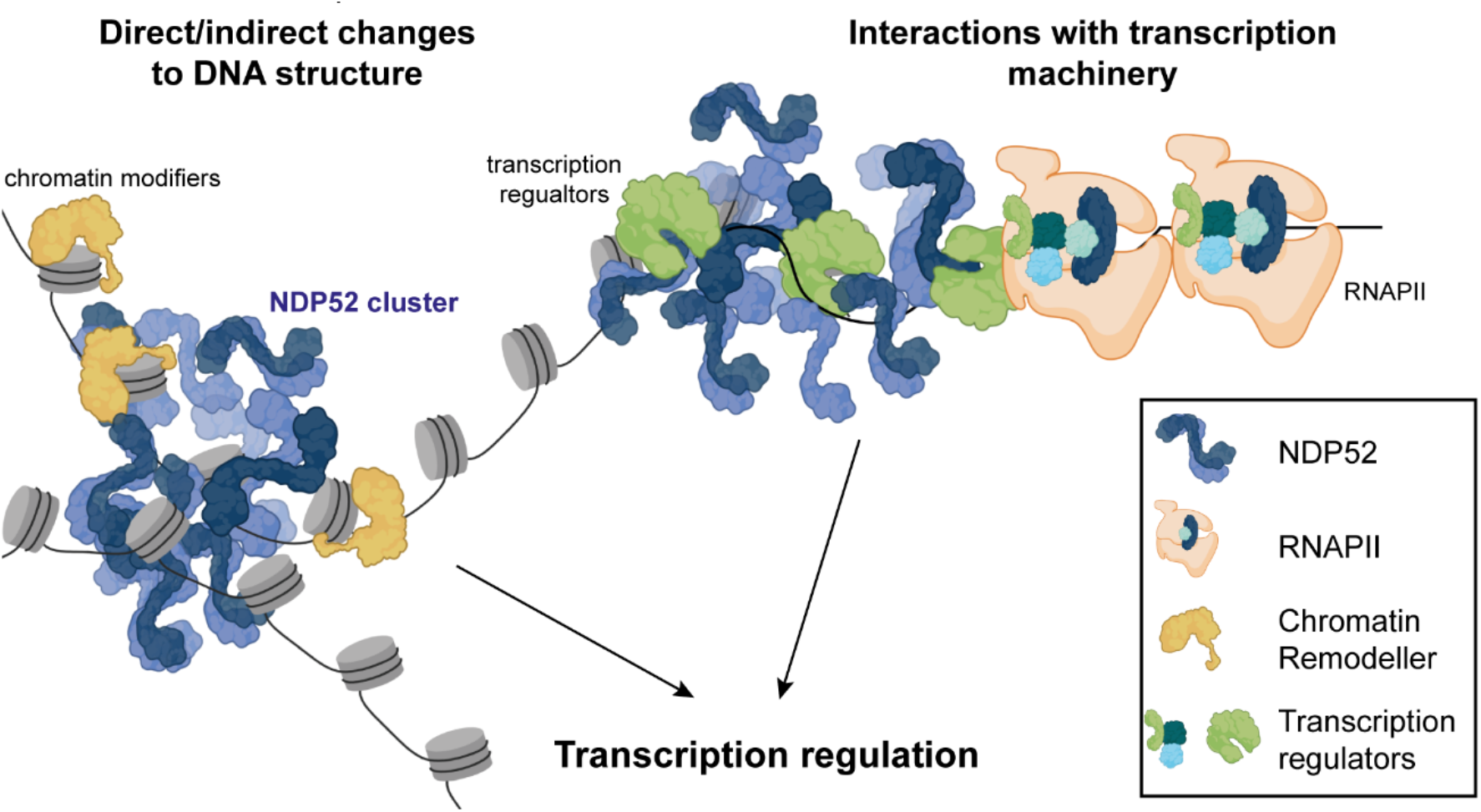
Model of possible mechanism for NDP52’s activity in transcription. NDP52 could directly interact with DNA in the nucleus, or with chromatin modifiers (*e.g*. histone modifiers), to cause local changes to chromatin structure. Conversely, interactions with transcription factors/coactivators and transcription machinery could also modulate transcription activity of genes.

Equally, we also show that NDP52 binds specifically and with high affinity to dsDNA and we believe this interaction to be crucial for its observed role in transcription. Importantly, we have shown that NDP52 can be isolated in complex with different genomic *loci*, through ChIP-qPCR, and this could be a regulatory strategy for the protein. In fact, CoCoA, a gene paralog of NDP52 and known transcription co-activator, has been found present at different genes regulated by nuclear receptors, such as the *TFF1/PS2* gene ^36^. Although our biochemical studies also show for the first time that CoCoA can directly bind DNA, previous studies have determined that interactions of this protein with histone methyltransferases and acetyltransferases at the gene body allows the recruitment of basal transcriptional machinery, thus promoting transcriptional activity ^11,36,37^. It will be interesting, in future studies, to produce a more comprehensive analysis of NDP52-genomic DNA interactions, through ChIP-Seq. There are also no available ChIP-Seq data for CoCoA. It would be informative to determine how similar the genomic targets of these two proteins are, given their biochemical likeness.

Interestingly, when exploring NDP52 binding to DNA, we observed that NDP52 promotes changes in DNA structure *in vitro* - through bending, bridging and DNA looping. This, together with our proteomics data showing regulators of chromatin and DNA structure as possible binding partners of NDP52, suggests a role in chromatin regulation for NDP52. Chromatin conformation and structure are important determinants of accessibility to transcriptional machinery ^38^; as a result, chromatin regulation is directly linked to transcriptional activity. Our data suggest that, in addition to direct regulation at transcriptional sites, either through direct interactions with RNAPII or other transcription factors, NDP52 activity on DNA structure and links to chromatin organisation could also drive transcriptional regulation. This could also explain how overexpression of NDP52 leads to an increase in RNAPII clusters, as changes to chromatin accessibility might occur. In future studies, it will be important to address specifically the role of NDP52 in chromatin structure and regulation and explore some of the possible new interactions with DNA binding proteins and regulators identified in our proteomics data.

Our work has also provided a detailed biochemical analysis of NDP52. We show that NDP52, in solution, is predominantly dimeric, as previously suggested ^6^; although it can also associate into higher oligomeric forms, such as trimers and tetramers. We also found that, in addition to its coiled-coil region, the C-terminal and N-terminal domains can also independently interact and form oligomeric structures. We also provide evidence that NDP52 interacts with DNA through its zinc finger domains and that both the full-length and C-terminal domains of NDP52 can oligomerise with and around DNA *in vitro*. Furthermore, we show that NDP52 can modify the local conformation of DNA, *in vitro*. In cells, this could provide a mechanism for a possible function of NDP52 in chromatin structure regulation, by increasing local concentrations of the protein around DNA. Moreover, given that the C-terminal domain is the main region for interactions with DNA, it is possible that, in the cell, NDP52 binds to DNA through this region whilst simultaneously sustaining interactions with other proteins through its N-terminal and coiled-coil regions. AFM data suggests a variety of spatial orientations might be available during oligomerisation of the protein, but further studies will be necessary.

Whilst the C-terminal of NDP52 is crucial for its DNA binding activity, our proteomics data also indicate that many important regulatory proteinprotein interactions also occur through this region. In fact, previous studies have identified the C-terminal domain of NDP52 as the main interacting region with Myosin VI and ubiquitin ^39^. Interestingly, the majority of common hits found in recombinant proteomics between full-length NDP52 and CoCoA arise from the C-terminal domain region of the protein. This is not surprising as this region displays high levels of amino acid homology in both proteins.

Interestingly, our proteomics data with endogenous NDP52 following treatment with α-amanitin, showed an enrichment for proteins involved in cell stress and DNA damage response. Although this was not explored in this study, it will be interesting to understand, in the future, how nuclear NDP52 responds to different cellular stresses. In the cytoplasm, NDP52 is known to be activated by certain cell stresses, namely in response to bacterial or viral infections or the presence of damaged organelles ^39–41^. It is not yet known how cell stress might affect nuclear levels, organisation or the nuclear activity of NDP52. It is possible that NDP52 has a dual cytoplasmic-nuclear role in cells and that its nuclear activity is linked to its cytoplasmic function in autophagy. Following cellular infection innate immunity and apoptotic pathways are activated in the cytoplasm that lead to the translocation of different proteins into the nucleus for their activity in transcription, chromatin and DNA repair regulation ^42–44^. Understanding the molecular role of NDP52 and its nuclear activity in context of its already known cytoplasmic function in autophagy will be important in future studies.

Overall, here, we provide evidence for NDP52 as a novel transcriptional regulator, with possible functions in chromatin structure and organisation.

## MATERIALS AND METHODS

### Constructs

A list of constructs is provided in Supplementary Table 1.

### Cell culture and Transfection

HeLa (ECACC 93021013) and MCF-7 (ECACC 86012803) cells were cultured in MEM Alpha medium (Gibco), with GlutaMax (no nucleosides), supplemented with 10% Fetal Bovine Serum (Gibco), 100 μg/mL streptomycin (Gibco) and 100 units/mL penicillin, at 37°C and 5% CO_2_. To inhibit transcription, cells were treated with 5μg/mL a-amanitin for 4 hours at 37°C and 5% CO_2_.

For transient transfection of Halo-NDP52, HeLa cells cultured in Nunc LabTek dishes (Merk) were transfected using Lipofectamine 2000 (Invitrogen) for 24h. Following this, cells were used for live-cell imaging using Fluorescence Recovery After Photobleaching (FRAP) or aberration-corrected Multi-Focal Microscopy (acMFM).

### Immunofluorescence

Following nuclear staining using Hoechst 33342 (Thermo Scientific), HeLa and MCF-7 cells cultured on glass coverslips were fixed for 15 minutes at room temperature, in 4%(w/v) paraformaldehyde (PFA) in Phosphate-buffered saline (PBS). Residual PFA was then quenched for 15 minutes using 50mM Ammonium Chloride in PBS.

Cells were permeabilised and blocked in 2% (w/v) Bovine Serum Albumin (BSA), 0.1% (v/v) Triton X-100 in PBS for 30 minutes. Cells were then labelled against endogenous proteins for 1 hour in 2% (w/v) BSA, with appropriate primary antibody and, subsequently, with appropriate fluorophore-conjugated secondary antibodies. When using anti-phospho antibodies, the immunofluorescence protocol was performed in Trisbuffered saline (TBS).

For endogenous NDP52, rabbit anti-NDP52 (1:200, Genetex GTX115378) antibodies were used. For RNA Polymerase II, mouse anti-RNAPII phospho Ser 5 (1:500, Abcam, ab5408) was used. Secondary conjugated antibodies Donkey anti-rabbit Alexa 647 (1:500, Abcam, 181347) and Donkey anti-mouse Alexa 488 (1:500, Abcam, ab181289) were used.

Coverslips were then mounted on microscope slides in Mowiol solution (10%(w/v) mowiol 4-88, 25%(v/v) glycerol, 0.2M Tris-HCl, pH 8.5) with 2.5%(w/v) DABCO (Sigma).

### Confocal Microscopy

Fixed cells were imaged using a Zeiss LSM980, with a Plan-Achromat 63 x 1.4 NA oil immersion objective (Carl Zeiss, 420782-9900-000). Three laser lines: 405, 488 and 561 were used for excitation of Hoechst, Alexa-fluor 488 and Alexa-fluor 647 fluorophores. Built-in multi-band dichroic mirror MBS405/488/561 (Carl Zeiss, 1784-995) were used to reflect excitation laser beams onto samples. For fluorescence signal collection, the used emission spectral bands were: 410–524 nm (Hoechst), 493–578 nm (Alexa-fluor 488) and 564–697 nm (Alexa-fluor 647). The green channel (Alexa-fluor 488) was imaged using a 1 gallium arsenide phosphide (GaAsP) detector, while the blue (Hoechst) and red (Alexa-fluor 647) channels were imaged using two multi-anode photomultiplier tubes (MA-PMTs). For imaging acquisition and rendering, Zeiss ZEN Blue software (v2.3) was used. Confocal Images were deconvolved using the Zeiss Zen Blue software (v2.3), using the regularized inverse filter method.

### Stochastic Optical Reconstruction Microscopy (STORM)

No. 1.5, 25mm round glass coverslips were cleaned by incubation with etch solution (5:1:1 ratio of H_2_O: H_2_O_2_ (50% wt in H_2_O stabilised, Fisher Scientific): NH_4_OH (ACS reagent, 29-30% NH3 basis, Sigma) for 2 hours in a 70°C water bath. Cleaned coverslips were washed in filtered water and ethanol and allowed to dry before cell seeding.

Cells were fixed for 15 minutes in 4%(w/v) PFA in PBS and residual PFA was quenched with 50mM Ammonium Chloride in PBS for 15 minutes. Immunofluorescence was performed using filtered TBS. Cells were first permeabilised and blocked for 30 minutes in 3% (w/v) BSA, 0.1% (v/v) Triton X-100. Cells were then incubated in primary antibody for 1 hour, at the same dilution as for the normal immunofluorescence protocol. Cells were washed three times (10 minutes each wash) with 0.2% (w/v) BSA, 0.05% (v/v) Triton X-100 in TBS. Cells were subsequently incubated in an appropriate fluorophore-conjugated secondary antibody for 1 hour, at a 1:250 dilution, in 3% (w/v) BSA, 0.1 % (v/v) Triton X-100. Cells were washed in TBS and PBS and fixed in 4% (w/v) PFA in PBS a second time. Cells were stored in PBS supplemented with 0.02% (w/v) NaN3 in the dark until imaging.

Before imaging, coverslips were washed in filtered H_2_O and assembled into Attofluor cell chambers (Invitrogen). Imaging was performed in STORM buffer - 10% (w/v) glucose, 10mM NaCl, 50mM Tris-HCl pH8.0 - supplemented with GLOX solution (5.6% (w/v) glucose oxidase and 3.4 mg/mL catalase in 50mM NaCl, 10mM Tris-HCl pH 8.0) and 0.1% (v/v) 2-mercaptoethanol.

STORM imaging was performed using a Zeiss Elyra PS.1 system. For sample illumination HR Diode 488 nm (100mW) and HR Diode 642 nm (150mW) lasers were used, where power density on the sample was 7-12kW/cm^2^ and 7-14kW/cm^2^, respectively. Built-in multi-band dichroic mirror MBS 405/488/642 (Carl Zeiss 1784-996) were used to reflect excitation laser beams onto samples. To reduce background fluorescence levels, imaging was performed using Highly Inclined and Laminated (HILO) illumination with a 100x NA 1.46 oil immersion objective (Carl Zeiss alpha Plan-Apochromat, 420792-9800-000). For fluorescence signal collection, a BP 420-480/BP 495-550/LP 650 multi-bandpass emission filter (Carl Zeiss 1769-207) was used and a final image was acquired using an Andor iXon DU 897 EMCCD camera with 25msec exposure, for 25000 frames.

Image processing was performed in Zeiss Zen Black software. For two-colour STORM images, channel alignment was performed following a calibration procedure using pre-mounted MultiSpec beads (Carl Zeiss, 2076-515). For the calibration procedure, the affine method was performed, to account for lateral stretching and tilting between the two channels. This was performed for each day of acquisitions. Blinking event detection was then performed in Zeiss Zen Black software using a 9-pixel mask with a signal-to-noise ratio of 6, accounting for overlap, to allow localisation of molecules in dense environments. Final molecule positions were then determined through fitting of a 2D Gaussian function. Molecule positions were subjected to model-based cross-correlation drift correction. For Alexa-fluor-647 and Alexa-fluor-488 labelled molecules, the typical mean value of localisation precision was 20 nm and 30 nm, respectively. Molecule localisation tables were exported as .txt files for further analysis using ClusDoC software ^22^.

### ClusDoC

Following export of molecule positions from Zeiss ZEN Black software, STORM data were further analysed using ClusDoC software (https://github.com/PRNicovich/ClusDoC) ^22^. The nucleus was selected as an ROI for cluster analysis. The Ripley K function was first calculated for the ROI selected, to identify the r max. This value was then used in DBSCAN analysis for single-channel images or ClusDoC analysis for two-channel images. Minimum cluster size was set to 5 molecules with a smoothing value set at 7 and an epsilon value set at the mean localisation precision value for the dye. Other parameters remained at default values.

### Aberration Corrected Multi-Focal Microscopy

Cells transiently expressing Halo-NDP52 were labelled for 15 minutes with 10 nM Halo tag-JF549 ligand in cell culture medium at 37°C, 5% CO_2_. Cells were then washed three times in complete media and incubated for at least 30 minutes at 37°C, 5% CO_2_ before imaging. For imaging, cell media was replaced with FluoroBrite DMEM medium (Thermo Fisher Scientific). Single-molecule tracking experiments were performed using an aberration-corrected multi-focal microscope (acMFM), described in Abrahamsson *et al*., ^45^. Briefly, a custom optical system appended to the detection path of an optical Nikon Ti microscope was used. The detection path of the microscope included a diffractive multifocal grating in a conjugate pupil plane, a chromatic correction grating, to reverse spectral dispersion, and a nine-faceted prism followed by the final imaging lens. A 561 nm laser was used for excitation, with a 4-6 kW/cm^2^ power density at the back aperture of a 100x 1.4NA objective (Nikon).

AcMFM imaging produces nine separate, simultaneous images, each representing a separate focal plane, with an axial separation of ca. 400 nm between them. Field of view is ca. 20 μm. The nine-image array was digitised via an EMCCD camera (iXon Du897, Andor), at up to 32msec temporal resolution, with a typical duration of 30 seconds.

3D+t single-molecule images were reconstructed via a calibration procedure in Matlab (MathWorks) that accounts and calculates (1) inter-plane spacing, (2) affine transformation for the correct alignment of each focal plane in xy and (3) slight variations in the detection efficiency in each plane - typically less than 5-15% from the mean.

Reconstructed data were pre-processed, including background subtraction and deconvolution (3-5 Richardson-Lucy iterations) and/or Gaussian denoising prior to 3D particle tracking using the MOSAIC software suite. Maximum particle displacement was set at 400 nm and a minimum 10 frames was required. Detected tracks were reconstructed and diffusion constants calculated through MSD analysis using custom Matlab software assuming an anomalous diffusion model.

### Fluorescence Recovery After Photobleaching

Cells transiently expressing Halo-NDP52 were labelled for 15 minutes with 10 nM Halo tag-JF549 ligand in cell culture medium at 37°C, 5% CO_2_. Cells were then washed three times in CO_2_-independent medium (ThermoFisher) before imaging.

FRAP measurement was performed using a Zeiss LSM 880 system equipped with a 100x NA 1.46 oil immersion objective (Carl Zeiss alpha Plan-Apochromat, 420792-9800-000). For sample illumination a 20 mW 561 nm diode laser was used. Built-in multi-band dichroic mirror MBS 458/561 was used to reflect excitation laser beams onto samples. For fluorescence signal collection, the wavelengths from 566 nm to 685 nm were captured using a multianode photomultiplier tube (MA-PMT) with 0.96 μs pixel dwell time. The detector master gain was 900, and digital gain was 1.

Ten frames of confocal microscopy image under 8 mW 561 nm laser illumination were acquired before photobleaching. Selected regions of interest (ROIs) were exposed to full laser power, followed by 100 seconds of confocal microscopy image acquisition. The time course of fluorescence intensity from the selected ROIs was recorded by Zeiss ZEN 2.3 Blue software. Fluorescence intensity time traces from ROIs, whole cell areas and background areas were exported as .txt files, and then were analysed using easyFRAP Software (https://easyfrap.vmnet.upatras.gr/?AspxAutoDetectCookieSupport=1) ^46^.

### RNA-Sequencing and Analysis

Total RNA was extracted from three replicates of NDP52 KD (using CALCOCO2 siRNA, Ambion, 4392420) and scrambled siRNA (using control siRNA, Qiagen, 1027280) in MCF-7 and HeLa cells. Ice cold TRIzol reagent was added to each cell culture dish and homogenised. The mixture was incubated at room temperature for 10 minutes. Chloroform was then added to the mixture and incubated at room temperature for 5 minutes. The samples were centrifuged at 8,000 *xg*. The top, aqueous layer was collected and isopropanol was added. The mixture was then centrifuged at 12, 000 *xg* for 30 minutes and the supernatant discarded. The pellet was washed in 75% (v/v) ethanol and centrifuged at 7, 500 *xg* for 5 minutes. The pellet was air dried and resuspended in RNAse-free H_2_O. RNA concentration and quality was then assessed by measuring absorbance at 260 nm and A260/A280 ratio. RNA samples were stored at −80°C. The following procedures were performed by GENEWIZ and Glasgow Polyomics. The RNA-seq libraries were prepared using Poly-A selection. Resulting libraries concentration, size distribution and quality were assessed on a Qubit fluorometer and on an Agilent 2100 bioanalyzer. Paired-end sequencing (2×150 bp) was then performed on an Illumina NovaSeq next generation sequencer for HeLa cells (GENEWIZ) and (2×75 bp) on a HiSeq sequencer for MCF7 cells (Glasgow Polyomics).

Sequence reads were trimmed to remove possible adapter sequences and nucleotides with poor quality using Trimmomatic v.0.36. The trimmed reads were mapped to the Homo sapiens GRCh38 reference genome available on ENSEMBL using the STAR aligner v.2.5.2b. BAM files were generated as a result of this step. Unique gene hit counts were calculated by using featureCounts from the Subread package v.1.5.2. After extraction of gene hit counts, the gene hit counts table was used for downstream differential expression analysis. Using DESeq2, a comparison of gene expression between the customer-defined groups of samples was performed. The Wald test was used to generate p-values and log_2_ fold changes. Genes with an adjusted p-value < 0.05 and absolute log_2_ fold change > 1 were called as differentially expressed genes for each comparison.

Differentially expressed genes by at least 1.5-fold (−0.5≥ log_2_FC ≥0.5) and adjust p-value <0.05 were subjected to Gene Ontology analysis, using iDEP93 (http://bioinformatics.sdstate.edu/idep93/) ^47^. RNA-Seq data were deposited in the Gene Expression Omnibus (GEO) database under the accession number GSE188567.

### Chromatin Immunoprecipitation and qPCR

To identify NDP52-DNA interactions, ChIP was performed using anti-rabbit NDP52 antibody (Genetex GTX115378). HeLa cells were crosslinked by adding formaldehyde directly to the cell medium, to a final concentration of 0.75% (v/v). Cells were left to incubate with gentle rotation at room temperature for 10 minutes. The reaction was stopped by adding glycine to a final concentration of 125 mM and incubating the mixture for 5 minutes at room temperature with rotation. Cells were washed twice with cold PBS and scraped in cold PBS. All cells were collected by centrifugation at 1, 000 *xg* at 4°C for 5 minutes. The pellet was resuspended in ChIP lysis buffer - 50 mM HEPES-KOH pH 7.5, 140 mM NaCl, 1 mM EDTA pH 8.0, 1% (v/v) Triton X-100, 0.1% (m/v) Sodium Deoxycholate, 0.1% (m/v) SDS - supplemented with protease inhibitors, using 750 μL per 1×10^7^ cells. Cells were sonicated using a diogenode bioruptor sonicator to shear DNA until an average DNA fragment size of 200-800bp was achieved. Fragment size was determined using a 1.5% agarose gel. Cell debris were removed through centrifugation of samples at 8, 000 *xg* for 10 minutes at 4°C.The supernatant, enriched for chromatin, was stored at −80°C until used for immunoprecipitation experiments.

Chromatin fractions were diluted 1:10 in RIPA buffer - 50 mM Tris-HCl pH 8.0, 150 mM NaCl, 2 mM EDTA pH 8.0, 1% (v/v) NP40, 0.5% (m/v) Sodium Deoxycholate, 0.1% (m/v) SDS - supplemented with protease inhibitors. Three samples were used for immunoprecipitation with NDP52 and three samples for no-antibody control (beads only). 10 % of total chromatin was removed as input sample and stored at −20°C. All samples were ple-cleared using protein A magnetic beads (Thermo Fisher Scientific) for 30 minutes at 4°C with end-to-end rotation. Immunoprecipitation replicates were incubated overnight with NDP52 antibody (1:50 dilution) at 4°C with end-to-end rotation. The following day, 40 μL of protein A magnetic beads, pre-equilibrated in RIPA buffer, were added to each sample, including the no-antibody controls. Samples were incubated with end-to-end rotation at 4°C for 1 hour. Following this, beads were collected using a magnetic rack and washed twice in low-salt buffer (20 mM Tris-HCl pH 8.0, 150 mM NaCl, 0.1% (m/v) SDS, 1% (v/v) Triton X-100, 2 mM EDTA) followed by a wash in high-salt (20 mM Tris-HCl pH 8.0, 500 mM NaCl, 0.1% (m/v) SDS, 1% (v/v) Triton X-100, 2 mM EDTA), a wash in LiCl buffer (10 mM Tris-HCl pH 8.0, 250 mM LiCl, 1% (m/v) Sodium deoxycholate, 1% (v/v) NP40, 1 mM EDTA pH 8.0) and, finally, in TE buffer (10 mM Tris-HCl pH 8.0; 1 mM EDTA pH 8.0).

DNA was eluted by incubating the beads with 120 μL elution buffer (1%(w/v) SDS; 100mM NaHCO_3_) at 30°C, with shaking. To reverse crosslinking, eluted protein-DNA complexes and input samples were incubated overnight with 4.8 μL NaCl (5M) and 2 μL RNAse A (10mg/mL) at 65°C with shaking. The following day, samples were incubated with Proteinase K for 1 hour at 60°C. The DNA was purified using phenol:chloroform extraction and samples analysed using QuantiNova SYBR Green qPCR kit (Qiagen). A list of qPCR primers for ChIP is supplied in supplementary Table 2.

### Protein expression and purification in Escherichia coli

Recombinant protein expression was performed in *E. coli* BL21 DE3 cell (Invitrogen) in Luria Bertani media. Proteins were purified by affinity chromatography, using HisTrap FF columns (GE Healthcare). Protein fractions were further purified using Size Exclusion Chromatography, using a Superdex 200 16/600 column (GE Healthcare).

### Size Exclusion Chromatography and Multi-Angle Light Scattering

100 μL samples of recombinant proteins, at concentrations of 1mg/mL (NDP52-FL and CoCoA) and 5mg/mL (_N_NDP52, _C_NDP52 and ZF2) were loaded onto a Superdex 200 (30 x 1cm) analytical gel filtration column (GE Healthcare), equilibrated in 50mM Tris pH7.5, 150 mM NaCl, 1mM DTT and controlled by Waters 626 HPLC at room temperature. A Viskotek SEC-MALS 9 and Viscotek RI detector VE3580 (Malvern Panalytical) were used for detection. Analysis was performed using Omnisec software (Malvern Panalytical).

### Dynamic Light Scattering

Dynamic light scattering measurements were performed at 20°C, using a Zetasizer Nano ZS DLS system (Malvern Panalytical). Before measuring light scattering intensity at 90° angle, samples were centrifuged at 20 000*xg* for 10 minutes. Analysis was undertaken using the Zetasizer software.

### Microscale Thermophoresis

Recombinantly purified protein constructs were labelled with RED-tris NTA dye (NanoTemper Technologies GmbH) in PBS to a concentration of 100 nM. A 20μM stock of non-labelled protein was also prepared. This stock was used in a 16-step serial dilution in PBS buffer. For oligomerisation studies,10μM of protein was used as the highest ligand concentration, with Red-tris-NTA labelled protein kept at a final concentration of 50 nM for all reactions.

Reactions were incubated for 15 minutes at room temperature in the dark and loaded into Monolith NT.115 Capillaries (NanoTemper Technologies GmbH). Microscale thermophoresis measurements were performed using a Monolith NT.115 (NanoTemper Technologies GmbH) with 20%(RED) LED and high MST power. Binding assays were run as three independent experiments and the data were fitted using a *K_D_* model with ligand-induced initial fluorescence change, as described by Jerabek-Willemsen *et al*. ^48^.

### Circular Dichroism

Recombinantly purified constructs were prepared in 50mM Tris-HCl pH7.5, 150 mM NaCl. Circular dichroism spectra were obtained from 200 μL samples in a 1-mm cuvette, in a J175 spectropolarimeter from Jasco, with data collected at 0.5 nm intervals with averaging of 16 scans. For thermostability data, spectra were collected between 20 and 90 °C, and mean residue ellipticity values at 222 nm or 215 nm wavelength were fitted to a simple sigmoidal curve.

### Electrophoretic Mobility Shift Assay

Reactions were performed in a final volume of 30 μL, with 250 nM ds40 and increasing concentrations of NDP52-FL (between 0.5 and 3μM) in 50 mM Tris-HCl pH 7.5, 50 mM NaCl and 3 mM MgCl_2_. After 5 minutes incubation, reactions were supplemented with 3μL of 30% (v/v) glycerol, loaded on a 3% agarose gel and run in Tris-Borate buffer at 60V. Gels were incubated for 30 minutes in ethidium bromide, washed for 20 minutes with H_2_O and visualised under UV light.

### Nano-Differential Scanning Fluorimetry

Thermostability of recombinantly produced proteins NDP52-FL, _C_NDP52 and ZF2 was assessed using nano-DSF, at protein concentrations 50 μM, 75 μM and 100 μM, respectively. Samples were prepared in 50 mM Tris-HCl pH 7.5, 150 mM NaCl and loaded in nanoDSF Grade Standard Capillaries (NanoTemper Technologies GmbH), for NDP52-FL, or nanoDSF Grade High Sensitivity Capillaries (NanoTemper Technologies GmbH), for _C_NDP52 and ZF2. Data were acquired using a Prometheus NT.48 (NanoTemper Technologies GmbH). Thermal denaturation of proteins was detected with heating in a linear thermal ramp (2°C/min^-1^) between 20 and 90°C, with an excitation power of 60-90%. Temperature unfolding was detected by following fluorescence emission at 350 and 330 nm wavelength. Melting temperatures were determined as the maxima of the first derivative of the ratio 350nm/330nm, using NanoTemper software (NanoTemper Technologies GmbH).

### Mass Photometry

Before measurements, samples were centrifuged at 20 000 xg for 10 min. The samples were then diluted to 10 nM immediately prior to measurements in 50mM Tris-HCl pH7.5, 150 mM NaCl. Measurements were performed on clean glass coverslips and recorded on the OneMP mass photometer (Refeyn Ltd) for 60 s. Each measurement was repeated at least 3 times. The recorded videos were analyzed using DiscoverMP (Refeyn Ltd) to quantify protein binding events. The molecular weight was obtained by contrast comparison with known mass standards (BSA, Urease and IgG) measured on the same day.

### Sample preparation for AFM

Preparation of DNA and protein samples for imaging was carried out as described fully in a published protocol^49^. Linear 339 bp DNA molecules (ds339), NDP52-FL and _C_NDP52 were adsorbed onto freshly cleaved mica disks (diameter 5 mm, Agar Scientific, UK), separately and in combination, at room temperature using poly-L-lysine (PLL) ^49^. Briefly, 20 μL PLL 0.01% solution, MW 150,000–300,00 (Sigma-Aldrich) was pipetted onto freshly cleaved mica to adsorb for 1 min. The PLL coated surface was washed in a stream of MilliQ^®^ ultrapure water, resistivity >18.2 MΩ to remove non-adsorbed PLL. To immobilise DNA and DNA-protein complexes 20 μL of 50 mM TRIS, 150 mM NaCl buffer solution was pipetted onto the PLL coated mica and the following masses of sample added: 7ng 339 bp linear DNA; 10 ng NDP52 with 6 ng 339 bp linear DNA; and 10 ng of _c_NDP52 with 14 ng 339 bp linear DNA. To immobilise NDP52-FL and _c_NDP52 alone, 20 μL of 50 mM TRIS, 150 mM NaCl was pipetted onto the PLL coated mica and 3 - 8 ng of protein added to the buffer solution. All samples were adsorbed for 10 minutes followed by four washes in the same buffer to remove unbound DNA or protein. NDP52-FL was also adsorbed onto freshly cleaved mica using Ni^2+^ ions. Here 30 μL of 50 mM TRIS, 150 mM NaCl, 2 mM NiCl_2_ was placed onto the mica and between 3.5 and 7 ng of NDP52-FL was added and the sample incubated for 30 minutes before washing four times in the same buffer as before.

### AFM Imaging

All AFM measurements were performed in liquid following a published protocol ^49^. Experiments were carried out in PeakForce Tapping mode on a FastScan Dimension XR AFM (Bruker) using FastScan D AFM probes (Bruker). Continuous force–distance curves were recorded over 40 nm (PeakForce Tapping amplitude of 20 nm), at a frequency of 8 kHz with the tip-sample feedback set by PeakForce setpoints in the range of 5–12 mV as referenced from the force baseline resulting in peak forces of 40–100 pN. Images were recorded at 512 × 512 pixels to ensure a resolution ≥ 1 nm/pixel at line rates of 1–4 Hz.

### AFM image processing

TopoStats ^50^, a package of python scripts, was used to automate the processing of the AFM data and the analysis of the DNA and protein molecules. The code is freely available at https://github.com/AFM-SPM/TopoStats. Raw AFM data were processed using the ‘pygwytracing.py’ script which utilises the Gwyddion ‘pygwy’ module for automated image correction, molecule identification and morphological analysis.

Basic image processing is performed by the ‘editfile’ function, using various Gwyddion processes to align and level data as well as correcting imaging artefacts. A gaussian filter (*σ*=1) of 1.5 pixels (1–2nm) was applied to remove pixel errors and high-frequency noise.

Protein molecules are identified in the images of NDP52-FL and _C_NDP52 using a combination of Gwyddion’s automated masking protocols so that masks define the positions of individual molecules in the images. These masked molecules within a flattened AFM image are identified using the ‘mask_outliers’ function, which masks data points with height values that deviate from the mean by a customisable multiplier of σ (with 3σ corresponding to a standard gaussian). This multiplier was optimised to select features based on their height in order to correctly mask the protein molecules and oligomers, and the values were 0.7σ for _C_NDP52 and 0.57σ for NDP52-FL. Example masked data created using these parameters is shown in Supplementary Fig.3K. The Gwyddion functions ‘grains_remove_touching_border’ and ‘grains_remove_by_size’ are used to remove molecules that are cut off by the edge of the image and those that are < 50 nm^2^ respectively. Large aggregates and small contaminants are removed using the ‘removelargeobjects’ and ‘removesmallobjects’ functions respectively, which use the function ‘find_median_pixel_area’ to determine the median size of the masked molecules then removes objects outside of a customisable size range based on this. The morphological properties of individual masked molecules are calculated for each image using the ‘grainanalysis’ function, which uses the Gwyddion function ‘grains_get_values’ to obtain statistical properties, including the length and width of the masked molecules. These are referred to in Gwyddion as ‘GRAIN_VALUE_MINIMUM_BOUND_SIZE’ and ‘GRAIN_VALUE_MAXIMUM_BOUND_SIZE’ respectively, and measure the maximum and minimum bounding sizes of the 2D mask for each molecule.

Each grain’s values are appended to an array [appended_data] for morphological analysis of individual molecules from all images in a single directory. This array is converted to a pandas dataframe ^51^ using the ‘getdataforallfiles’ function and saved out using the ‘savestats’ function as ‘.json’ and ‘.csv’ files with the name of the directory in the original path.

Statistical analysis and plotting are performed using the ‘Plotting.py’ script. This script uses the ‘importfromjson’ function to import the .json file exported by ‘savestats’ in pygwytracing.py and calculates various morphological properties from the masked molecules, including the maximum and minimum bounding size. Distributions are generated and plotted for the maximum and minimum bound sizes using the matplotlib ^52^ and seaborn libraries within the function ‘plotdist2var’ (https://zenodo.org/record/12710#.YfhFffXP1lY).

The binding position preference for NDP52-FL on ds339 was measured manually by loading the processed images into ImageJ ^53^ and using the measurement tools to determine the distance from the nearest end that the protein binding occurred.

### DNA binding Assays

FITC-labelled and unlabelled oligonucleotides were purchased from Sigma-Aldrich. For dsDNA preparation, equimolar concentrations of complementary ssDNA oligonucleotides were mixed ^54^. A list of oligonucleotides is provided in Supplementary Table 3.

Recombinant NDP52 (NDP52-FL, _C_NDP52, _N_NDP52 and ZF2) and CoCoA constructs were titrated into 100nM of FITC labelled ssDNA 40 bp or dsDNA, 15 or 40 bp, in 50mM Tris-HCl pH7.5, 150mM NaCl, 1mM DTT. Measurements were performed using a ClarioStar microplate reader (BMG Labtech). Fluorescence excitation was performed at 495 nm wavelength and emission spectra were measured between 515 and 570 nm wavelength, with fluorescence intensity values taken at 520 nm. Change in fluorescence was plotted in function of protein concentration, using three independent replicates for each experiment. Titration curves for NDP52-FL, CoCoA and _N_NDP52 were fitted to a binding quadratic equation:

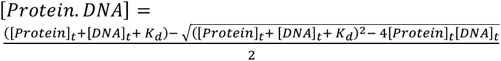

For _C_NDP52 and ZF2, a modified quadratic equation, accounting for a linear portion of the curve was used:

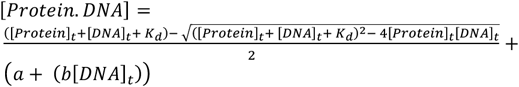

### Small-Angle X-Ray Scattering and Ab initio envelope calculation

Recombinantly expressed and purified NDP52-FL in SAXS buffer (50mM Tris-HCl pH7.5, 150mM NaCl, 1 mM DTT) was used for SEC-SAXS experiments at a concentration of 5mg/mL. NDP52-FL was analysed using a Superdex 200 increase 3.2/300 column, at a flow rate of 0.075mL/min (Cytiva Life Sciences), using an Agilent 1200 HPLC system (Agilent LC). SEC-SAXS experiments were performed at the B21 Beamline, Diamond Light Source UK, by core facility staff. For SEC-SAXS analysis and envelope generation, ScÅtter software (Version J) was used in combination with the ATSAS package ^55^.

### Nuclear isolation and extract preparation

Nuclear isolation was performed as previously described and characterised ^15,56,57^. Briefly, HeLa cells were collected and washed once with ice-cold PBS, then washed with ice cold Hypotonic buffer N (10 mM Hepes pH 7.5, 2 mM MgCl_2_, 25 mM KCl supplemented with 1 mM PMSF, 1 mM DTT and Protease Inhibitors (Thermo Fisher Scientific)). Cells were resuspended in cold Hypotonic buffer N and incubated on ice for 1h. Following this, cells were homogenised on ice using a glass Dounce homogeniser (Wheaton) and cell lysate was supplemented with sucrose solution to a final concentration of 220 mM, before centrifugation. The pellet, corresponding to isolated nuclei, was washed with cold Buffer N (10 mM Hepes pH 7.5, 2 mM MgCl2, 25 mM KCl, 250mM sucrose, 1 mM PMSF, 1 mM DTT, supplemented with Protease Inhibitor Cocktail.

For nuclear extract preparation, nuclei were incubated in nuclear ‘Hypotonic lysis buffer’ (10 mM Hepes pH 7.5, 2mM MgCl2, 25 mM KCl, 0.1%(V/V) Triton X-100, 0.1%(V/V) NP-40, supplemented with 1 mM PMSF, 1 mM DTT and Protease Inhibitors) for 1 hour on ice. Lysed nuclei were then used for recombinant protein pull-downs.

### Immunoblotting

Cell pellets from HeLa and MCF-7 cells, following NDP52 KD or control siRNA, were heat-denatured and resolved by SDS-PAGE. Membranes were probed against actin (Abcam, ab6276) and NDP52 by incubation with rabbit polyclonal primary antibody (1:2000 dilution, GeneTex, GTX115378) and, subsequently, a goat anti-rabbit antibody, coupled to horseradish peroxidase (1:15 000 dilution, Abcam, ab6721). Bands were visualised with ECL Western Blotting detection reagents (Invitrogen) using a ChemiDoc gel imager (Bio-Rad). For Ponceau S staining, membranes were incubated for 5 minutes in Ponceau S reagent (Sigma), washed three times with water and then imaged.

### Co-immunoprecipitation

HeLa cells (non-treated or following α-amanitin treatment for 4 hours) were collected and centrifuged at 500*xg* for 5 minutes at 4 °C. 1×10^6^ cells were used per co-immunoprecipitation assay. Each pellet was resuspended in 200 μL of Lysis buffer (10mM Hepes pH7.5, 2 mM MgCl_2_, 25mM KCl, 0.1mM DTT, 0.01mM PMSF, 0.1%(V/V) Triton X-100, 0.1%(V/V) NP40 and supplemented with protease inhibitors). Cells were left in Lysis buffer, on ice, for 1 hour. All samples were ple-cleared using protein A magnetic beads (Thermo Fisher Scientific) for 30 minutes at 4°C with end-to-end rotation. Immunoprecipitation replicates were incubated overnight with NDP52 antibody (1:100 dilution) at 4°C with end-to-end rotation. The following day, 50 μL of protein A magnetic beads, preequilibrated in Lysis buffer, were added to each sample, including the no-antibody controls, and incubated at 4°C with end-to-end rotation for 2 hours. Following this, beads were collected using a magnetic rack and washed three times with PBS. After removing all PBS, 50μL of loading buffer (NuPAGE LDS sample buffer supplemented with 50 mM DTT) were added and samples were incubated at 95°C for 10 minutes. Samples were then loaded on SDS-PAGE gels for ingel protein digestion for LC-MS/MS.

### Recombinant protein pull-downs

Following recombinant protein purification, 2.5mg of protein, per replicate, were incubated with nuclear extract (2×10^6^ nuclei per pull-down) at 4°C with end-to-end rotation, for 4 hours. Following this, 100 μL of Ni^2+^ magnetic beads (HisPur Ni-NTA ThermoFisher), preequilibrated in equilibration buffer (50mM Tris-HCl pH 7.5, 150 mM NaCl, 20 mM imidazole, 1mM DTT), were added to each sample, including the no-protein controls, and incubated at 4°C with end-to-end rotation for 2 hours. Following this, beads were collected using a magnetic rack and washed three times with ‘low imidazole buffer’ (50mM Tris-HCl pH 7.5, 150 mM NaCl, 40 mM imidazole, 1mM DTT). Following this step, three elutions were performed using ‘high imidazole buffer’ (50mM Tris-HCl pH 7.5, 150 mM NaCl, 400 mM imidazole, 1mM DTT). Eluted samples were loaded on SDS-PAGE gels for in-gel protein digestion for LC-MS/MS.

### In-gel digestion and LC-MS/MS

Following co-immunoprecipitation or recombinant protein pull-downs, samples were run only within the stacking portion of SDS-PAGE gels. Following this, gels were then stained, leaving a single band in the stacking portion of the gel, with all the protein content of each sample. Gel bands for each replicate were extracted, cut into 1×1mm squares and transferred into clean 1.5mL tubes. Gel particles were incubated with 50mM ammonium bicarbonate and acetonitrile in a 1:1 ratio at room temperature for 15 minutes and then centrifuged at 8, 000 *xg* for 60 seconds and the supernatant discarded. Samples were then incubated in acetonitrile for 15 minutes and centrifuged to remove supernatant. Gel particles were then incubated in 10mM DTT, 50mM ammonium bicarbonate and incubated at 56°C for 30 mins. Following this, the samples were centrifuged, the supernatant removed, and samples briefly incubated in acetonitrile until gel pieces shrunk. Samples were centrifuged and 55mM iodoacetamide in 50mM ammonium bicarbonate solution was added so that all gel particles were submerged. Samples were incubated in the dark, at room temperature, for 20 minutes. Following centrifugation, the supernatant was removed and washed in 50mM ammonium bicarbonate solution:acetonitrile (1:1) for 15 minutes, followed by 50mM ammonium bicarbonate for 15 minutes and acetonitrile for 15 minutes (between each step samples were centrifuged and supernatant discarded). Samples were then centrifuged and all liquid removed.

For tryptic digestion, gel particles were incubated in digestion buffer (25 mM ammonium bicarbonate, 10%(V/V) acetonitrile and 10ng/μL Trypsin (Sigma Aldrich, EMS0006)) for 30 minutes on ice. Digestion buffer was replenished as needed during this process to ensure gel particles were covered in solution. After 30 minutes, excess digestion buffer was removed from each sample and replaced with 25mM ammonium bicarbonate, 10%(V/V) acetonitrile solution. Samples were incubated overnight at room temperature.

The next day, acetonitrile was added to each tube and samples were sonicated in an ultrasound bath for 15 minutes. Samples were then centrifuged and the supernatant, containing digested protein for mass spectrometry, was transferred into clean 1.5mL tubes. 50%(v/v) acetonitrile and 5%(v/v) formic acid solution was added to gel particles and these were sonicated again. The supernatant from this step was combined with the previously collected sample into the same tube. Extracted protein samples were vacuum dried and resuspended in 10%(v/v) acetonitrile, 0.1%(v/v) trifluoroacetic acid for nanoLC-MS.

Peptides were separated on a HSS T3 Acquity column (Waters) 75 μm i.d. x 15 cm (1.8 μm, 100A) using an Acquity M-Class UPLC (Waters), elution was performed with a linear gradient from 3 to 40% B over 40 mins (solvent A = 0.1% formic acid, solvent B = 0.1% formic acid, acetonitrile) and the eluate directed via a nanospray source to a Synapt G2-Si (Waters) with data collected in UDMSe mode. Mass spectrometry data were imported into the software package Progenesis QI (Non-Linear Dynamics) and searched against a protein database using an MSe Search algorithm with an FDR set to 4%. Progenesis QI software (Waters) provided quality control information and quantification of peptides. The peptides were assigned using the ‘human proteome including enolase v5 2017’ from UNIPROT as a reference library, accounting for trypsin cleavage, carbamidomethyl modifications to cysteine residues and methionine oxidation. Maximum protein mass was set to 500kDa with a maximum of one missed cleavage allowed.

For peptide and protein assignments, a minimum of 3 fragments per peptide was required and a minimum of 5 fragments per protein. All assigned proteins contained at least one unique peptide. Following PCA analysis, replicates that didn’t cluster were excluded. Hits with a log_2_FC > 1 and ANOVA p<0.05, compared to controls (protein A or Ni^2+^ magnetic beads-pull downs) were considered for further analysis. Protein hits were submitted to Gene Ontology analysis using Gene Ontology Resource (https://geneontology.org).

The mass spectrometry proteomics data have been deposited to the ProteomeXchange Consortium via the PRIDE ^58^ partner repository with the data identifier PXD030238.

### Electron Microscopy

Cells grown on Aclar membrane (Agar Scientific) were fixed in 2% (w/v) formaldehyde and 0.5% glutaraldehyde in CAB (100mM Sodium cacodylate buffer pH7.2) for 2 hours at Room Temperature. The sample was washed 2 x10 minutes in CAB. Cells were dehydrated by incubation in an ethanol gradient, 50% ethanol for 10 min, 70% ethanol overnight, and 90% ethanol for 10 min followed by three 10-min washes in 100% dry ethanol. Cells were then suspended in LR White resin medium grade (London Resin Company) for 4h and then in fresh LR White resin overnight. Following 2 x 4-hour changes in fresh LR White resin samples were placed in sealed gelatine capsules and were polymerised upright at 60°C for 20 hours. Ultrathin sections were cut using a Leica EM UC7 ultramicrotome equipped with a diamond knife (DiATOME 45°). Sections (80 nm) were collected on uncoated 400-mesh gold grids.

Samples were blocked in a 20μl drop of 2% BSA in TBST (20mM Tris, 500mM NaCl, 0.1% BSA and 0.05% Tween 20) at room temperature for 30 min. Grids were then transferred directly into a 20μl drop of Rabbit anti-NDP52 (1:200, Genetex GTX115378) TBST and incubated for 1 hour. Grids were washed in 6 x TBST. Grids were then moved into a drop of goat anti-rabbit IgG 5nm gold (British Biocell International) diluted 1:50 and then moved to a fresh drop of the same antibody and incubated for 30 min. Excess antibody was removed by washing in 6 x 20μl drops of TBST and 6 x 20μl drops of milliQ water and dried.

Grids were stained for 15 min in 4.5% uranyl acetate in 1% acetic acid solution and then washed in 6 x 20μl drops of milliQ water. Grids were then stained with Reynolds lead citrate for 3 min and washed in 6 x 20μl drops of milliQ water. Electron microscopy was performed using a JEOL-1230 transmission electron microscope operated at an accelerating voltage of 80 kV equipped with a Gatan One View digital camera.

### Graphics

Unless stated, data fitting and plotting was performed using GraphPad Prism 9 and Grafit Version 5 (Erithacus Software Ltd). Cartoons were generated using the BioRender software.

## Data availability

All raw data are available upon request from the corresponding author. The mass spectrometry proteomics data have been deposited to the ProteomeXchange Consortium with the data identifier PXD030238. RNA-Seq data were deposited in the Gene Expression Omnibus (GEO) database under the accession number GSE188567.

## ACKNOWLEDGEMENTS

We thank the UKRI-MRC (MR/M020606/1) and UKRI-STFC (19130001) for funding to C.P.T. and the UKRI-MRC (MR/R024871/1) for a Rutherford Innovation fellowship to A.L.B.P. Aberration-corrected multi-focal microscopy was performed in collaboration with the Advanced Imaging Center at Janelia Research Campus, a facility jointly supported by the Howard Hughes Medical Institute and the Gordon and Betty Moore Foundation. We also thank Darren Griffin (University of Kent) and Alessia Buscaino (University of Kent) for sharing of equipment, and Satya Khuon (Janelia Research Campus) for assisting with cell culture. The JF549 dyes were kindly provided by Luke Lavis (Janelia Research Campus). We thank Anthony Maxwell and Lesley Mitchenhall (John Innes Centre) for assistance preparing ds339 DNA, and Lynn Zechiedrich and Jonathan Fogg (Baylor College of Medicine) for providing plasmids for this work. We also thank Robert Turner (University of Sheffield) for Research Software Engineering support. We wish to acknowledge the Henry Royce Institute for Advanced Materials, funded through EPSRC grants EP/R00661X/1, EP/S019367/1, EP/P02470X/1 and EP/P025285/1 and Robert Moorehead for Dimension XR access and support at Royce@Sheffield

## AUTHOR CONTRIBUTIONS

A.dS and C.P.T. conceived the study. A.dS, E.R, A.L.B.P., Y.H-G. and C.P.T. designed experiments. A.dS and C.P.T. performed single molecule imaging experiments. A.dS, D.E.R and A.L.B.P carried out AFM imaging and A.dS, D.E.R, M.D and A.L.B.P carried out AFM analysis. Imaging was supported by L.W., M.M-F., T-L.C., and J.A. A.dS and Y.H-G. performed and analyzed the genomics experiments. A.dS, H.C.W.R., K. P., S.Y.Z.R and F.L. expressed, purified and performed experiments with recombinant proteins. L. W., J.A., A.dS. and C.P.T. contributed to single molecule data analysis. A.dS, A.S., P.I.J.E., and K.H. performed and analyzed the mass spectrometry experiments. I.B. prepared samples and performed electron microscopy. C.P.T. supervised the study. A.dS and C.P.T. wrote the manuscript with comments from all authors.

