## Supplementary Information and Figures for "Autophagy receptor NDP52 alters DNA conformation to modulate RNA Polymerase II transcription"

### Supplementary Figures

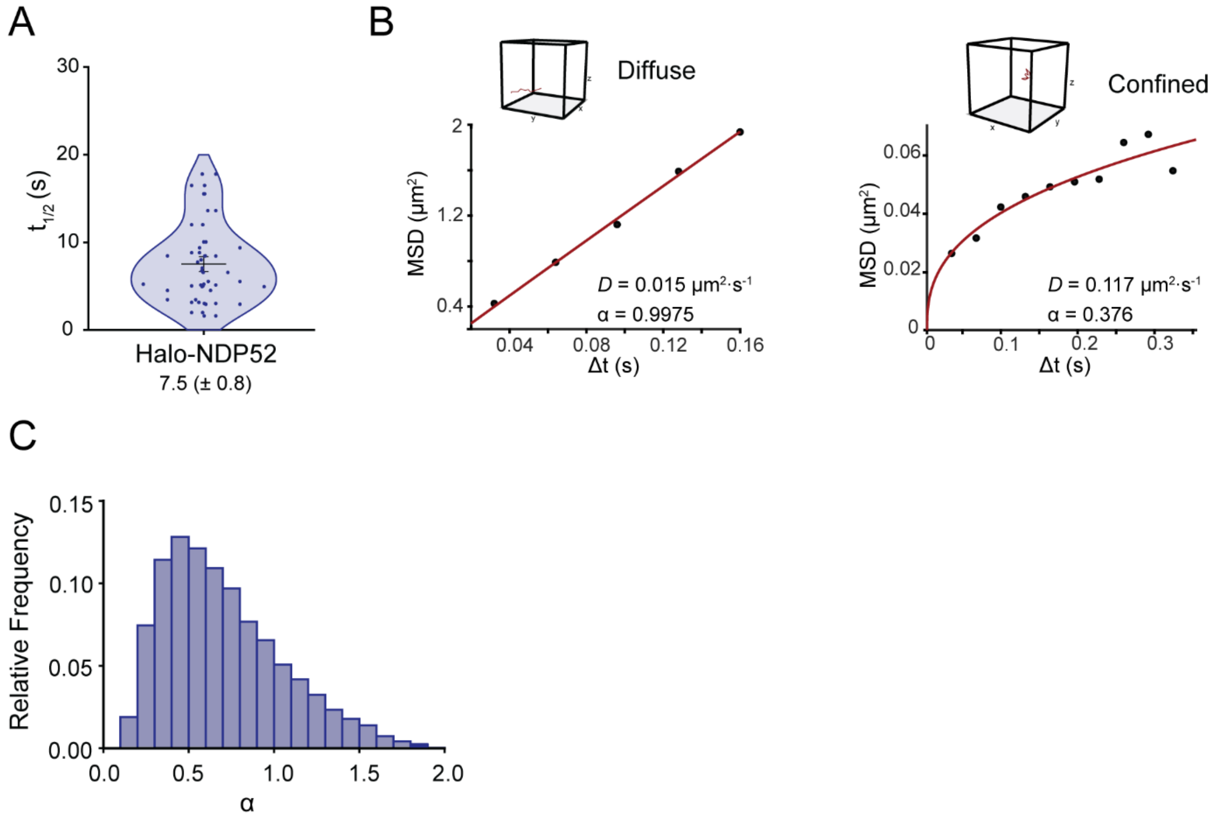


**Supplementary Figure 1: Nuclear dynamics of NDP52. (A)** Violin plot of recovery time ($t_{1/2}$) calculated from FRAP data for Halo NDP52. Each individual point represents one cell. Mean ± SEM value is shown. $n$ = 27 cells. **(B)** Example MSD curves extracted and fitted from acMFM single-molecule tracks. 3D trajectory of each example is shown above the graph and calculated values for diffusion coefficient ($D$) and anomalous diffusion (α) are shown. **(C)** Histogram of calculated α values for all molecules using acMFM. Graph represents values for 14 322 molecules from 51 cells.

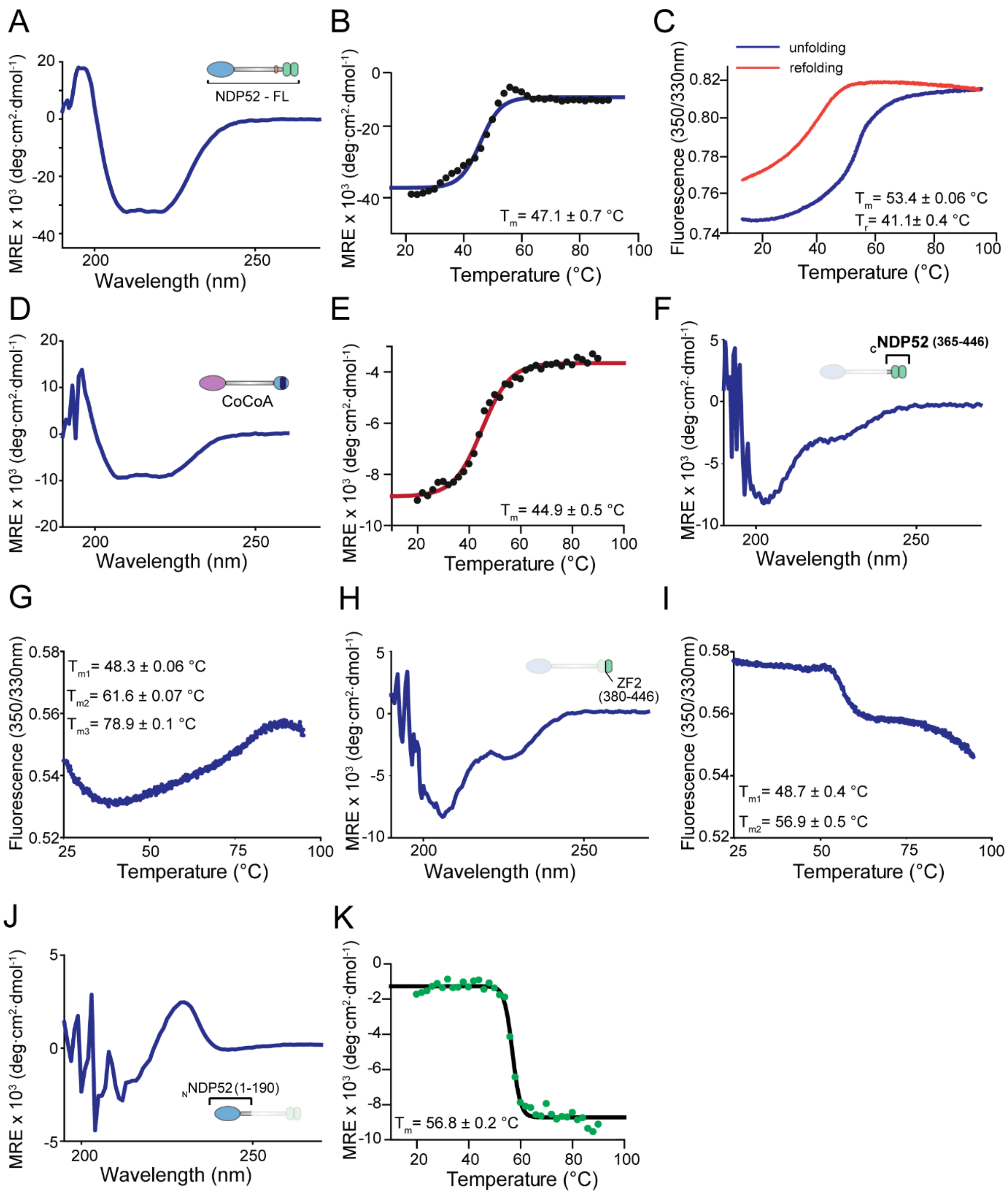


**Supplementary Figure 2: Folding and stability of NDP52 and CoCoA recombinant constructs. (A)** CD spectra of NDP52-FL. **(B)** Thermal denaturation of NDP52-FL following MRE values at 222 nm wavelength. **(C)** Nano-DSF curves for NDP52 thermal denaturation and refolding. **(D)** CoCoA CD spectra. **(E)** Thermal denaturation of CoCoA, following MRE values at 222nm wavelength. **(F)** $_{C}$NDP52 CD spectra. **(G)** Nano-Differential Scanning Fluorimetry (nano-DSF) for $_{C}$NDP52 during thermal denaturation. **(H)** ZF2 CD spectra. **(I)** Nano-DSF for ZF2 during thermal denaturation. **(J)** $_{N}$NDP52 CD spectra. **(K)** Thermal denaturation of $_{N}$NDP52, following MRE values at 215nm wavelength.

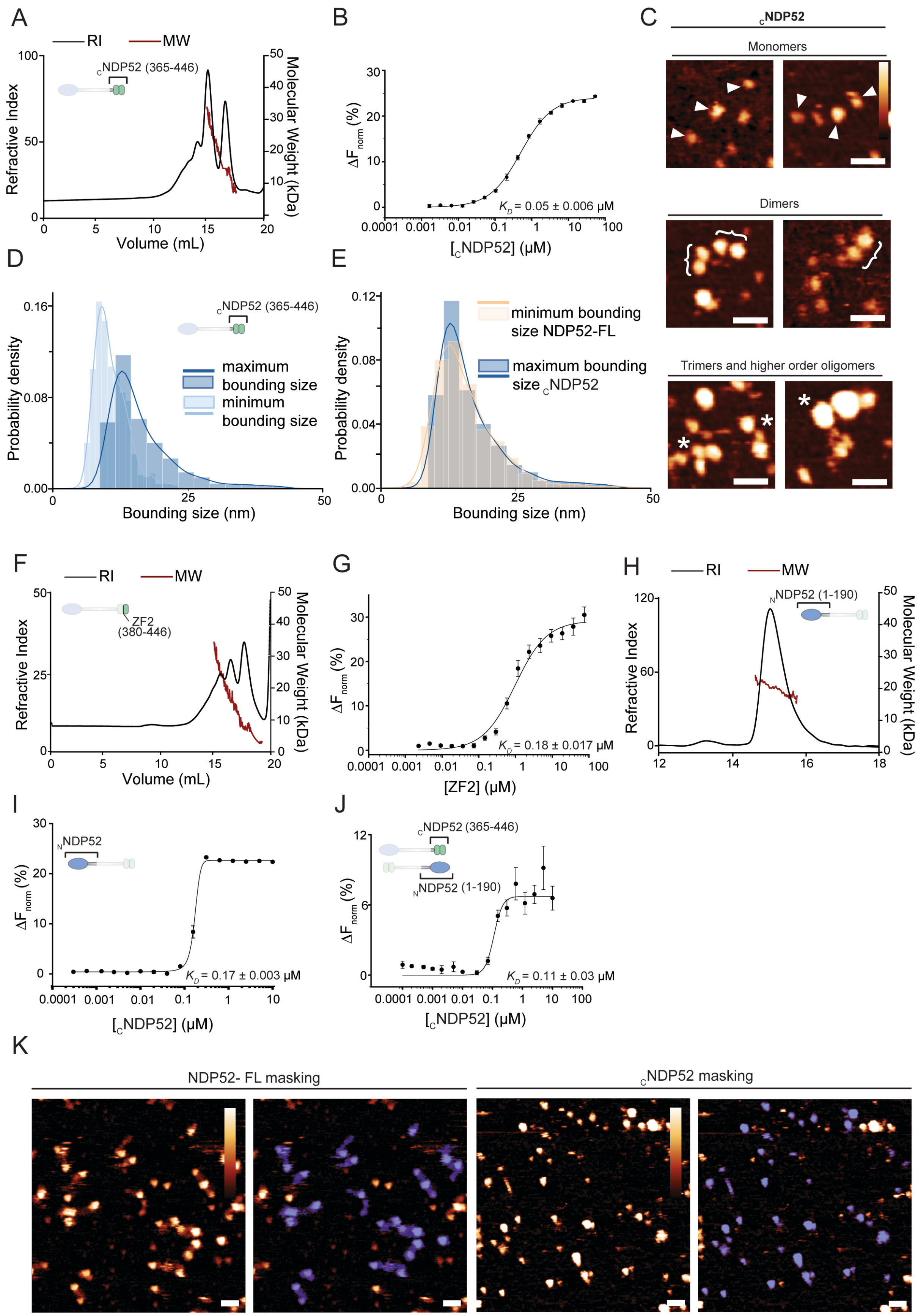


**Supplementary Figure 3: Oligomerisation of individual NDP52 domains. (A)** SEC-MALS profile for the C-terminal of NDP52 ($_C$NDP52), which encompasses both zinc finger domains. Refractive index (RI) trace is shown in black, as well as the calculated molecular weight values, across the peaks (in red). **(B)** Microscale thermophoresis, showing oligomerisation of $_{C}$NDP52, with the calculated $K_D$ displayed in the graph. Values plotted represent average ± SEM of three individual experiments. **(C)** Atomic force microscopy imaging (AFM) of $_{C}$NDP52 showing monomeric, dimeric and higher oligomeric forms of the protein. Scale bar = 25 nm. Height scale = 4.5 nm (scale bar inset in C). **(D)** Histogram and kernel density estimate plots of minimum and maximum bounding sizes, i.e. length and width for $_{C}$NDP52 particles, calculated from masked molecules in AFM images (Supplementary Fig.3K). Peaks in KDE plots were used to determine particle size (KDE max ± SD); minimum = 9 ± 3 nm, maximum = 13 ± 6. N = 1 909 molecules. **(E)** Kernel density estimate distribution showing maximum bounding sizes of $_{C}$NDP52 (data shown in Fig.S3D) compared to minimum bounding size for NDP52-FL (data shown in Fig.4L). **(F)** SEC-MALS trace for ZF2. RI trace is shown in black and calculated molecular weight values are shown in red. **(G)** Microscale thermophoresis, showing oligomerisation of ZF2, with the calculated $K_D$ displayed in the graph. Values represent average ± SEM of three individual experiments. **(H)** SEC-MALS trace for the N-terminal of NDP52 ($_{N}$NDP52), comprising the SKICH domain and a small portion of the coiled-coil region. RI trace is shown in black and calculated molecular weight values are shown in red. **(I)** Microscale thermophoresis, showing oligomerisation of $_{N}$NDP52, with the calculated $K_D$ displayed in the graph. **(J)** MST curve showing interaction between labelled $_{N}$NDP52 titrated with $_{C}$NDP52, with the calculated $K_D$ displayed in the graph. Values represent average ± SEM of three individual experiments. **(K)** AFM images of NDP52-FL and $_{C}$NDP52 with corresponding images showing molecules masked (in blue) and measured by Topostats. Scale bar = 25 nm. Height scale = 4.5 nm (scale bar inset).

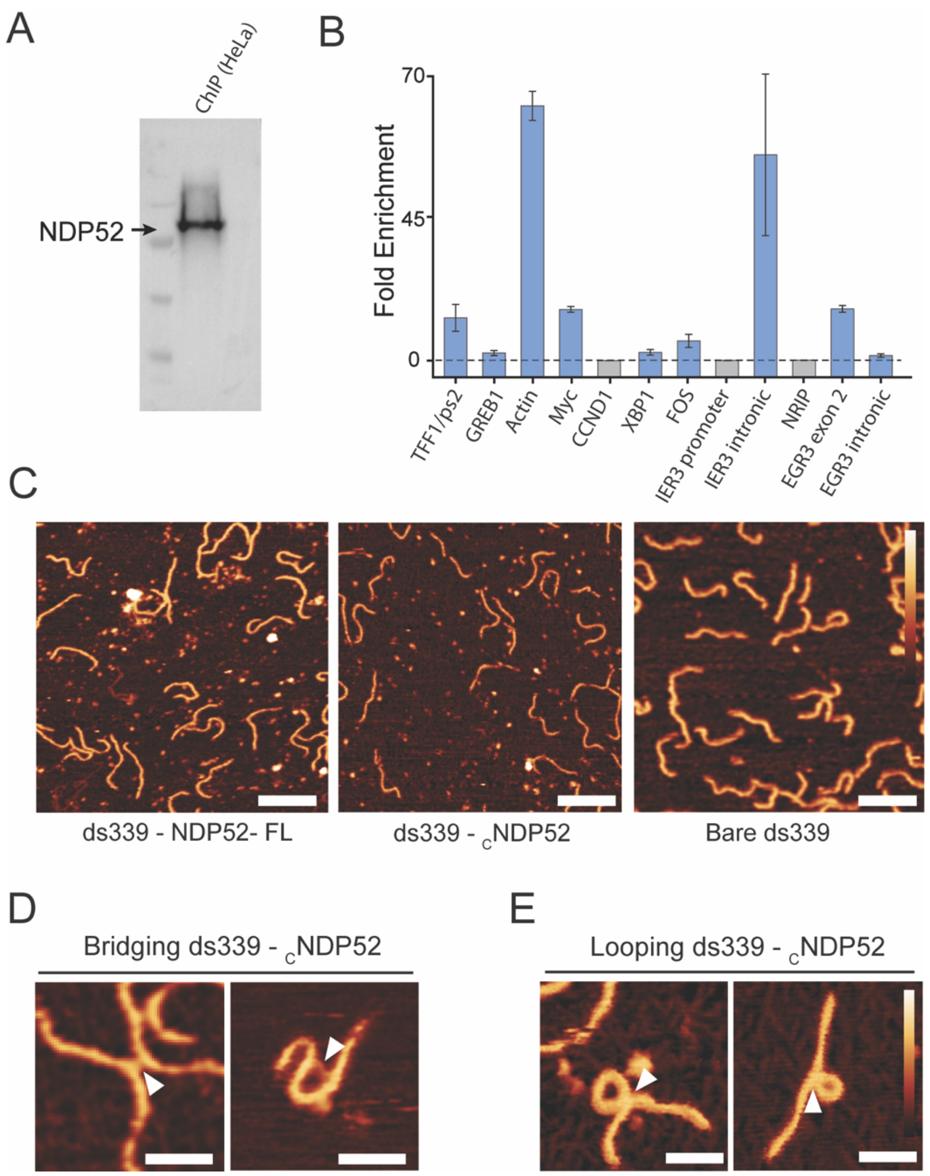


**Supplementary Figure 4: NDP52 binding to DNA in cells and *in vitro*. (A)** Western blot showing presence of NDP52 in chromatin-enriched ChIP sample. **(B)** NDP52 ChIP-qPCR against *loci* specified in the label. Enriched *loci* are shown in blue. Values represent mean ± SEM of three individual experiments. **(C)** AFM images of DNA co-adsorbed with NDP52-FL, $_C$NDP52 and without protein. Scale bars = 100 nm**.** Height scale = 4.5 nm. **(D)** AFM images of ds339 co-adsorbed with $_C$NDP52. Arrowheads show $_C$NDP52 bridging DNA strand(s). Scale bars = 25 nm and height scale inset in E. **(E)** AFM images of ds339 co-adsorbed with $_C$NDP52. Arrowheads show DNA looping induced by $_C$NDP52. Scale bars = 25 nm**.** Height scale = 4.5 nm.

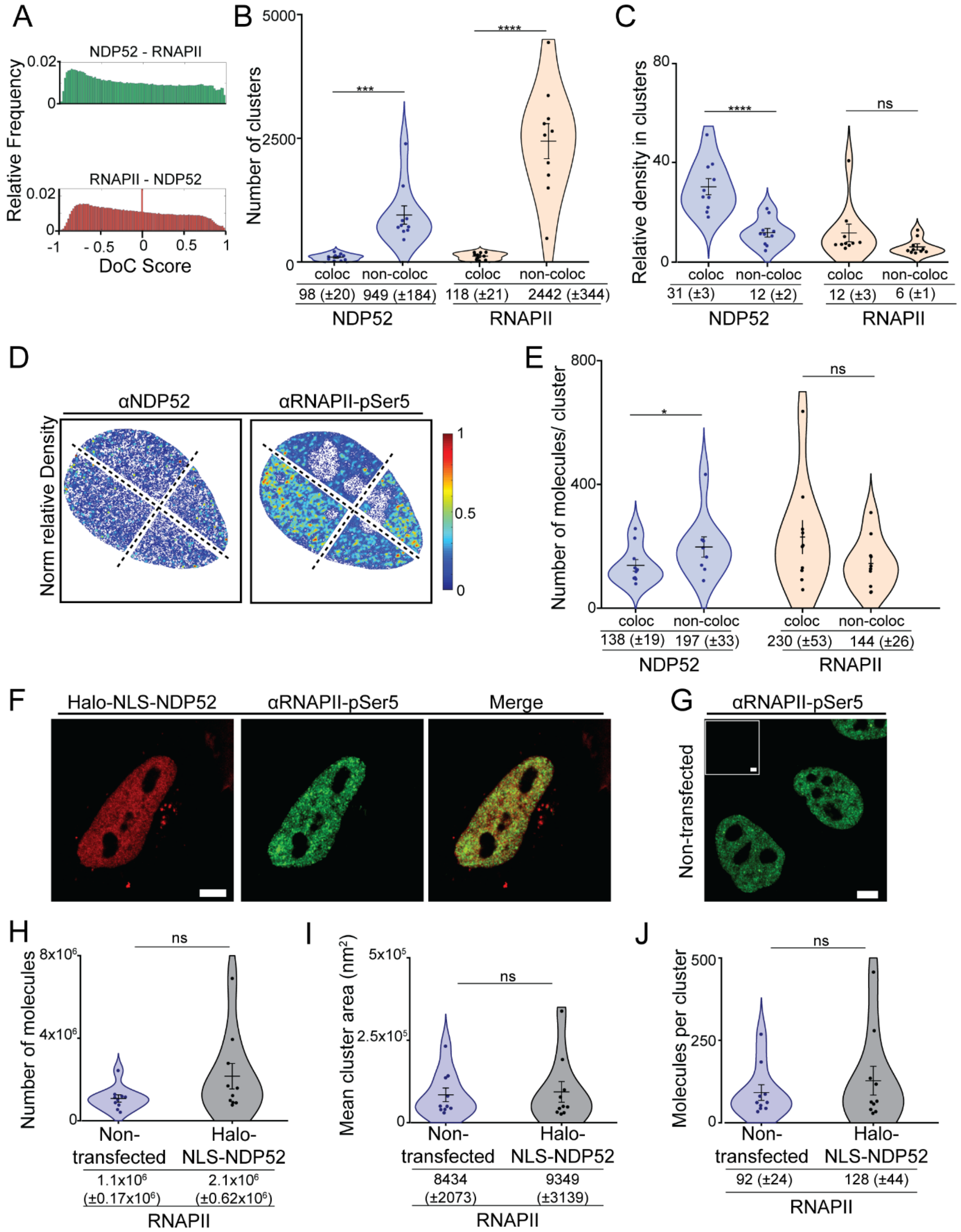


**Supplementary Figure 5: Colocalisation of NDP52 with RNAPII-pSer5. (A)** DoC score histogram for example STORM rendering in Figure 7C and colocalisation map shown in Figure 8D. DoC score of 1 represents perfect colocalisation between molecules, and DoC score -1 represents segregation. DoC score of 0.4 was used as colocalisation threshold. **(B)** Number of clusters considered colocalised or non-colocalised for both NDP52 and RNAPII-pSer5. **(C)** Relative density of molecules in colocalised and non-colocalised NDP52 or RNAPII-pSer5 clusters. **(D)** Example of normalised relative density heat maps for NDP52 and RNAPII-pSer5. Heat maps correspond to the same cell shown in Figure 8B and D. Red represents regions of high molecular density and blue of low molecular density. **(E)** Mean number of molecules per colocalised or non-colocalised clusters for NDP52 or RNAPII-pSer5. n = 10 cells **(F)** Confocal images of HeLa cell transiently expressing Halo-NLS-NDP52 and immunofluorescently stained with RNAPII-pSer5. Scale bar = 5µm. **(G)** Confocal image of non-transfected HeLa cell with immunofluorescent staining of RNAPII-pSer5. Scale bar = 5µm. **(H)** Number of RNAPII-pSer5 molecules in non-transfected or transiently expressing Halo-NLS-NDP52 HeLa cells. **(I)** Mean cluster area in $nm^2$ for RNAPII-pSer5 molecules in non-transfected or transiently expressing Halo-NLS-NDP52 HeLa cells. **(J)** Number of molecules per RNAPII-pSer5 cluster in in non-transfected or transiently expressing Halo-NLS-NDP52 HeLa cells. Mean ± SEM values are shown. Each point represents the average value per cell. n = 10 cells (Non-transfected) n = 10 cells (Halo-NLS-NDP52). *p<0.05; ***p<0.001; ****p<0.0001 by two-tailed t-test.

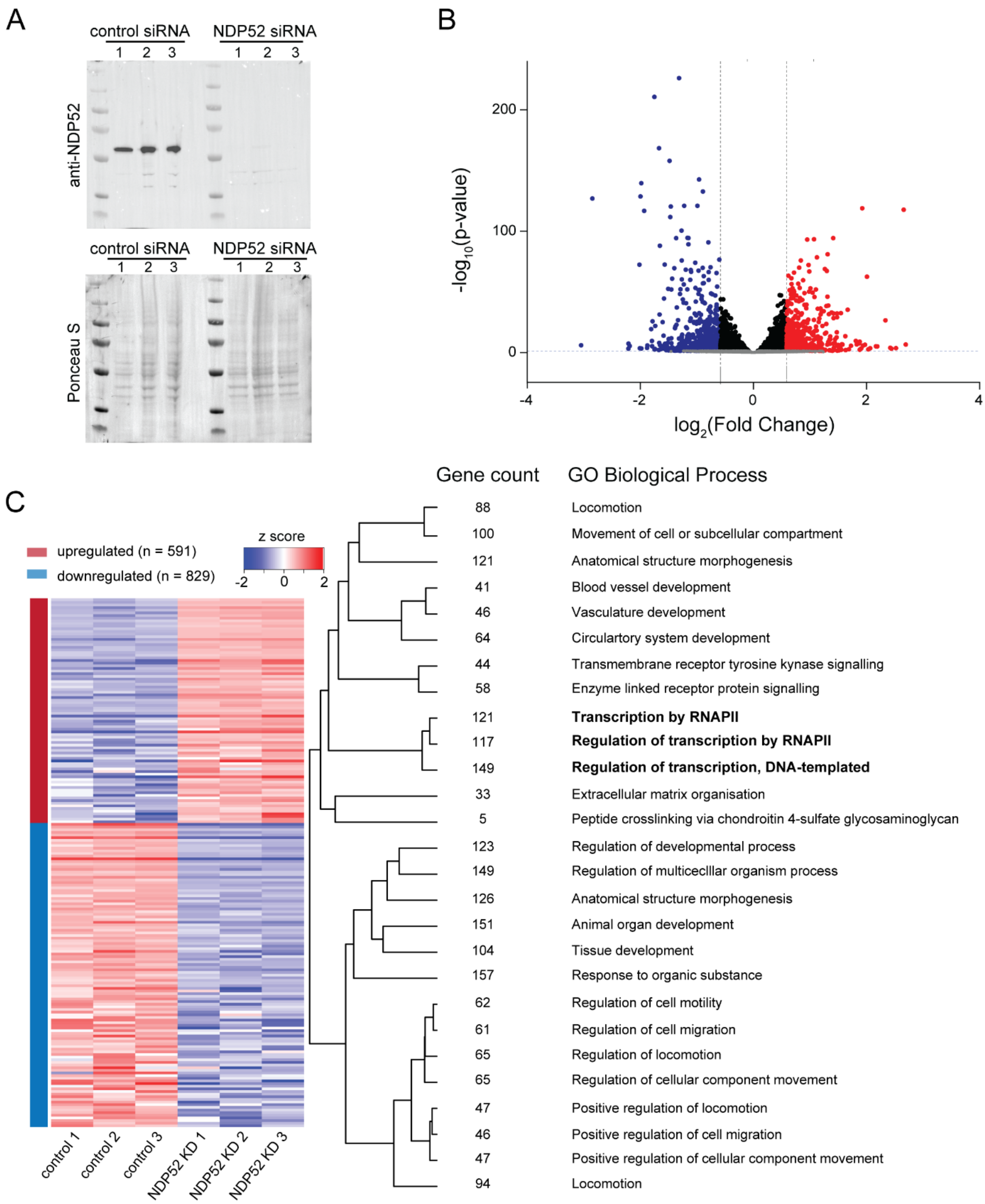


**Supplementary Figure 6: Impact of NDP52 knockdown in gene expression in HeLa cells.** **(A)** Western blot showing NDP52 KD replicates for RNA-Seq. On top, immunoblot against NDP52 is shown and, on the bottom panel, the same blot stained with Ponceau S is shown as loading control. **(B)** Volcano plot of differentially expressed genes following NDP52 KD. **(C)** Heat map showing top down and upregulated genes in HeLa cells showing z-score values for control siRNA and NDP52 KD, obtained from three independent experiments. Dendrogram and enriched GO terms are shown. Number of genes differentially expressed for each GO term is also displayed. All GO terms shown have FDR<0.01. For details see Supplementary Table 4.

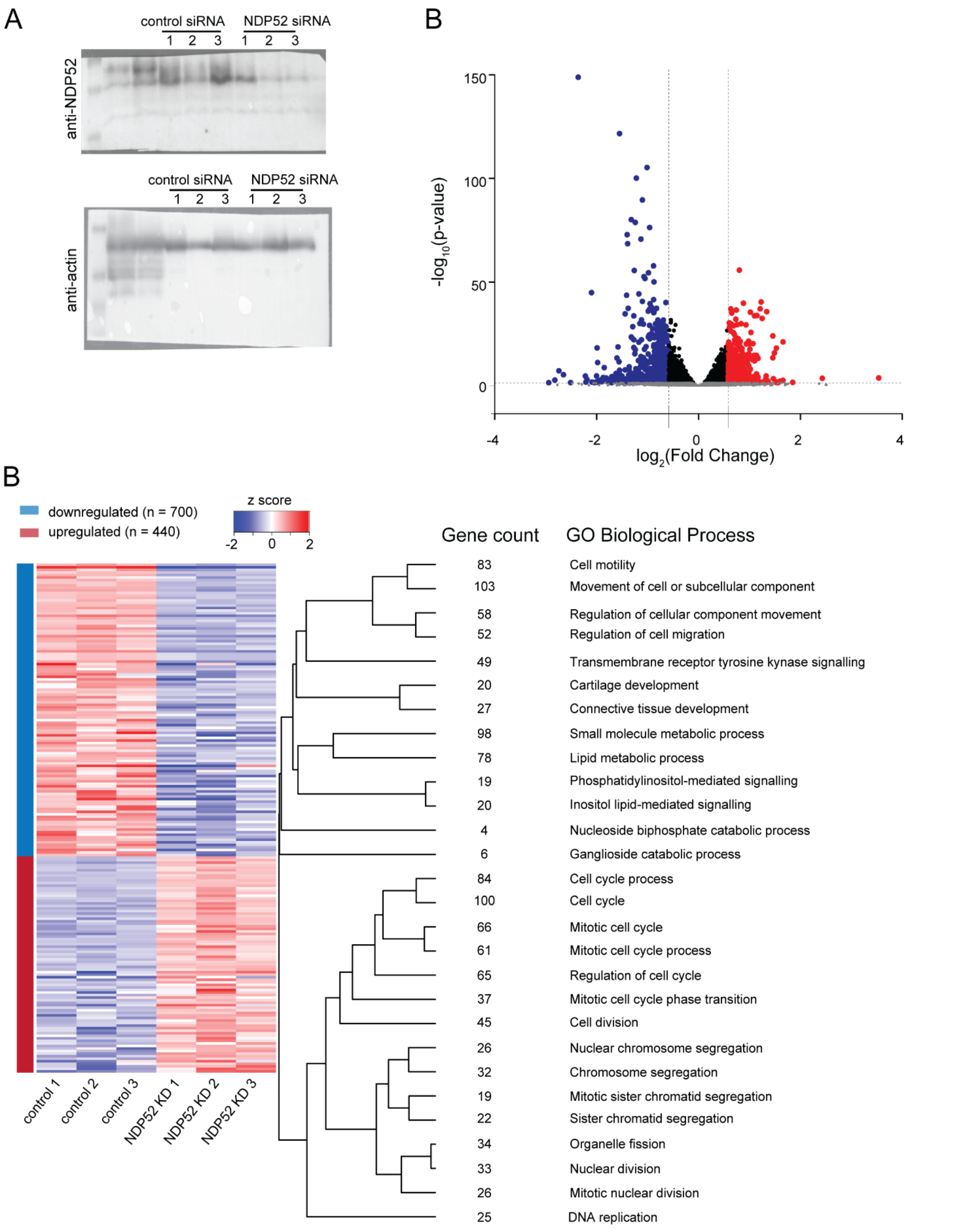


**Supplementary Figure 7: Impact of NDP52 knockdown in gene expression in MCF-7 cells. (A)** Western blot showing NDP52 KD replicates for RNA-Seq. On top, immunoblot against NDP52 is shown and, on the bottom panel, the anti-actin immunoblot is shown. **(B)** Volcano plot of differentially expressed genes following NDP52 KD in MCF-7 cells. **(C)** Heat map showing top down and upregulated genes and z-score values for control siRNA and NDP52 KD from three independent experiments. Dendrogram and enriched GO terms are shown. Number of genes differentially expressed for each GO term is also displayed. All GO terms shown have FDR<0.01. For details see Supplementary Table 5.

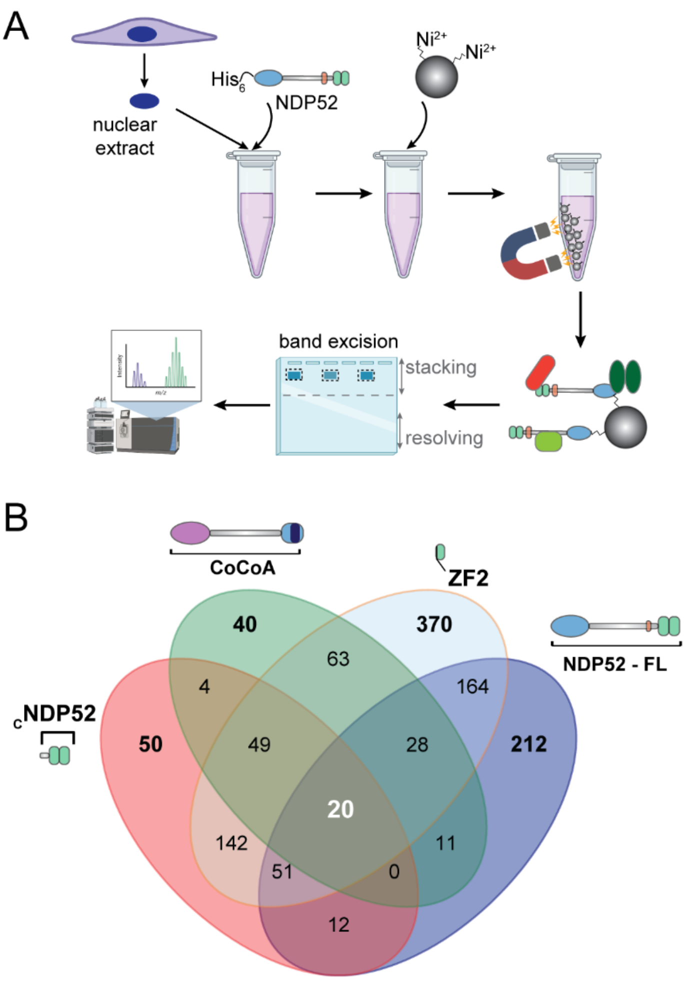


C

| Hit | ZF2 | | known function |
|---|---|---|---|
| | log$_2$(FC) | p | |
| GATAD2B | ∞ | ** | transcription |
| SWT1 | ∞ | *** | transcription |
| PARG | ∞ | ** | DNA repair |
| WIZ | ∞ | ** | transcription |
| SCMH1 | ∞ | *** | transcription |
| FANCA | ∞ | ** | DNA repair |
| ARNTL | ∞ | *** | transcription |
| RECQL5 | ∞ | * | helicase |
| TTF2 | ∞ | **** | transcription |
| RERE | 7.1 | **** | transcription |
| ARID1B | ∞ | *** | chromatin regulation |
| KDM5C | ∞ | ** | chromatin regulation |
| FUBP1 | ∞ | * | transcription |
| SETDB1 | ∞ | ** | chromatin regulation |
| NELFE | ∞ | * | transcription |
| MED12 | ∞ | *** | transcription |
| TCF4 | -- | -- | transcription |
| SKIV2L | ∞ | * | helicase |

**Supplementary Figure 8: Recombinant and endogenous NDP52 interactomes in non-treated and α-amanitin treated HeLa (A)** Diagram depicting recombinant protein proteomics protocol. Briefly, following nuclear isolation from HeLa cells, recombinant constructs were incubated with nuclear extract. His-tagged constructs bound to interacting partners were pulled-down using magnetic Ni-NTA beads. Samples were run in SDS-PAGE gels and all proteins from each replicate were excised as a single band from the stacking portion of the polyacrylamide gel, for LC-MS/MS experiments. **(B)** Venn diagram of hits found in label-free quantitative LC-MS/MS data for pull-downs of recombinant protein from HeLa nuclear extract. **(C)** Examples of top hits for NDP52 and CoCoA shown in Figures 8B and 8D, and their identification in ZF2 proteomics data. log$_2$FC is relative to beads control.

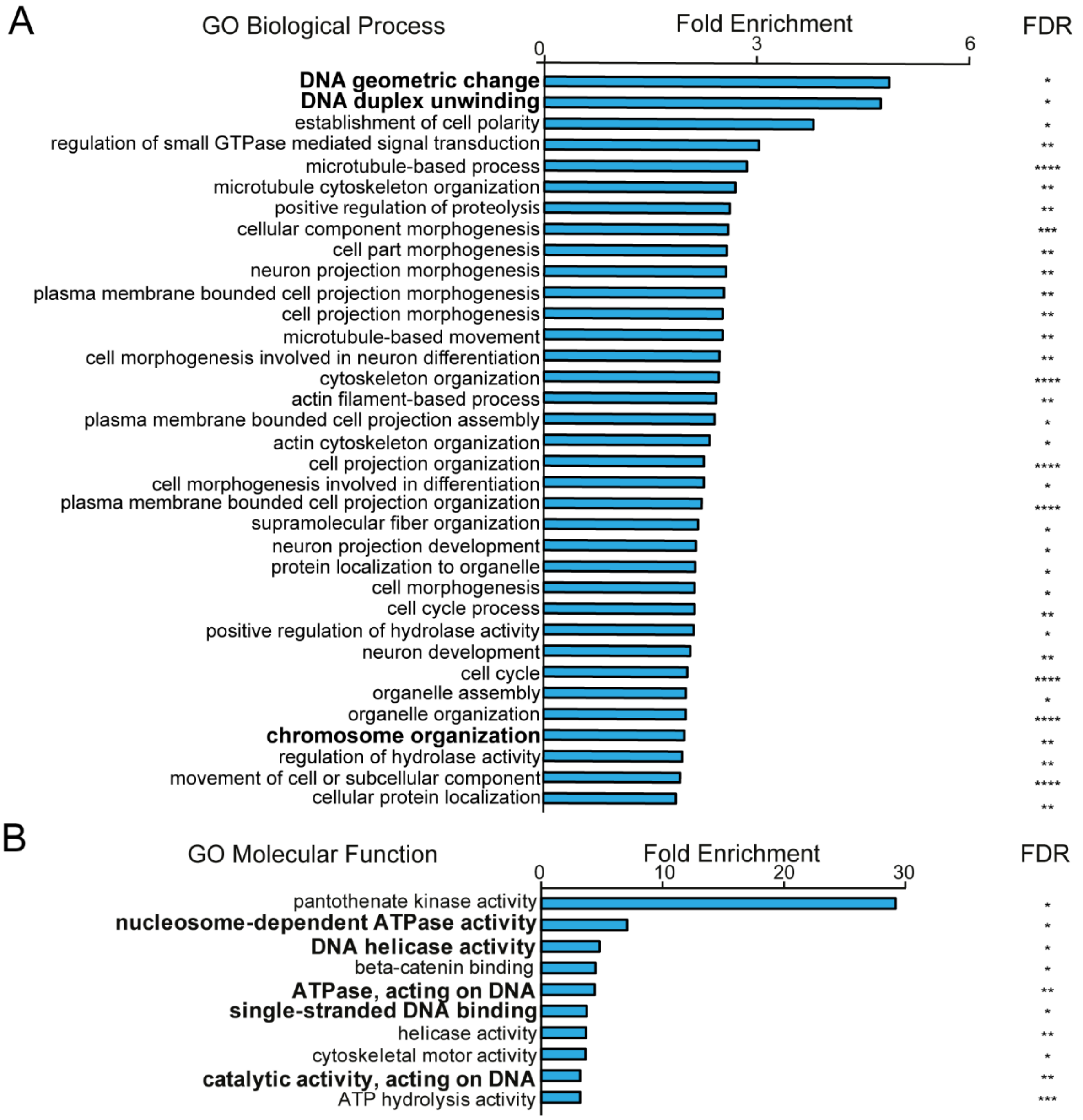


**Supplementary Figure 9: GO Biological Process and Molecular Function analysis for NDP52-FL-interacting proteins in HeLa nuclear extract. (A)** Top 35 Gene Ontology Biological Process annotation terms for NDP52-FL interactions. **(B)** Top 10 Gene Ontology molecular function terms for identified interactors of NDP52-FL. Fold enrichment and FDR values are shown. * FDR<0.05, ** FDR<0.01, *** FDR<0.001, **** FDR<0.0001. For full lists refer to Supplementary tables 6 and 7.

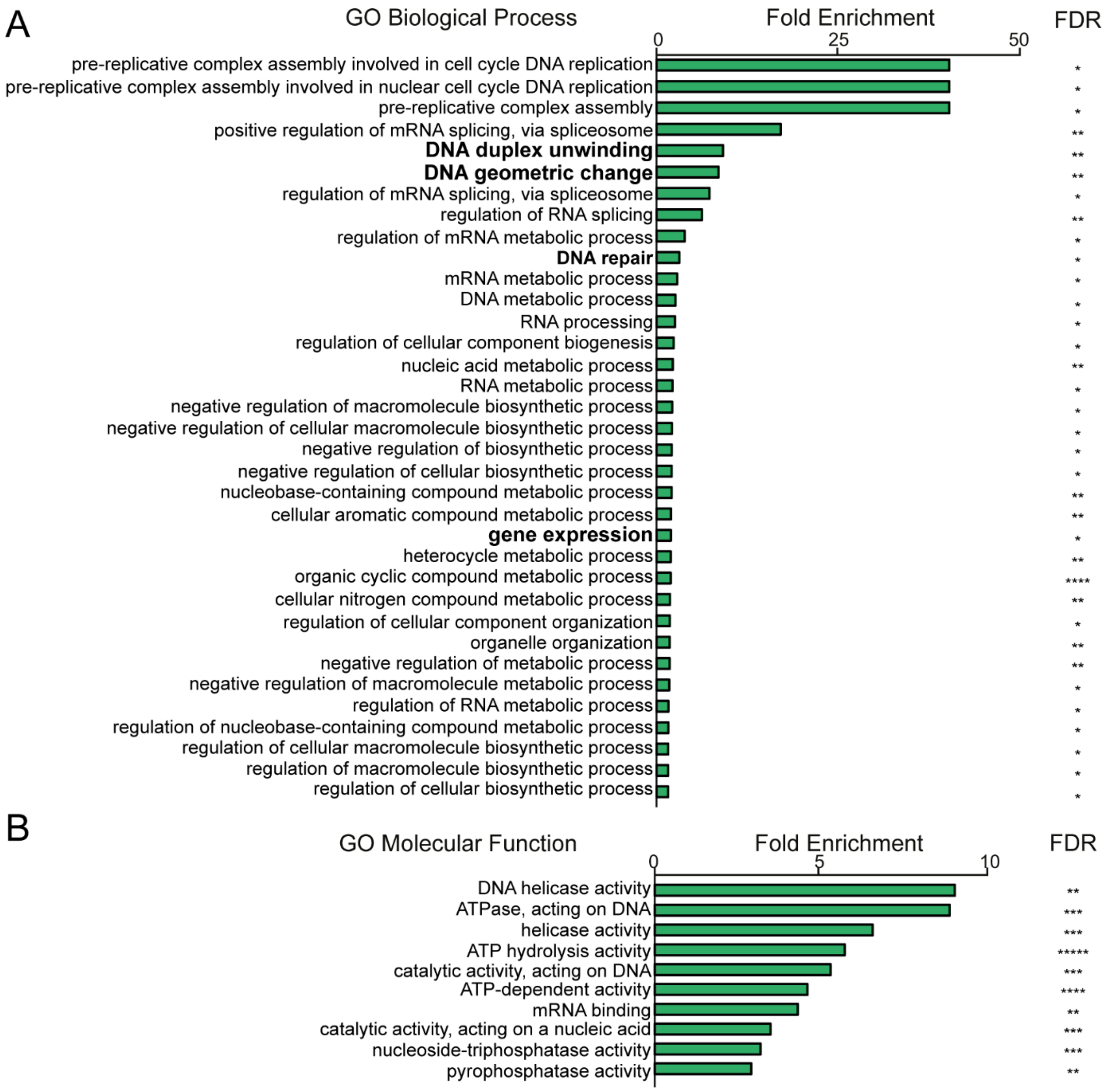


**Supplementary Figure 10: GO Biological Process and Molecular Function analysis for CoCoA-interacting proteins in HeLa nuclear extract. (A)** Top 35 Gene Ontology Biological Process annotation terms for CoCoA interactions. **(B)** Top 10 Gene Ontology molecular function terms for identified interactors of CoCoA. Fold enrichment and FDR values are shown. * FDR<0.05, ** FDR<0.01, *** FDR<0.001, **** FDR<0.0001. For full lists refer to Supplementary tables 8 and 9.

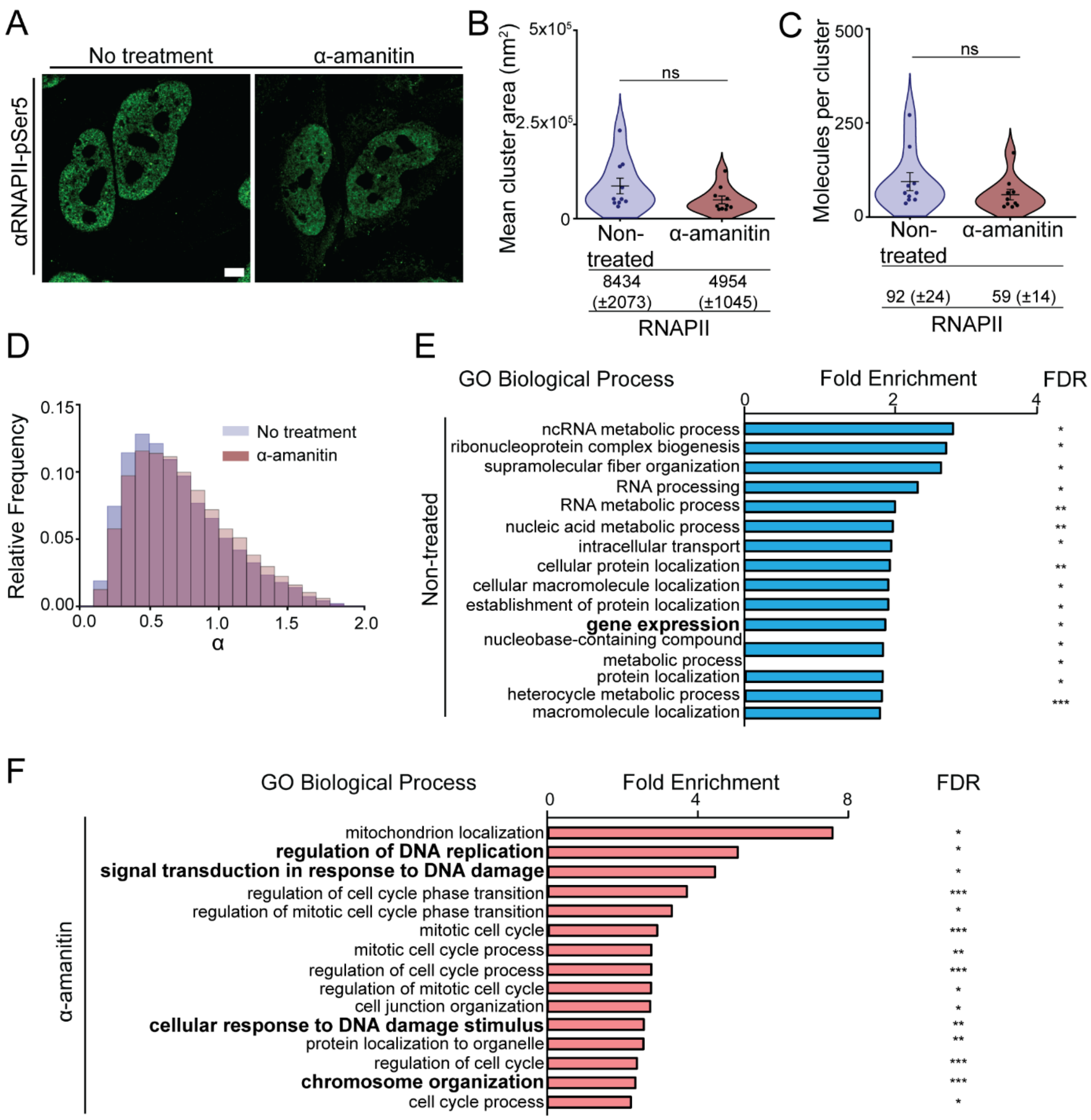


**Supplementary Figure 11: Effect on transcription inhibition on the nuclear spatial organisation and dynamics of NDP52. (A)** Confocal images of RNAPII-pSer5 in non-treated and α-amanitin treated HeLa cells. **(B)** Mean cluster area in nm$^2$ for RNAPII-pSer5 in non-treated and α-amanitin treated cells. **(C)** Number of RNAPII-pSer5 per cluster in non-treated and α-amanitin treated cells. Mean ± SEM values are shown. Each point represents the average value per cell. *n* = 10 cells (non-treated) *n* = 10 cells (α-amanitin). **(D)** Anomalous diffusion (α) histograms for cells transiently expressing Halo-NLS-NDP52, non-treated or following α-amanitin treatment. Histogram represents values for 14 322 molecules from 51 cells non-treated condition (in blue – same as shown in Supplementary Figure 2C) and 14 492 molecules from 50 cells for α-amanitin treatment (in red). **(E-F)** Changes in the interactome of endogenous NDP52 in non-treated and α-amanitin treated HeLa cells. **(E)** Top 15 Gene Ontology Biological Process annotation terms for NDP52 interactions in non-treated HeLa cells. **(F)** Top 15 Gene Ontology molecular function terms for identified for interactions of NDP52 following α-amanitin treatment. Fold enrichment and FDR values are shown. * FDR<0.05, ** FDR<0.01, *** FDR<0.001, **** FDR<0.0001. For full lists refer to Supplementary tables 10 and 11.

### Supplementary Tables

**Supplementary Table 1 – List of constructs.**

| **Construct** | **Source** |
|---|---|
| Human pET151 Halo-NDP52 | Synthetic Gene |
| Human pET151 Halo-NLS-NDP52 | Synthetic Gene |
| Human pET151 NDP52-FL (1-end) | Synthetic Gene |
| Human pET151 CoCoA (1-end) | Synthetic Gene |
| Human pET151 $_{C}$NDP52 (365-end) | Synthetic Gene |
| Human pET151 ZF2 (380-end) | Synthetic Gene |
| Human pET151 $_{N}$NDP52 (1-190) | Synthetic Gene |

**Supplementary Table 2 – ChIP-qPCR primers**

| Description | Oligonucleotide sequence |
|---|---|
| TFF1/PS2 Forward | CCATGTTGGCCAGGCTAGTC |
| TFF1/PS2 Reverse | ACAACAGTGGCTCACGGGGT |
| GREB1 Forward | GCAGTGAAAAAAAGTGTGGCAACTGGG |
| GREB1 Reverse | GACCCACAGAAATGAAAAGGCAGCAAACT |
| Actin Forward | GACCTTCAACACCCCAGCC |
| Actin Reverse | GTCACGCACGATTTCCCGCT |
| Myc Forward | GGCTCACCCTTGCTGATGCT |
| Myc Reverse | GCTCTGGGCACACACATTGG |
| CCND1 Forward | GCCAGACAACATTAGCCACA |
| CCND1 Reverse | CCCATTTCTTAGCTGGGTCA |
| XBP1 Forward | ATACTTGGCAGCCTGTGACC |
| XBP1 Reverse | GGTCCACAAAGCAGGAAAAA |
| FOS Forward | GGGCACTTACACACACATGC |
| FOS Reverse | ATGATACTCGCGAGCAGAGG |
| IER3 Promoter Forward | CCCCTGCTTCTTCTCAGTTG |
| IER3 Promoter Reverse | AAAAGATGCACGGATTGGAG |
| IER3 Intronic Forward | CGCCGAAGTCTCACACAGTA |
| IER3 Intronic Reverse | AGGAGAAGAAATGGGGAGGA |
| NRIP Forward | GACCTGGCCCAGATAGATCA |
| NRIP Reverse | TATAATGTAGGGGCGCAACC |
| EGR3 exon 2 Forward | GGTGCCTGAGAAGAGGTGAG |
| EGR3 exon 2 Reverse | CCATGTGGATGAATGAGGTG |
| EGR3 Intronic Forward | TCACGTACCACACACACACG |
| EGR3 Intronic Reverse | TCCGGGTCTGAAACTACCTG |

**Supplementary Table 3 – List of DNA constructs.**

| Description | Oligonucleotide sequence |
|---|---|
| ds15A 3'FAM | TTAGTTGTTCTCTGG* |
| ds15B | CCAGAGAACAACTAA |
| ds40A 3'FAM | TTAGTTGTTCGTAGTGCTCGTCTGGCTCTGGATTACCCGC* |
| ds40B | GCGGGTAATCCAGAGCCAGACGAGCACTACGAACAACTAA |
| ss40 3'FAM | TTAGTTGTTCGTAGTGCTCGTCTGGCTCTGGATTACCCGC* |
| ds339 | TATGTTGTGTTTTACAGTATTATGTAGTCTGTTTTTTATGCA AAATCTAATTTAATATATTGATATTTATATCATTTTACGTTTC TCGTTCAGCTTTTTTATACTAACTTGAGCGAAACGGGAAGG GTTTTCACCGATATCACCGAAACGCGCGAGGCAGCTGTAT GGCGAAATGAAAGAACAAACTTTCTTGTACGCGGTGGTGA GAGAGAGAGAGAGATACGACTACTATCAGCCGGAAGCCTA TGTACCGAGTTCCGACACTTTCATTGAGAAAGATGCCTCA GCTCTGTTACAGGTCACTAATACCATCTAAGTAGTTGATTC ATAGTGACTGCA |

**Supplementary Table 4 – HeLa RNA-Seq enriched GO Biological processes**

| $p_{adj}$ | Gene count | Pathways |
|---|---|---|
| 6.12E-13 | 157 | Response to organic substance |
| 1.23E-11 | 149 | Regulation of multicellular organismal process |
| 6.06E-11 | 126 | Anatomical structure morphogenesis |
| 6.06E-11 | 46 | Positive regulation of cell migration |
| 9.75E-11 | 61 | Regulation of cell migration |
| 9.75E-11 | 65 | Regulation of locomotion |
| 9.75E-11 | 47 | Positive regulation of locomotion |
| 9.75E-11 | 47 | Positive regulation of cellular component movement |
| 1.34E-10 | 104 | Tissue development |
| 2.83E-10 | 62 | Regulation of cell motility |
| 3.50E-10 | 123 | Regulation of developmental process |
| 3.74E-10 | 65 | Regulation of cellular component movement |
| 6.00E-10 | 151 | Animal organ development |
| 8.97E-10 | 94 | Locomotion |
| 0.000123 | 64 | Circulatory system development |
| 0.000173 | 5 | Peptide cross-linking via chondroitin 4-sulfate glycosaminoglycan |
| 0.000219 | 33 | Extracellular matrix organization |
| 0.000362 | 46 | Vasculature development |
| 0.000963 | 121 | Anatomical structure morphogenesis |
| 0.001308 | 117 | Regulation of transcription by RNA polymerase II |
| 0.00131 | 100 | Movement of cell or subcellular component |
| 0.001842 | 121 | Transcription by RNA polymerase II |
| 0.001842 | 58 | Enzyme linked receptor protein signaling pathway |
| 0.001842 | 44 | Transmembrane receptor protein tyrosine kinase signaling pathway |
| 0.002537 | 88 | Locomotion |
| 0.003258 | 41 | Blood vessel development |
| 0.003744 | 149 | Regulation of transcription, DNA-templated |

**Supplementary Table 5 – MCF-7 RNA-Seq enriched GO Biological processes**

| $p_{adj}$ | Gene count | Pathways |
|---|---|---|
| 7.33E-05 | 27 | Connective tissue development |
| 0.000519 | 49 | Transmembrane receptor protein tyrosine kinase signaling pathway |
| 0.00239 | 6 | Ganglioside catabolic process |
| 0.00239 | 20 | Cartilage development |
| 0.002642 | 78 | Lipid metabolic process |
| 0.002642 | 103 | Movement of cell or subcellular component |
| 0.002642 | 20 | Inositol lipid-mediated signaling |
| 0.002904 | 52 | Regulation of cell migration |
| 0.002904 | 4 | Nucleoside bisphosphate catabolic process |
| 0.002904 | 98 | Small molecule metabolic process |
| 0.002904 | 58 | Regulation of cellular component movement |
| 0.002955 | 19 | Phosphatidylinositol-mediated signaling |
| 0.002955 | 83 | Cell motility |
| 2.69E-16 | 100 | Cell cycle |
| 4.74E-16 | 84 | Cell cycle process |
| 7.62E-14 | 61 | Mitotic cell cycle process |
| 7.97E-14 | 66 | Mitotic cell cycle |
| 2.62E-11 | 45 | Cell division |
| 7.52E-11 | 32 | Chromosome segregation |
| 4.30E-10 | 65 | Regulation of cell cycle |
| 7.62E-09 | 22 | Sister chromatid segregation |
| 9.57E-09 | 26 | Nuclear chromosome segregation |
| 2.36E-08 | 33 | Nuclear division |
| 5.40E-08 | 26 | Mitotic nuclear division |
| 7.25E-08 | 25 | DNA replication |
| 7.25E-08 | 34 | Organelle fission |
| 8.16E-08 | 19 | Mitotic sister chromatid segregation |
| 2.36E-07 | 37 | Mitotic cell cycle phase transition |

**Supplementary Table 6 – Enriched GO Biological processes from recombinant NDP52-FL nuclear interactome**

| Fold Enrichment | Genes in list | FDR | Functional Category – Biological Process |
|---|---|---|---|
| 4.75 | 11 | 0.0142 | DNA geometric change (GO:0032392) |
| 3.8 | 10 | 0.0315 | DNA duplex unwinding (GO:0032508) |
| 3.03 | 12 | 0.0407 | establishment of cell polarity (GO:0030010) |
| 2.86 | 24 | 0.00249 | regulation of small GTPase mediated signal transduction (GO:0051056) |
| 2.7 | 56 | 3.17E-08 | microtubule-based process (GO:0007017) |
| 2.62 | 36 | 2.63E-04 | microtubule cytoskeleton organization (GO:0000226) |
| 2.6 | 25 | 0.01 | positive regulation of proteolysis (GO:0045862) |
| 2.58 | 38 | 2.40E-04 | cellular component morphogenesis (GO:0032989) |
| 2.57 | 32 | 0.00194 | cell part morphogenesis (GO:0032990) |
| 2.54 | 30 | 0.00305 | neuron projection morphogenesis (GO:0048812) |
| 2.52 | 30 | 0.00511 | plasma membrane bounded cell projection morphogenesis (GO:0120039) |
| 2.52 | 30 | 0.00535 | cell projection morphogenesis (GO:0048858) |
| 2.48 | 23 | 0.0303 | microtubule-based movement (GO:0007018) |
| 2.47 | 26 | 0.0197 | cell morphogenesis involved in neuron differentiation (GO:0048667) |
| 2.43 | 74 | 1.22E-08 | cytoskeleton organization (GO:0007010) |
| 2.41 | 37 | 0.00142 | actin filament-based process (GO:0030029) |
| 2.34 | 24 | 0.0411 | plasma membrane bounded cell projection assembly (GO:0120031) |
| 2.26 | 32 | 0.0102 | actin cytoskeleton organization (GO:0030036) |
| 2.26 | 65 | 3.72E-06 | cell projection organization (GO:0030030) |
| 2.23 | 30 | 0.023 | cell morphogenesis involved in differentiation (GO:0000904) |
| 2.18 | 61 | 2.57E-05 | plasma membrane bounded cell projection organization (GO:0120036) |
| 2.15 | 29 | 0.0413 | supramolecular fiber organization (GO:0097435) |
| 2.14 | 36 | 0.0119 | neuron projection development (GO:0031175) |
| 2.13 | 35 | 0.0166 | protein localization to organelle (GO:0033365) |
| 2.13 | 37 | 0.0143 | cell morphogenesis (GO:0000902) |
| 2.12 | 44 | 0.00294 | cell cycle process (GO:0022402) |
| 2.07 | 32 | 0.0353 | positive regulation of hydrolase activity (GO:0051345) |
| 2.03 | 43 | 0.0068 | neuron development (GO:0048666) |
| 2.01 | 63 | 2.44E-04 | cell cycle (GO:0007049) |
| 2.01 | 38 | 0.028 | organelle assembly (GO:0070925) |
| 1.99 | 173 | 4.25E-16 | organelle organization (GO:0006996) |
| 1.96 | 52 | 0.00292 | chromosome organization (GO:0051276) |
| 1.93 | 52 | 0.00423 | regulation of hydrolase activity (GO:0051336) |

| | | | |
|---|---|---|---|
| 1.87 | 70 | 2.45E-04 | movement of cell or subcellular component (GO:0006928) |
| 1.87 | 66 | 0.00143 | cellular protein localization (GO:0034613) |
| 1.86 | 46 | 0.0318 | biological adhesion (GO:0022610) |
| 1.84 | 66 | 0.00211 | cellular macromolecule localization (GO:0070727) |
| 1.84 | 45 | 0.0404 | cell adhesion (GO:0007155) |
| 1.83 | 45 | 0.0397 | regulation of cellular component biogenesis (GO:0044087) |
| 1.83 | 56 | 0.00954 | regulation of organelle organization (GO:0033043) |
| 1.81 | 56 | 0.0094 | protein-containing complex assembly (GO:0065003) |
| 1.79 | 105 | 6.16E-06 | cellular component assembly (GO:0022607) |
| 1.79 | 243 | 1.15E-18 | cellular component organization (GO:0016043) |
| 1.77 | 114 | 2.75E-06 | cellular component biogenesis (GO:0044085) |
| 1.75 | 61 | 0.0108 | protein-containing complex subunit organization (GO:0043933) |
| 1.7 | 247 | 2.10E-18 | cellular component organization or biogenesis (GO:0071840) |
| 1.66 | 83 | 0.00228 | protein localization (GO:0008104) |
| 1.65 | 69 | 0.0208 | cell development (GO:0048468) |
| 1.6 | 96 | 0.00143 | cellular localization (GO:0051641) |
| 1.6 | 97 | 0.00299 | regulation of cellular component organization (GO:0051128) |
| 1.59 | 90 | 0.00614 | nervous system development (GO:0007399) |
| 1.51 | 98 | 0.00314 | regulation of catalytic activity (GO:0050790) |
| 1.5 | 119 | 0.00295 | regulation of molecular function (GO:0065009) |

**Supplementary Table 7 – Enriched GO Molecular Functions from recombinant NDP52-FL nuclear interactome**

| Fold Enrichment | Genes in list | FDR | Functional Category – Molecular Function |
|---|---|---|---|
| 29.2 | 3 | 0.0468 | pantothenate kinase activity (GO:0004594) |
| 7.08 | 6 | 0.0372 | nucleosome-dependent ATPase activity (GO:0070615) |
| 4.8 | 9 | 0.0213 | DNA helicase activity (GO:0003678) |
| 4.47 | 10 | 0.0173 | beta-catenin binding (GO:0008013) |
| 4.41 | 12 | 0.00517 | ATPase, acting on DNA (GO:0008094) |
| 3.74 | 12 | 0.0182 | single-stranded DNA binding (GO:0003697) |
| 3.7 | 15 | 0.00408 | helicase activity (GO:0004386) |
| 3.66 | 11 | 0.0373 | cytoskeletal motor activity (GO:0003774) |
| 3.2 | 19 | 2.85E-03 | catalytic activity, acting on DNA (GO:0140097) |
| 3.2 | 23 | 4.85E-04 | ATP hydrolysis activity (GO:0016887) |
| 3.13 | 16 | 0.0125 | phosphatase binding (GO:0019902) |
| 2.8 | 22 | 4.03E-03 | mRNA binding (GO:0003729) |
| 2.76 | 33 | 1.01E-04 | GTPase activator activity (GO:0005096) |
| 2.74 | 36 | 4.11E-05 | ATP-dependent activity (GO:0140657) |
| 2.68 | 34 | 1.14E-04 | GTPase regulator activity (GO:0030695) |
| 2.68 | 34 | 1.09E-04 | nucleoside-triphosphatase regulator activity (GO:0060589) |
| 2.59 | 18 | 0.0366 | microtubule binding (GO:0008017) |
| 2.53 | 21 | 0.0184 | cadherin binding (GO:0045296) |
| 2.39 | 33 | 0.00139 | cell adhesion molecule binding (GO:0050839) |
| 2.18 | 41 | 0.00144 | enzyme activator activity (GO:0008047) |
| 2.16 | 33 | 0.00824 | catalytic activity, acting on a nucleic acid (GO:0140640) |
| 2.15 | 25 | 0.0483 | actin binding (GO:0003779) |
| 2.14 | 39 | 2.85E-03 | structural molecule activity (GO:0005198) |
| 2.14 | 55 | 6.53E-05 | cytoskeletal protein binding (GO:0008092) |
| 2.07 | 83 | 9.42E-07 | adenyl ribonucleotide binding (GO:0032559) |
| 2.06 | 79 | 1.52E-06 | ATP binding (GO:0005524) |
| 2.05 | 83 | 9.16E-07 | adenyl nucleotide binding (GO:0030554) |
| 2 | 32 | 3.13E-02 | nucleoside-triphosphatase activity (GO:0017111) |
| 1.99 | 86 | 1.63E-06 | RNA binding (GO:0003723) |

| | | | |
|---|---|---|---|
| 1.85 | 91 | 9.13E-06 | purine ribonucleotide binding (GO:0032555) |
| 1.84 | 87 | 1.97E-05 | purine ribonucleoside triphosphate binding (GO:0035639) |
| 1.84 | 91 | 1.37E-05 | purine nucleotide binding (GO:0017076) |
| 1.84 | 91 | 1.30E-05 | ribonucleotide binding (GO:0032553) |
| 1.84 | 62 | 0.00117 | protein-containing complex binding (GO:0044877) |
| 1.81 | 58 | 2.94E-03 | enzyme regulator activity (GO:0030234) |
| 1.78 | 104 | 6.38E-06 | carbohydrate derivative binding (GO:0097367) |
| 1.76 | 98 | 2.05E-05 | nucleotide binding (GO:0000166) |
| 1.75 | 98 | 1.96E-05 | nucleoside phosphate binding (GO:1901265) |
| 1.64 | 87 | 1.29E-03 | enzyme binding (GO:0019899) |
| 1.62 | 100 | 4.22E-04 | anion binding (GO:0043168) |
| 1.61 | 104 | 2.46E-04 | small molecule binding (GO:0036094) |
| 1.49 | 153 | 5.15E-05 | nucleic acid binding (GO:0003676) |
| 1.46 | 223 | 3.07E-07 | heterocyclic compound binding (GO:1901363) |
| 1.45 | 225 | 2.42E-07 | organic cyclic compound binding (GO:0097159) |
| 1.27 | 198 | 1.24E-02 | ion binding (GO:0043167) |
| 1.17 | 432 | 7.60E-07 | protein binding (GO:0005515) |
| 0.43 | 18 | 0.0057 | molecular transducer activity (GO:0060089) |
| 0.43 | 18 | 5.56E-03 | signaling receptor activity (GO:0038023) |
| 0.36 | 13 | 2.75E-03 | transmembrane signaling receptor activity (GO:0004888) |
| 0.09 | 2 | 1.92E-05 | G protein-coupled receptor activity (GO:0004930) |

**Supplementary Table 8 – Enriched GO Biological Processes from recombinant CoCoA nuclear interactome**

| Fold Enrichment | Genes in list | FDR | Functional Category – Biological Process |
|---|---|---|---|
| 40.3 | 3 | 0.0489 | pre-replicative complex assembly involved in cell cycle DNA replication (GO:1902299) |
| 40.3 | 3 | 0.0478 | pre-replicative complex assembly involved in nuclear cell cycle DNA replication (GO:0006267) |
| 40.3 | 3 | 0.0467 | pre-replicative complex assembly (GO:0036388) |
| 17.1 | 4 | 0.0498 | positive regulation of mRNA splicing, via spliceosome (GO:0048026) |
| 9.17 | 8 | 0.00699 | DNA duplex unwinding (GO:0032508) |
| 8.55 | 8 | 0.0089 | DNA geometric change (GO:0032392) |
| 7.3 | 8 | 0.0191 | regulation of mRNA splicing, via spliceosome (GO:0048024) |
| 6.27 | 10 | 0.00944 | regulation of RNA splicing (GO:0043484) |
| 3.88 | 12 | 0.0398 | regulation of mRNA metabolic process (GO:1903311) |
| 3.15 | 16 | 0.034 | DNA repair (GO:0006281) |
| 2.84 | 17 | 0.0491 | mRNA metabolic process (GO:0016071) |
| 2.63 | 20 | 0.0415 | DNA metabolic process (GO:0006259) |
| 2.55 | 23 | 0.0295 | RNA processing (GO:0006396) |
| 2.37 | 24 | 0.049 | regulation of cellular component biogenesis (GO:0044087) |
| 2.25 | 48 | 0.00101 | nucleic acid metabolic process (GO:0090304) |
| 2.21 | 33 | 0.0182 | RNA metabolic process (GO:0016070) |
| 2.17 | 36 | 0.0159 | negative regulation of macromolecule biosynthetic process (GO:0010558) |
| 2.13 | 35 | 0.0205 | negative regulation of cellular macromolecule biosynthetic process (GO:2000113) |
| 2.11 | 37 | 0.0164 | negative regulation of biosynthetic process (GO:0009890) |
| 2.09 | 36 | 0.0201 | negative regulation of cellular biosynthetic process (GO:0031327) |
| 2.09 | 55 | 0.00169 | nucleobase-containing compound metabolic process (GO:0006139) |
| 1.99 | 57 | 0.0012 | cellular aromatic compound metabolic process (GO:0006725) |
| 1.98 | 42 | 0.0165 | gene expression (GO:0010467) |
| 1.97 | 55 | 0.00232 | heterocycle metabolic process (GO:0046483) |

| | | | |
|---|---|---|---|
| 1.96 | 61 | 8.69E-04 | organic cyclic compound metabolic process (GO:1901360) |
| 1.86 | 63 | 0.00204 | cellular nitrogen compound metabolic process (GO:0034641) |
| 1.83 | 46 | 0.0341 | regulation of cellular component organization (GO:0051128) |
| 1.82 | 65 | 0.00189 | organelle organization (GO:0006996) |
| 1.82 | 57 | 0.00862 | negative regulation of metabolic process (GO:0009892) |
| 1.75 | 51 | 0.0312 | negative regulation of macromolecule metabolic process (GO:0010605) |
| 1.65 | 65 | 0.0199 | regulation of RNA metabolic process (GO:0051252) |
| 1.63 | 68 | 0.02 | regulation of nucleobase-containing compound metabolic process (GO:0019219) |
| 1.61 | 67 | 0.0273 | regulation of cellular macromolecule biosynthetic process (GO:2000112) |
| 1.6 | 67 | 0.0331 | regulation of macromolecule biosynthetic process (GO:0010556) |
| 1.59 | 69 | 0.0327 | regulation of cellular biosynthetic process (GO:0031326) |
| 1.58 | 92 | 0.00213 | cellular component organization or biogenesis (GO:0071840) |
| 1.56 | 88 | 0.00597 | cellular component organization (GO:0016043) |
| 1.5 | 76 | 0.0424 | regulation of gene expression (GO:0010468) |

**Supplementary Table 9 – Enriched GO Molecular Functions from recombinant CoCoA nuclear interactome**

| Fold Enrichment | Genes in list | FDR | Functional Category – Biological Process |
|---|---|---|---|
| 9.02 | 7 | 0.0035 | DNA helicase activity (GO:0003678) |
| 8.87 | 10 | 1.14E-04 | ATPase, acting on DNA (GO:0008094) |
| 6.55 | 11 | 4.08E-04 | helicase activity (GO:0004386) |
| 5.71 | 17 | 1.49E-05 | ATP hydrolysis activity (GO:0016887) |
| 5.29 | 13 | 4.30E-04 | catalytic activity, acting on DNA (GO:0140097) |
| 4.59 | 25 | 5.28E-07 | ATP-dependent activity (GO:0140657) |
| 4.3 | 14 | 0.00157 | mRNA binding (GO:0003729) |
| 3.48 | 22 | 1.61E-04 | catalytic activity, acting on a nucleic acid (GO:0140640) |
| 3.18 | 21 | 9.63E-04 | nucleoside-triphosphatase activity (GO:0017111) |
| 2.9 | 21 | 0.00322 | pyrophosphatase activity (GO:0016462) |
| 2.88 | 21 | 0.00338 | hydrolase activity, acting on acid anhydrides, in phosphorus-containing anhydrides (GO:0016818) |
| 2.88 | 21 | 0.00326 | hydrolase activity, acting on acid anhydrides (GO:0016817) |
| 2.79 | 50 | 6.17E-08 | RNA binding (GO:0003723) |
| 2.58 | 41 | 1.93E-05 | ATP binding (GO:0005524) |
| 2.47 | 41 | 4.09E-05 | adenyl ribonucleotide binding (GO:0032559) |
| 2.45 | 41 | 4.43E-05 | adenyl nucleotide binding (GO:0030554) |
| 2.35 | 46 | 3.13E-05 | purine ribonucleoside triphosphate binding (GO:0035639) |
| 2.29 | 47 | 3.42E-05 | ribonucleotide binding (GO:0032553) |
| 2.26 | 46 | 7.87E-05 | purine ribonucleotide binding (GO:0032555) |
| 2.25 | 52 | 1.73E-05 | nucleotide binding (GO:0000166) |
| 2.25 | 52 | 1.55E-05 | nucleoside phosphate binding (GO:1901265) |
| 2.24 | 46 | 8.08E-05 | purine nucleotide binding (GO:0017076) |
| 2.15 | 55 | 2.07E-05 | anion binding (GO:0043168) |
| 2.06 | 55 | 7.10E-05 | small molecule binding (GO:0036094) |
| 2 | 85 | 5.46E-08 | nucleic acid binding (GO:0003676) |
| 1.94 | 47 | 0.00209 | carbohydrate derivative binding (GO:0097367) |
| 1.79 | 113 | 1.12E-08 | heterocyclic compound binding (GO:1901363) |

| | | | |
|---|---|---|---|
| 1.76 | 113 | 1.86E-08 | organic cyclic compound binding (GO:0097159) |
| 1.21 | 185 | 1.48E-04 | protein binding (GO:0005515) |

**Supplementary Table 10 – Enriched GO Biological processes from endogenous NDP52 interactome in non-treated HeLa cells**

| Fold Enrichment | Genes in list | FDR | Functional Category – Biological Process |
|---|---|---|---|
| 2.77 | 24 | 0.00874 | ncRNA metabolic process (GO:0034660) |
| 2.68 | 21 | 0.0354 | ribonucleoprotein complex biogenesis (GO:0022613) |
| 2.61 | 25 | 0.0127 | supramolecular fiber organization (GO:0097435) |
| 2.3 | 36 | 0.00466 | RNA processing (GO:0006396) |
| 2 | 52 | 0.00234 | RNA metabolic process (GO:0016070) |
| 1.97 | 73 | 7.16E-5 | nucleic acid metabolic process (GO:0090304) |
| 1.95 | 45 | 0.0179 | intracellular transport (GO:0046907) |
| 1.93 | 49 | 0.0117 | cellular protein localization (GO:0034613) |
| 1.91 | 49 | 0.0122 | cellular macromolecule localization (GO:0070727) |
| 1.91 | 45 | 0.0223 | establishment of protein localization (GO:0045184) |
| 1.87 | 69 | 5.52E-4 | gene expression (GO:0010467) |
| 1.84 | 84 | 8.18E-5 | nucleobase-containing compound metabolic process (GO:0006139) |
| 1.82 | 64 | 0.00324 | protein localization (GO:0008104) |
| 1.81 | 88 | 9.04E-5 | heterocycle metabolic process (GO:0046483) |
| 1.8 | 77 | 5.44E-4 | macromolecule localization (GO:0033036) |
| 1.79 | 75 | 9.15E-4 | cellular localization (GO:0051641) |
| 1.77 | 88 | 1.38E-4 | cellular aromatic compound metabolic process (GO:0006725) |
| 1.71 | 101 | 8.49E-5 | cellular nitrogen compound metabolic process (GO:0034641) |
| 1.7 | 92 | 3.51E-4 | organic cyclic compound metabolic process (GO:1901360) |
| 1.62 | 71 | 0.0248 | regulation of cellular component organization (GO:0051128) |
| 1.59 | 73 | 0.0459 | cellular component biogenesis (GO:0044085) |
| 1.56 | 158 | 1.03E-5 | cellular component organization or biogenesis (GO:0071840) |
| 1.54 | 151 | 3.12E-5 | cellular component organization (GO:0016043) |
| 1.51 | 160 | 2.61E-5 | macromolecule metabolic process (GO:0043170) |

**Supplementary Table 11 – Enriched GO Biological processes from endogenous NDP52 interactome in HeLa cells following α-amanitin treatment**

| Fold Enrichment | Genes in list | FDR | Functional Category – Biological Process |
|---|---|---|---|
| 7.49 | 7 | 0.0339 | mitochondrion localization (GO:0051646) |
| 5 | 10 | 0.0279 | regulation of DNA replication (GO:0006275) |
| 4.39 | 11 | 0.0339 | signal transduction in response to DNA damage (GO:0042770) |
| 3.65 | 25 | 1.66E-4 | regulation of cell cycle phase transition (GO:1901987) |
| 3.28 | 17 | 0.0205 | regulation of mitotic cell cycle phase transition (GO:1901990) |
| 2.86 | 32 | 4.76E-4 | mitotic cell cycle (GO:0000278) |
| 2.73 | 26 | 0.00629 | mitotic cell cycle process (GO:1903047) |
| 2.73 | 32 | 8.54E-4 | regulation of cell cycle process (GO:0010564) |
| 2.72 | 22 | 0.0214 | regulation of mitotic cell cycle (GO:0007346) |
| 2.7 | 24 | 0.0142 | cell junction organization (GO:0034330) |
| 2.53 | 34 | 0.00241 | cellular response to DNA damage stimulus (GO:0006974) |
| 2.52 | 30 | 0.00806 | protein localization to organelle (GO:0033365) |
| 2.35 | 44 | 5.72E-4 | regulation of cell cycle (GO:0051726) |
| 2.31 | 44 | 7.19E-4 | chromosome organization (GO:0051276) |
| 2.19 | 33 | 0.0276 | cell cycle process (GO:0022402) |
| 2.17 | 49 | 8.66E-4 | cell cycle (GO:0007049) |
| 1.99 | 50 | 0.00578 | protein-containing complex subunit organization (GO:0043933) |
| 1.93 | 43 | 0.029 | protein-containing complex assembly (GO:0065003) |
| 1.89 | 49 | 0.0162 | cellular macromolecule localization (GO:0070727) |
| 1.88 | 42 | 0.0468 | negative regulation of signal transduction (GO:0009968) |
| 1.87 | 48 | 0.0282 | cellular protein localization (GO:0034613) |
| 1.82 | 114 | 5.61E-7 | organelle organization (GO:0006996) |
| 1.79 | 49 | 0.0389 | intracellular signal transduction (GO:0035556) |
| 1.79 | 67 | 0.00423 | nucleic acid metabolic process (GO:0090304) |
| 1.76 | 50 | 0.0479 | cellular response to stress (GO:0033554) |
| 1.74 | 179 | 2.7E-12 | cellular component organization or biogenesis (GO:0071840) |

| | | | |
|---|---|---|---|
| 1.73 | 73 | 0.00542 | cellular localization (GO:0051641) |
| 1.72 | 80 | 0.00233 | cellular component biogenesis (GO:0044085) |
| 1.72 | 170 | 7.99E-11 | cellular component organization (GO:0016043) |
| 1.68 | 71 | 0.0129 | cellular component assembly (GO:0022607) |
| 1.6 | 74 | 0.028 | nucleobase-containing compound metabolic process (GO:0006139) |
